# Reverse Proteolysis Uncovers a Hidden Dimension of the Peptidome

**DOI:** 10.1101/2025.09.02.673809

**Authors:** S. Yasin Tabatabaei Dakhili, Preety Panwar, Olivier Hinse, Eliot Mar, Jason Rogalski, Leonard J. Foster, Dieter Brömme

**Affiliations:** Faculty of Dentistry, Department of Oral Biological & Medical Sciences, The University of British Columbia; Vancouver, Canada; Faculty of Medicine, Department Biochemistry & Molecular Biology, Michael Smith Laboratories, Life Sciences Institute, The University of British Columbia; Vancouver, Canada

## Abstract

Proteases are conventionally regarded as degradative enzymes, yet their catalytic machinery also permits peptide bond formation through reverse proteolysis, a process that remains poorly characterized. Here, we show that lysosomal cysteine cathepsins catalyze iterative cycles of hydrolysis and ligation to generate multi-generational fusion peptides, including hybrids derived from host-viral protein substrates. Quantitative analysis demonstrates that peptide ligation can account for up to 4.5% of proteolytic turnover. This activity is strongly influenced by pH, substrate sequence, and post-translational modification, with citrullination and neutral pH favoring fusion peptide formation and the accumulation of more stable higher-order products. Using full-length protein substrates, we provide direct evidence that cathepsins can generate a hybrid insulin peptide previously identified as a Type 1 Diabetes (TID) autoantigen in patients. Moreover, several identified fusion peptides show effective binding to TID-associated HLA class II molecules. To examine whether ligation products can be captured under cellular conditions, we developed a click-based targeted transpeptide retrieval and purification strategy (CT-TRAP), which enabled detection of probe-derived cis/transpeptides in cell-based systems under controlled conditions. These findings establish reverse proteolysis by cysteine cathepsins as a quantifiable enzymatic pathway for generating non-genomically templated peptides, revealing an unrecognized dimension of lysosomal protease activity and peptide diversification.

## Introduction

Proteases are conventionally regarded as degradative enzymes that break down proteins into smaller fragments. However, the same catalytic machinery possesses the latent capacity to synthesize new peptide bonds by joining peptide fragments - a process known as reverse proteolysis or peptide ligation. Although this reactivity is a well-established tool in synthetic peptide chemistry ^1^, it has received little attention in the context of cellular peptide repertoires, leaving protease-catalyzed fusion or hybrid peptides (cis/transpeptides) as an underexplored component of the peptidome. Here, cis-peptides refer to ligation products formed from two peptide fragments derived from the same parent polypeptide, whereas trans-peptides arise from protease-catalyzed ligation between fragments originating from distinct protein substrates.

Recent studies have begun to shift this paradigm, showing that proteasomes can generate cis/transpeptides presented by Major Histocompatibility Complex (MHC) Class I molecules ^2–4^. Similarly, MHC Class II molecules, such as the Type 1 Diabetes (T1D)-implicated human leukocyte antigen (HLA)-DQ8, have been shown to present cis/transpeptides specifically recognized by disease-associated CD4^+^ T cells ^5^. Despite these advances, biochemical validation remains limited, fueling skepticism that reported cis/transpeptides may represent mass-spectrometric artifacts, mRNA splicing products, or genomic mutations ^6^. Moreover, the fundamental characteristics of reverse proteolysis, including enzyme-level substrate specificity, catalytic efficiency, the generation of higher-order cis/transpeptides, and modulation by factors like pH and post-translational modifications (PTMs), are poorly defined. A comprehensive understanding of the full catalytic potential of proteases is therefore urgently needed.

We hypothesize that cysteine cathepsins are well suited to catalyze cis/transpeptidation because their catalytic mechanism proceeds through a thioacyl enzyme intermediate, which can in principle be resolved by aminolysis. In addition, their high abundance in endolysosomal compartments, broad substrate specificity, and activity across a wide pH range ^7^, create conditions that can favor productive interactions with diverse peptide nucleophiles, while their central role in MHC class II associated antigen processing provides a biological context in which ligation products formed in this compartment could be functionally relevant ^8^. During ongoing proteolytic processing, cycles of hydrolysis and ligation mediated by these proteases could generate diverse repertoires of cis/transpeptides from distinct protein fragments. Such products may include hybrids combining pathogen- and host-derived sequences, providing a biochemical route for the emergence of a non-genomically templated peptide diversity from multiple protein sources during proteolytic processing. These hybrid peptides could increase the spectrum of sequences available for peptide–MHC interaction and therefore warrant consideration in the context of antigen processing mechanisms, where epidemiological links to microbial exposure have been reported, but the underlying biochemical mechanisms remain incompletely understood ^9–11^.

To address these challenges, we employed three complementary approaches. First, we conducted *in vitro* enzyme assays, progressing from small synthetic tagged peptide–peptide interactions to peptide–protein, and finally protein–protein interactions, to quantify the catalytic capacity of cysteine cathepsins and elucidate the effect of pH and PTMs. Second, we aimed to identify the specific proteases responsible for generating the clinically observed MHC Class II-associated transpeptides, such as those reported in T1D patients ^5^. Third, to address methodological limitations in detecting transpeptidation under cellular conditions, we developed a targeted chemical enrichment strategy, Click-based Targeted Transpeptide Retrieval and Purification (CT-TRAP), and applied it to a murine macrophage cell line to capture probe-derived cis/transpeptides.

Together, our findings demonstrate that lysosomal cysteine cathepsins can function iteratively to produce multi-generational cis/transpeptides, including pathogenic host-viral fusion products, and PTMs like citrullination and alterations in lysosomal pH can enhance this process. Considering the central role of cysteine cathepsins in MHC class II-dependent antigen processing ^8^, their potential in reverse proteolysis may widen our understanding of mechanisms that could influence antigen diversity. By expanding the catalytic repertoire of proteases beyond simple degradation, our work reframes cathepsins as dual-function molecular architects and establishes reverse proteolysis as a previously undervalued generator of antigenic diversity.

## Results

### Identification and quantification of multi-generational cis/transpeptides using a tagged synthetic peptide substrate assay

Although cathepsin-mediated transpeptidation has been reported ^12^, its quantitative competition with peptide bond hydrolysis remains undefined. It is also unclear whether cis- and transpeptide products are terminal species or can undergo further hydrolysis and re-ligation to generate higher-order cis/transpeptides. To address these questions, we used human cathepsin S (hCatS), a central protease in MHC class II antigen processing ^13–16^ with previously shown cis/transpeptidation activity ^17^.

We first investigated the role of free *N*-terminal peptides as nucleophiles in hCatS-catalyzed proteolysis at pH 6.5, a condition reflective of the early endolysosomes in antigen-presenting cells (APCs) ^18^. Using a Förster resonance energy transfer (FRET)-based substrate, Ac-`AE`(Edans)`FRTTSQGGK`-Dabcyl, we observed that the addition of a non-fluorescent competitor peptide (NH_2_-`TTSQGGK`-Biotin) increased the fluorescent readout by 1.6-fold (Fig. 1A). This enhancement was attributed not to increased hydrolysis but to the competitor peptide preventing the reverse proteolyis reaction that would otherwise quench fluorescence by ligating the NH_2_-`TTSQGGK`-Dabcyl hydrolysis fragment back to the FRET donor. We further confirmed this by adding the quenched product itself (NH_2_-`TTSQGGK`-Dabcyl), which suppressed the overall fluorescent readout by ∼3-fold. Together, these observations indicate that the presence and concentration of free *N*-terminal peptides can measurably influence the balance between hydrolytic and cis/transpeptidation under these conditions.

**Fig. 1:**
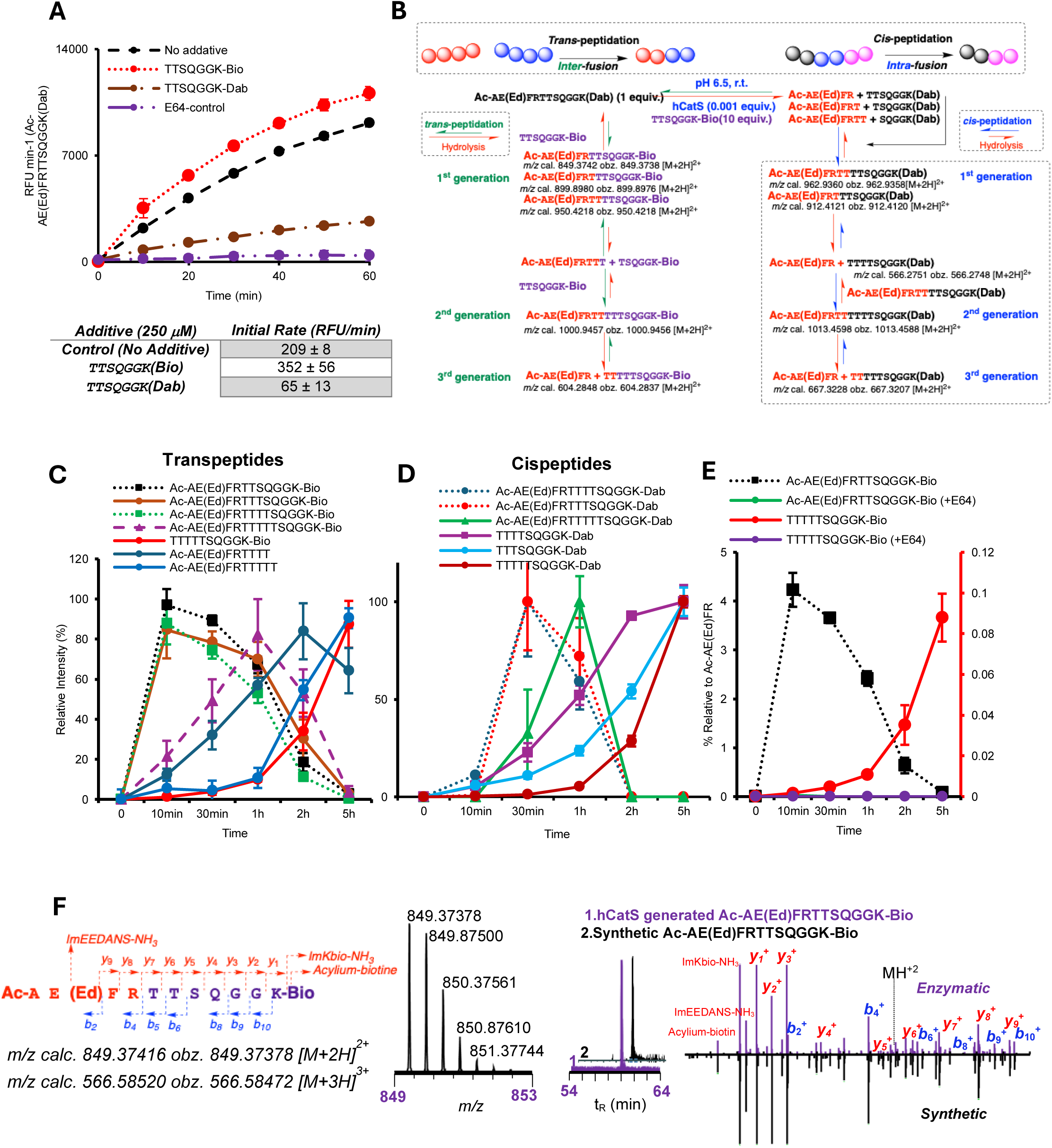
Cathepsin S-catalyzed multi-generational formation of cis/transpeptides using a model peptide substrate. (A) Time-course of hydrolysis of AC-`AE`(Ed)`FRTTSQGGK`(Dab) by hCatS in the presence or absence of NH_2_-`TTSQGGK`(Bio) or NH_2_-`TTSQGGK`(Dab) at pH 6.5. (B) Schematic of hCatS-generated transpeptides at pH 6.5 identified by LC-MS/MS. (C,D) Time-course analysis of transpeptides (C) and cispeptides (D) identified in (B). Peak areas were normalized to 100% at their maximal time points and plotted relative to that maximum. Comparisons are made within each peptide over time. (E) Percent formation of first-generation (Ac-`AE`(Ed)`FRTTSQGGK-`Bio) and third-generation (`TTTTTSQGGK`-Bio) transpeptides relative to first-generation hydrolysis product (Ac-`AE`(Ed)`FR`). (F) Comparison of retention time and fragment ion patterns between hCatS-generated Ac-`AE`(Ed)`FRTTSQGGK`(Bio) and its synthetic analogue (Bio: biotin; Dab: Dabcyl; Ed: Edans).

We then used LC-MS/MS to analyze the reaction products generated when hCatS was incubated with the FRET substrate and a 10-fold molar excess of NH_2_-`TTSQGGK`-Biotin. The peptides were strategically designed to yield distinct signature ions for Edans, Dabcyl, and Biotin (Fig. S1), considerably increasing the confidence of cis/transpeptide identification. Our analysis revealed a complex mixture of first- to third-generation cis/transpeptides within a 5 h reaction window (Fig. 1B, Fig. S2-S12). Kinetic analysis showed first-order transpeptides peaked rapidly within 10 min before undergoing gradual re-hydrolysis (Fig. 1C). In contrast, second- and third-order products emerged after 30 min and accumulated consistently over the 5 h period. Cispeptides followed a similar dynamic. First-order cispeptides (e.g., Ac-`AE`(Edans)`FRTTTTSQGGK`-Dabcyl) only became detectable after initial substrate hydrolysis, peaked at 30 min and declined after 2 h. Higher-order cispeptides (e.g., `TTTTTSQGGK`-Biotin) formed after 1 h and continued to be generated throughout the 5 h period (Fig. 1D). These time-course data establish that hCatS catalyzes a dynamic, iterative process governed by a complex and temporarily evolving interplay of hydrolysis and transpeptidation.

Since the immunological relevance of transpeptides depends not only on their identity but also on their abundance, we performed quantitative analysis of selected products using synthetic standards. This quantificiation revealed that a first-generation transpeptide (`AE`(Edans)`FR`-`TTSQGGK`(Biotin)) reached up to ∼4.5% of one of the main hydrolysis products (Ac-`AE`(Edans)`FR`-COOH) within 10 min (Fig. 1E). While these first-generation products were transient, a third-generation transpeptide (`TTTTTSQGGK`-Biotin) accumulated over 5 h, reaching 0.1% relative to Ac-`AE`(Edans)`FR`-COOH and notably exhibiting resistance to further hydrolysis by hCatS. The identity of these products was confirmed by matching retention times and fragmentation patterns with synthetic peptide standards (Fig. 1F, Fig. S13 and S14). Crucially, the addition of the pan-cathepsin inhibitor E-64 abolished all observed hydrolysis and transpeptidase activities, confirming that hCatS was solely responsible for catalyzing the reactions.

### Effect of nucleophilic peptide specificity and citrullination on cis/transpeptidation by hCatS

To investigate whether hCatS-mediated transpeptidation is a substrate-specific process, we used bovine serum albumin (BSA) as a model acceptor substrate and three distinct biotinylated nucleophilic donor peptides, each derived from a clinically relevant protein: NH_2_-`TTSQGGK`-Biotin from human platelet factor 4 (hPF4), NH_2_-`DLPGGGK`-Biotin from SARS-CoV-2 spike protein, and NH_2_-`NAVEGGK`-Biotin from islet amyloid polypeptide (IAPP). While SDS-PAGE analysis using Coomassie staining confirmed that the biotinylated peptides did not alter the overall pattern or quantity of BSA proteolytic fragments, we observed marked differences in the potency of each synthetic peptide to form transpeptides catalyzed by hCatS (Fig. 2A, B). Specificially, the hPF4-derived peptide was at least 4- to 5-fold more efficient in forming transpeptides than the other two peptides (Fig. 2C). This indicates that cathepsin-mediated transpeptidation is not solely governed by intrinsic nucleophilicity but depends on structural features of the nucleophilic peptide that enable productive engagement with the enzyme-bound acyl intermediate.

**Fig. 2:**
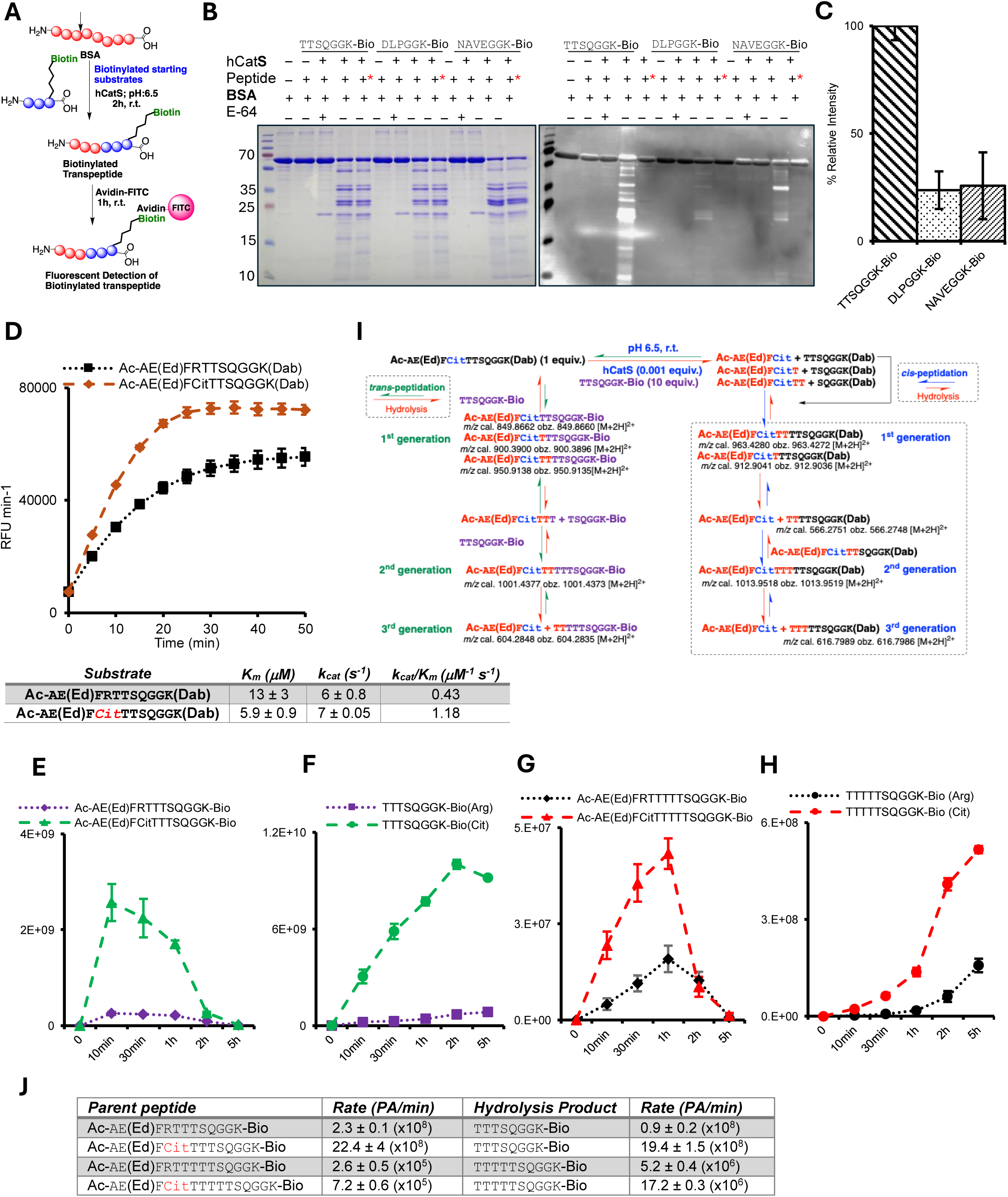
Nucleophilic peptide specificity and citrullination effects on the cis/transpeptidase activity of human cathepsin S. (A) Schematic of the method used to detect biotinylated cis/transpeptides generated by cathepsins. (B) SDS-PAGE of Coomassie-positive hydrolysis and cis/transpeptides and Avidin-FITC detected cis/transpeptides derived from bovine serum albumin (BSA) incubated with hCatS in the presence of three different biotinylated nucleophilic peptides at pH 6.5 for 2 h at RT, as well as in the presence or absence of the cathepsin inhibitor E-64. (C) Quantification of FITC detectable biotinylated cis/transpeptides in the presence of three different nucleophilic peptides. * Biotinylated peptide was added after BSA hydrolysis and CatS inhibition to exclude non-specific/non-covalent peptide binding. (D) Hydrolysis rates of Ac-`AE`(Ed)`FRTTSQGGK`(Dab) and Ac-`AE`(Ed)`FCitTTSQGGK`(Dab) by hCatS at pH 6.5 and their kinetic parameters in the accompanying table. (E) Proposed pathway for cis- and transpeptide generation from Ac-`AE`(Ed)F*Cit*TTSQGGK(Dab) by hCatS. (F,G) Time-course formation of transpeptides in independent reactions with citrullinated or non-citrullinated substrates. (H,I) Corresponding hydrolysis products from reactions in panels C and E. (J) Transpeptide formation rates in panels F-I, calculated as peak area divided by time. (Bio: biotin; Dab: Dabcyl; Ed: Edans).

Because citrullination (the post-translational modification of arginine residues) is strongly associated with autoimmunity such as in rheumatoid arthritis and known to alter protease recognition ^19^, we then investigated its effect on hCatS-mediated hydrolysis and transpeptidation. Kinetic analysis showed that a citrullinated FRET-based substrate was hydrolyzed with a 2.7-fold higher catalytic efficiency by hCatS compared to its non-citrullinated Arg analogue (Fig. 2D). This enhanced cleavage also led to a substantial increase in transpeptidation in the presence of the nucleophilic peptide, NH_2_-`TTSQGGK`-Biotin. Most citrullinated cis/transpeptides were generated at levels 2- to 10-fold higher than those from the non-citrullinated substrate (Fig. 2E-H, Fig. S21.A). Like the non-citrullinated version, the citrullinated substrate enabled the formation of multi-generational cis/transpeptides (Fig. 2I, Fig. S15-S20). Kinetic analysis of selected products further showed that the accelerated formation of citrullinated cis/transpeptides resulted in an increased abundance of stable cis/transpeptides (Fig. 2J, Fig. S21.B). These results establish that cathepsin-mediated transpeptidation is not a generic nucleophilic acyl-substitution reaction, but rather a selective, substrate-specific process. This preference likely arises from complex steric and electronic interactions between the amino acid sequence of the nucleophilic peptide and the enzyme’s prime subsite, and is also intrinsically dependent on the susceptibility of the original substrate for hydrolysis.

### *In vitro* generation of hybrid peptides corresponding to previously reported T1D-associated sequences

To investigate cysteine cathepsin-catalyzed transpeptidation using an autoimmune-relevant model system, we focused on the previously identified proinsulin-(IAPP) transpeptide (**`GQVELGGG`**`NAVEVLK`), which is presented by HLA-DQ8 and recognized by autoreactive CD4^+^ T cells isolated from patients with T1D ^5^. While cathepsins have been speculated to be involved in this process ^20,21^, and cathepsin L has shown the capacity to form related cis/transpeptides ^12^, the broader ability of various cathepsins to catalyze this and other unknown neoantigenic transpeptides remained poorly understood. Using recombinant human proinsulin in the presence of the biotinylated nucleophilic peptide derived from IAPP (NH2-`NAVEGGK`-Biotin) and hCatS at pH 6.5, we identified **`ELGGG`**`NAVEGGK`-Biotin (Fig. 3A) representing the identical fusion site between the C-peptide and IAPP as described by Tran et al. ^5^. Notably, we also observed a new product, **`QVELGGG`**`AVEGGK`-Biotin, with an ALC (Average Local Confidence) of 94% (Fig. 3B). Further analysis indicated that this product resulted from the high abundance of `AVEGGK`-Biotin (a hydrolysis byproduct of the original biotinylated peptide) and that the C-peptide sequence, `QVELGGG,` functions as a favorable acceptor substrate for hCatS. The formation of **`ELGGG`**`NAVEGGK`-Biotin, confirmed by a fragmentation pattern and retention time matching those of the synthetic standard (Fig. 3A; Fig. S23), was strongly pH dependent, with product yields more than twofold higher at endosomal pH 6.5 compared to lysosomal pH 4.5 (Fig. 3C).

**Fig. 3.**
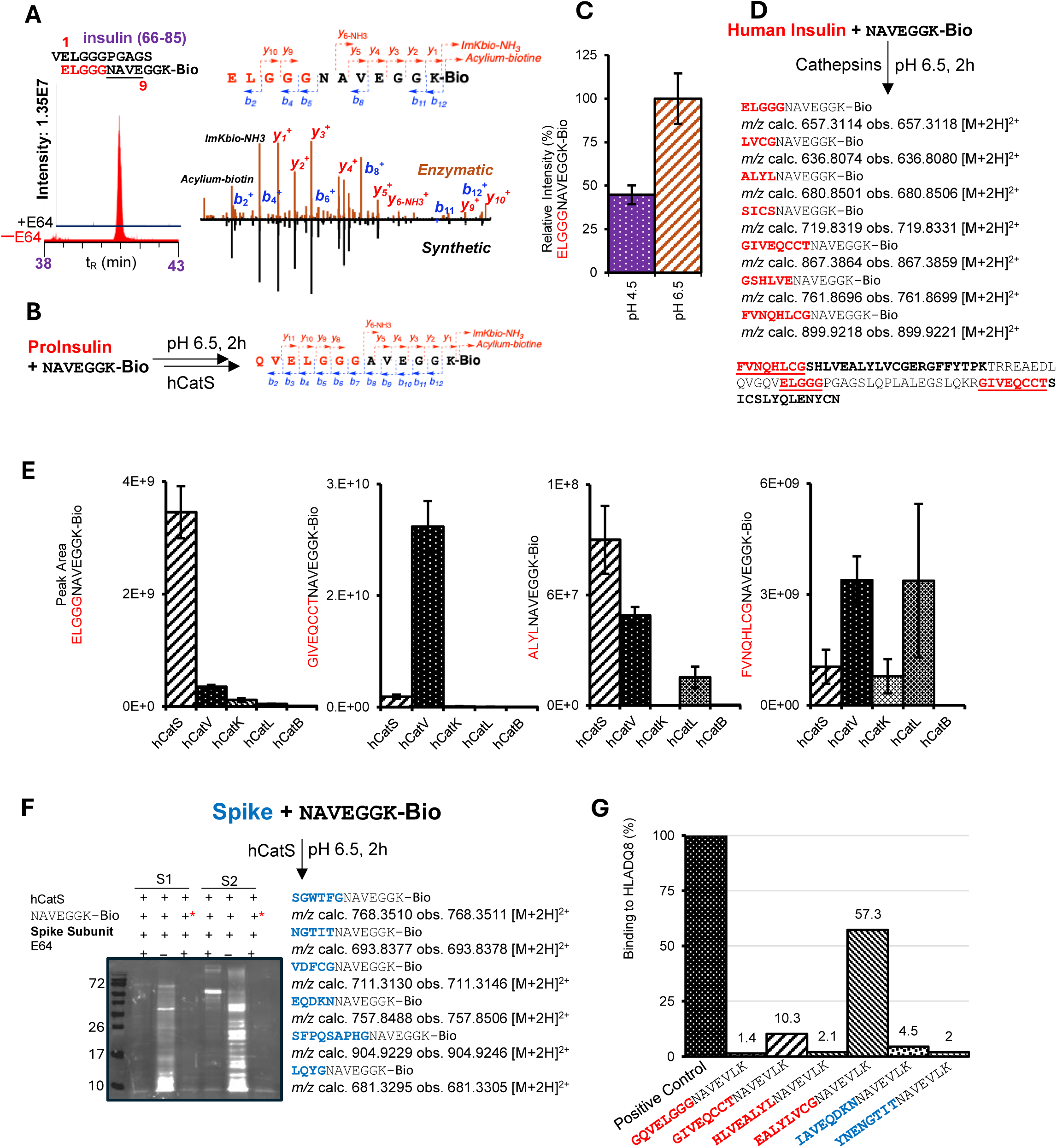
Generation of insulin-associated transpeptides by cysteine cathepsins. (A) Selected biotinylated transpeptides identified in a reaction mixture containing insulin and NH_2_-`NAVEGGK`-Biotin using LC-MS/MS. The insulin-based sequence in in red. (B) Chromatogram comparing retention times of ELGGGNAVEGGK-Bio generated in reactions with hCatS in the absence or presence of E-64 and fragment ion pattern of ELGGGNAVEGGK-Bio generated by hCatS and its synthetic analogue. (C) biotinylated transpeptides identified in a reaction mixture containing human proinsulin and NH_2_-`NAVEGGK`-Bio using LC-MS/MS. (D) Peak areas of ELGGGNAVEGGK-Bio generated by hCatS at pH 4.5 and pH 6.5. (E) Biotinylated peptides identified by LC-MS/MS from reactions containing cathepsins S,V, K, L, and B at pH 6.5. (F) Avidin-FITC blot of SARS-CoV-2 spike subunits (S1 and S2) incubated with hCatS and NH_2_-`NAVEGGK`-Biotin at pH 6.5 for 2 h at RT. (*) indicates control experiments where NH_2_-`NAVEGGK`-Biotin was added after a 2 h incubation of spike subunits with active hCatS, followed by E-64 inhibition. (G) HLA-DQ8 binding assay (performed by ProImmune Ltd.) compares the binding capacity of selected transpeptides shown in panel D and F (peptides were elongated *C*- and *N*-terminally to reach a length of 15 amino acids). (Bio: biotin).

We also observed this C-peptide-based fusion peptide (**`ELGGG`**`NAVEGGK`-Biotin) using mature insulin indicating that small contaminations with the C-peptide (Fig. S22) are sufficient to generate this known auto-antigen (Fig. 3A and Fig. S23). In addition, LC-MS/MS analysis confirmed additional multiple *trans*peptides (Fig. 3D), such as **`GIVEQCCT`**`NAVEGGK`-Biotin and **`ALYL`**`NAVEGGK`-Biotin (Fig. 3D and Fig. S23-S32).

Crucially, the presence of Glutamic acid (Glu) at positions 1 and 9 in two of these *trans*peptides (**`ELGGG`**`NAVEGGK`-Biotin and **`GIVEQCCT`**`NAVEGGK-`Biotin) corresponds to anchor residue patterns associated with strong HLA-DQ8 binding ^22^. The generation of fusion peptides with features consistent with known disease-associated epitopes, including a unique *trans*peptide with Gln at the P1 position (**`FVNQHLCG`**`NAVEGGK`-Biotin), demonstrates the catalytic capacity of hCatS to generate potential auto-antigens. This particular Gln-containing *trans*peptide is relevant because Gln can be selectively deaminated by tissue transglutaminase to Glu ^23^, which may further enhance its HLA-DQ8 binding compatibility. This finding resembles epitopes reported in other disease models, like celiac disease ^23^, but has not been previously identified as a T1D-associated epitope/neoepitope.

To determine whether *cis*/*trans*peptide generation is a general feature of cysteine cathepsins or unique to hCatS, we compared its transpeptidase activity with the ubiquitously expressed hCatB and hCatL and the tissue-specific hCatV and hCatK. This comparison is crucial, as hCatS is a prime candidate for driving autoimmune neoepitope formation in APCs, whereas other cathepsins might catalyze *cis*/*trans*peptides in distinct tissues or pathological settings. Human CatS produced the highest yield of the T1D *trans*peptide **`ELGGG`**`NAVEGGK`-Biotin. Furthermore, the immune tissue-dominant cathepsins (hCatS and hCatV) showed higher yields for most identified *trans*peptides compared to the ubiquitously expressed hCatL and hCatB (Fig. 3E). Human CatL demonstrated comparable potency for only one of the detectable *trans*peptides (**`FVNQHLCG`**`NAVEGGK`-Biotin). These findings suggest that variability in the tissue distribution of cysteine cathepsins, together with differences in their transpeptidation activity, may contribute to the generation of distinct fusion peptide repertoires in different tissue environments.

Given the epidemiological link between SARS-CoV-2 infection/vaccination and T1D ^24–26^, we investigated whether hCatS could generate *cis*/*trans*peptides between the human IAPP-derived nucleophilic peptide (NH_2_-`NAVEGGK`-Biotin) and the SARS-CoV-2 S1 and S2 subunits. We identified several biotinylated *trans*peptides, with the S2 subunit acting as a more efficient acceptor substrate than S1 (Fig. 3F, Fig. S33-S36). The formation of these products was confirmed to be enzyme-specific by its abolition upon addition of the cathepsin inhibitor E-64. Crucially, we discovered the host-viral *cis*/*trans*peptide **`EQDKN`**`NAVEGGK`-Biotin, which contains the signature Glu residues at positions 1 and 9 known to create super-agonists for HLA-DQ8 molecules ^22^. These results establish a plausible framework by which infection-derived peptide sequences could be enzymatically incorporated into fusion peptides, demonstrating that host- and viral-derived fragments can serve as substrates for cathepsin-mediated ligation under defined conditions.

### *In vitro* binding of cis/transpeptides to disease-relevant HLA molecules

Since stable peptide binding to HLA molecules is a prerequisite for formation of peptide HLA complexes capable of engaging T cell receptors, the binding affinity of selected trans peptides to disease associated HLA molecules was experimentally evaluated using a commercial HLA class II binding assay platform (ProImmune Inc.). Binding of six trans peptides, generated between insulin or spike derived fragments and `NAVEGGK-`Biotin, to the T1D associated HLA-DQ8 molecule (DQA1\**03:01-DQB1\**03:02) was evaluated using this platform. Three of these peptides, containing the critical Glu residues at positions 1 and 9, showed equal or higher binding affinity for HLA-DQ8 than the established T1D auto-antigen reference (5) (Fig. 3G). Notably, the insulin-derived 15-mer **`EALYLVCG`**`NAVEVLK` displayed a forty-one-fold greater affinity for HLA-DQ8 than the known reference peptide. Furthermore, two spike-derived transpeptides, **`IAVEQDKN`**`NAVELK` and **`YNENGTIT`**`NAVEVLK`, exhibited equal or moderately increased binding, despite the latter containing Glu residues at positions 1 and 10 (Fig. 3G). These experimental data demonstrate that cathepsin-mediated transpeptidation can generate fusion peptides with enhanced binding affinity to the disease-associated HLA class II molecule HLA-DQ8.

### Exploring cis/transpeptidation as a potential biochemical source of peptides that resemble disease-associated epitopes in autoimmune contexts with unresolved etiology

To broaden the scope of cathepsin-generated cis/transpeptides as a potential source of peptides resembling disease-associated epitopes, we investigated whether hCatS could generate fusion peptides derived from hPF4, a protein implicated in thrombotic autoimmune syndromes such as, heparin-induced (HIT) and vaccine-induced immune thrombotic thrombocytopenia (VITT).

To investigate this, we incubated full-length hPF4 in the presence of hCatS at pH 6.5 and a traceable nucleophilic peptide (NH_2_-**`TTSQ`**`GGK-`Biotin) whose sequence is located within a major PF4 epitope known to trigger thrombotic autoimmunity (HIT and VITT; **<u>`LQCLC`</u>`V`K**↓**TTSQVRP<u>RH</u>**) ^27,28^. Using LC-MS/MS, we identified a diverse array of cis/transpeptides (Fig. 4A; Fig. S37-S54), including a third-generation cispeptide, **`TTSTTTT`**`TTSQGGK`-Biotin. The presence of all intermediate fragments in the reaction mixture confirmed that cathepsin-mediated transpeptidation can generate a complex and diverse repertoire of potential antigens from a limited set of initial fragments (Fig. 4B). We also identified **`SLEVIK`**`TTSQGGK`-Biotin, a cispeptide containing part of a previously discovered VITT epitope ^29^. The formation of this peptide was completely abolished by the cathepsin inhibitor E-64, confirming again that the reaction was catalyzed by hCatS (Fig. 4C).

**Fig. 4.**
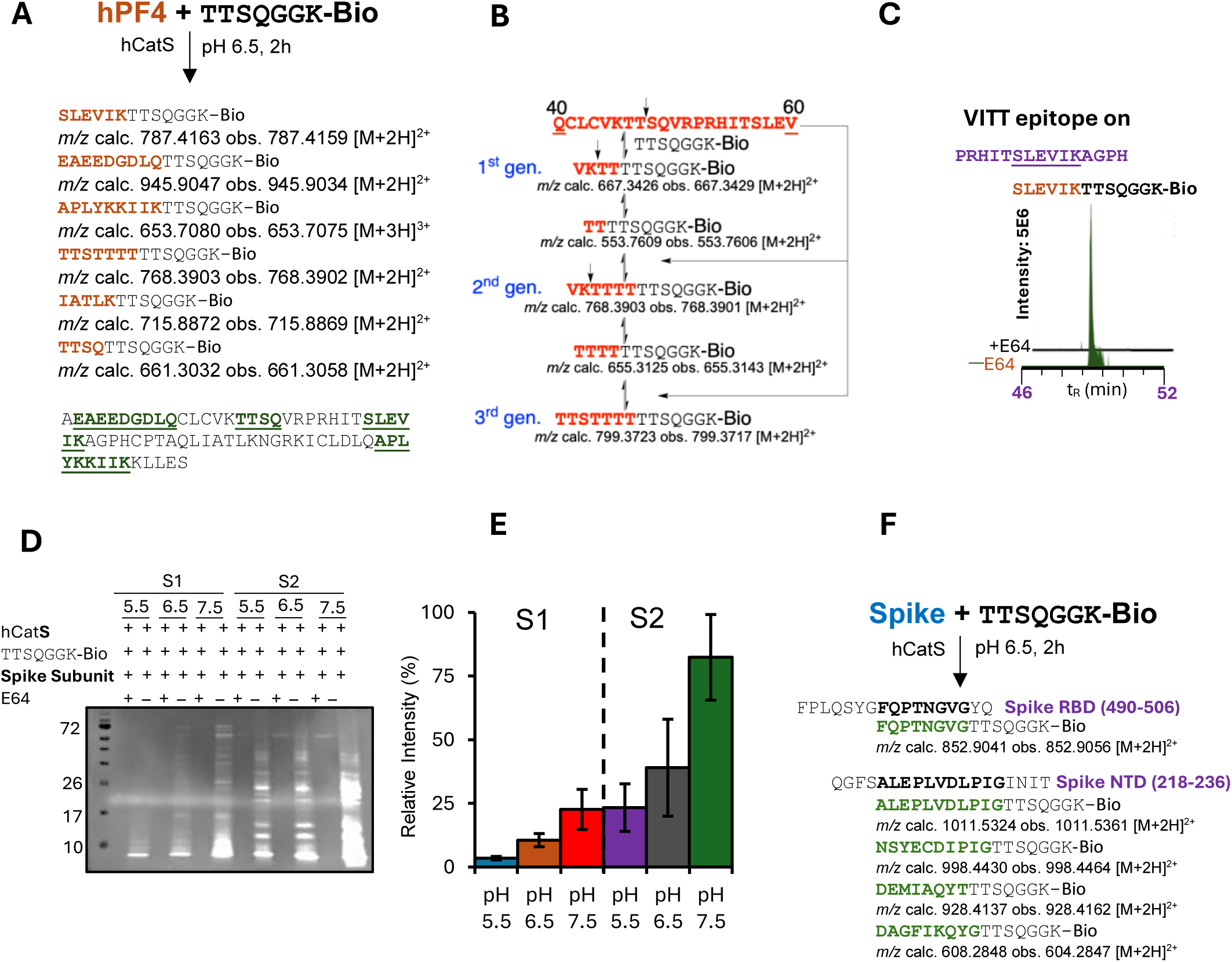
Generation of cis/transpeptides by hCatS in the presence of hPF4 or spike protein subunits and a hPF4 derived antigenic biotinylated nucleophilic peptide. (A) Selected biotinylated cis/transpeptides identified in reaction mixture containing hPF4, NH_2_-`TTSQGGK`-Bio and hCatS at pH 6.5 using LC-MS/MS and hPF4 amino acid sequence with identified acceptor peptide regions in bold. (B) Proposed mechanistic pathway for the formation of the identified third-generation transpeptide (**`TTSTTTT`**`TTSQGGK`-Bio). (C) Ion chromatogram of **`SLEVIK`**`TTSQGGK`-Bio generated by hCatS in the presence or absence of E-64. (D) Avidin-FITC blot of the reaction mixtures of S1 and S2 subunits (S1 and S2) incubated with hCatS and NH_2_-`TTSQGGK`-Bio for 2 h at RT at different pH values. (E) Quantification of Avidin-FITC signals from (D). (F) Biotinylated transpeptides identified by LC-MS/MS from reaction mixtures containing S1 and S2 subunits, NH_2_-`TTSQGGK`-Biotin, and hCatS at pH 6.5 for 2 h. (Bio: biotin).

Next, we tested the formation of transpeptides between the hPF4-derived nucleophilic peptide and SARS-CoV-2 spike subunits S1 and S2. Our results revealed the formation of higher molecular weight biotinylated products, which were more prominent in the presence of the S2 subunit and increased for both subunits with rising pH (Fig. 4D, E; S55A). This pH dependence is consistent with our previous observations for T1D-associated transpeptides (Fig. 3C).

Most notably, we discovered cis/transpeptides derived from two spike protein regions previously reported to be strongly antigenic ^30^: the S1 receptor-binding domain (RBD) (**`FQPTNGVG`**`TTSQGGK`-Biotin) and the *N*-terminal domain (NTD) (**`ALEPLVDLPIG`**`TTSQGGK`-Biotin) (Fig. 4F; Fig. S56-S60). Furthermore, this hCatS-orchestrated transpeptidase activity was also observed with the full-length, glycosylated spike protein (Fig. S55B), demonstrating that this activity is not confined to *E.coli*-expressed, non-glycosylated spike subunits but extends to physiologically relevant viral proteins.

These findings show that hCatS can generate hybrid peptides bridging self and viral sequences, providing a biochemical framework for considering how such fusion peptides could arise in infection-associated inflammatory environments.

### Cis/transpeptidation between full-length proteins: Exploring complex cis/transpeptides derived from SARS-CoV-2 spike, insulin, and hPF4 proteins

To model physiologically relevant conditions more closely, we investigated the capability of hCatS to generate complex cis/transpeptides using full-length SARS-CoV-2 Spike subunits, insulin, and hPF4 proteins. Using LC-MS/MS and *de novo* peptide sequencing, we successfully identified a diverse array of first- and second-generation cis/transpeptides. The analysis identified several high-confidence cis/transpeptides formed from insulin and spike proteins, which were reproducibly detected in all three experimental replicates and absent in controls (Fig. 5A).

**Fig. 5.**
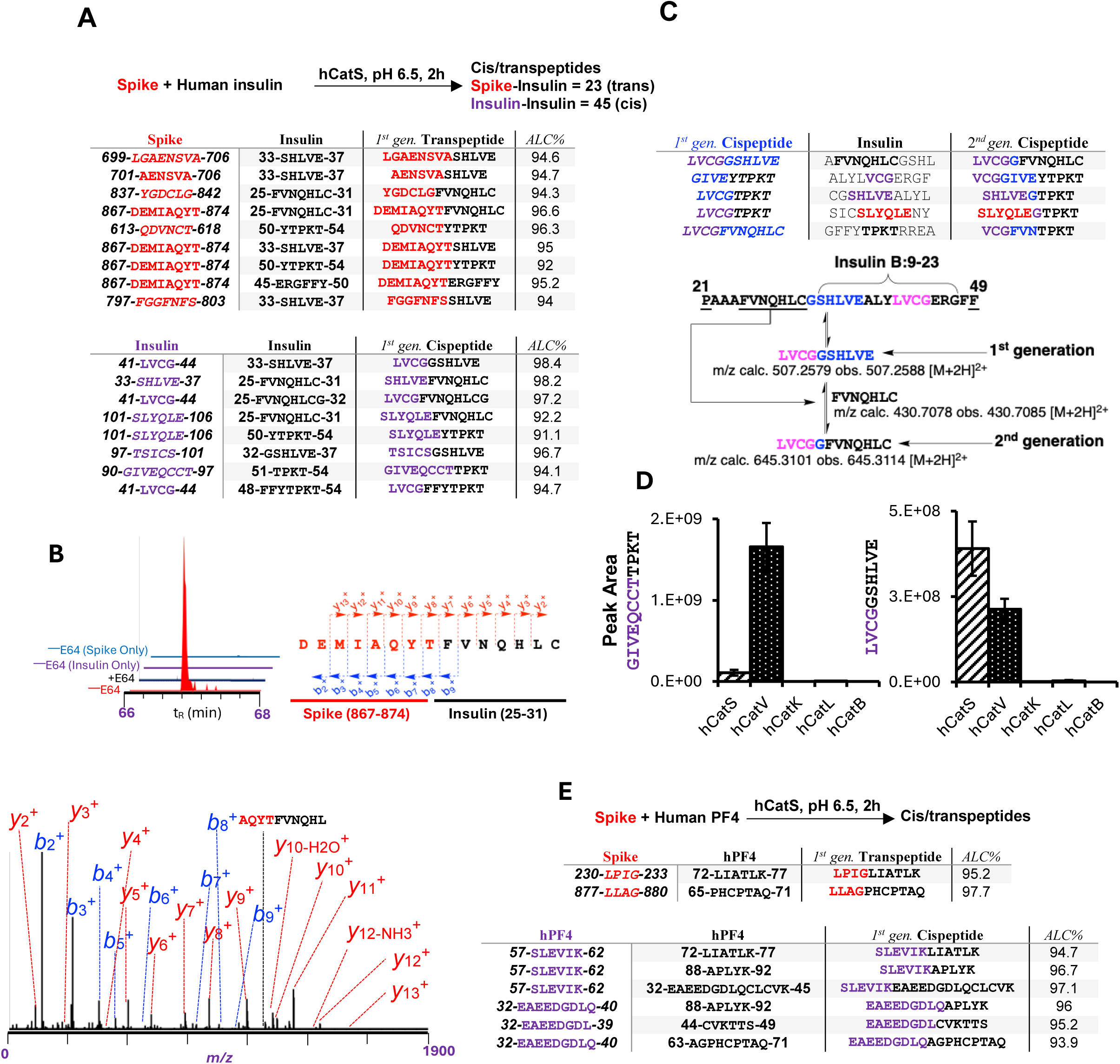
Cis/transpeptides generated by hcatS and other cysteine cathepsins from full-length insulin, hPF4, and S1/S2 subunit proteins. (A) Spike-Insulin trans- and insulin-insulin cispeptides derived from reaction mixtures containing SARS-CoV-2 spike protein, human insulin, and hCatS at pH 6.5 for 2 h at RT (N=3). Peptides were identified by LC–MS/MS *de novo* sequencing with ALC% ≥ 90, ensuring high-confidence assignments. (B) Ion chromatograms of a selected transpeptide generated by hCatS in the presence or absence of E-64. (C) Selected first- and second-generation cispeptides derived from insulin by hCatS and their proposed pathway of formation. (D) Comparing the formation of two selected cispeptides by different cysteine cathepsins. (E) First-generation of cis/transpeptides identified from a reaction mixture containing spike proteins, hPF4, and hCatS at pH 6.5 for 2 h at RT.

We identified and manually verified over 26 tandem mass spectra of novel transpeptides with a *de novo* confidence score (ALC score) exceeding 90% (Fig. 5A, Figs. S62–S72). Of particular interest was the identification of the spike-insulin transpeptide, **`DEMIAQYT`**`FVNQHLC` (Fig. 5B; Fig. S73), a first-generation product supported by a high ALC score of 96.6%. This sequence represents a fusion between SARS-CoV-2 spike and human insulin fragments where several residue positions (P1, P4, P6, and P9) correspond to sites previously reported to contribute to superagonist binding to HLA-DQ8^31^. Furthermore, we identified the first examples of second-generation cispeptides such as **`LVCG`***<u>`G`</u>*`FVNQHLC` (Fig. 5C, Fig. S74) and established the presence of all necessary precursor peptides in the reaction mixture.

To confirm that hybrid peptide generation from full-length proteins is also enzyme-specific, we compared the efficacy of hCatS, hCatV, hCatK, hCatL, and hCatB in forming two first-generation insulin-associated cispeptides (**`GIVEQCCT`**`TPKT` and **`LVCG`**`GSHLVE`). Consistent with our earlier findings (Fig. 3D), hCatS and hCatV produced higher cispeptide yields than the other proteases tested, while hCatL, hCatK, and hCatB produced only negligible amounts of these cispeptides (Fig. 5D). These results strongly reinforce that the nature of cis/transpeptidation products is highly dependent on the specificity of individual cathepsins.

Finally, using full-length spike and hPF4 proteins, we identified cis/transpeptides derived from PF4 and PF4–spike fragments, including sequences overlapping with previously reported sites to be antigenic in thrombotic thrombocytopenia syndromes such as HIT and VITT. We identified eight reproducibly detected hPF4–hPF4 and hPF4-spike cis/transpeptides that were absent in controls (Fig. 5E, S75). Of special interest was a hPF4-hPF4 cis-peptide (**`SLEVIK`**`LIATLK`) that contains part of a previously discovered VITT epitope ^29^. This **`SLEVIK`** fragment was also observed to fuse with the hPF4-derived nucleophilic peptide in our previous experiment (Fig. 4A, B). The generation of antigenic hPF4 cispeptides may reveal at least two potential PF4 epitopes that could facilitate antibody interactions with with neighboring PF4 molecules in hPF4 tetrameric complexes described for VITT ^32^.

### Identification of cis/transpeptides in cells using CT-TRAP

To examine whether probe-involved ligation chemistry can be captured under cellular conditions, we focused on antigen-presenting cells using RAW 264.7 macrophages as a model system. To achieve this, we developed a chemical enrichment and detection strategy termed: Click-based Targeted Transpeptide Retrieval and Purification (CT-TRAP) (Fig. 6A). The CT-TRAP probe (NH_2_-TTSQGG**K****(N3)**YGRKKRRQRRRK-5-FAM) was designed with a modular architecture. It contains an *N*-terminal nucleophilic peptide bearing the `TTSQ`-motif, which serves as a competitive substrate for transpeptidation and represents the variable region of the probe that can be adapted to different sequence contexts. A Lys(N3) residue was strategically incorporated as a bioorthogonal click handle, enabling post-lysis copper-catalyzed azide-alkyne cycloaddition (CuAAC) derivatization. This reaction Introduces a Lys-conjugated 4-PEG-Biotin tag that generates a diagnostic fragment ion (m/z 270.1276) in tandem mass spectra (Fig. S76) and permits enrichment of tagged species using streptavidin magnetic beads. The probe also contains a fluorogenic 5-FAM moiety to monitor uptake and a TAT sequence (YGRKKRRQRRRK) to facilitate lysosomal delivery. Using fluorescence microscopy, lysosomal uptake of the probe in RAW 264.7 macrophages was observed at high efficiency (Fig. 6A; Fig. S77).

**Fig. 6.**
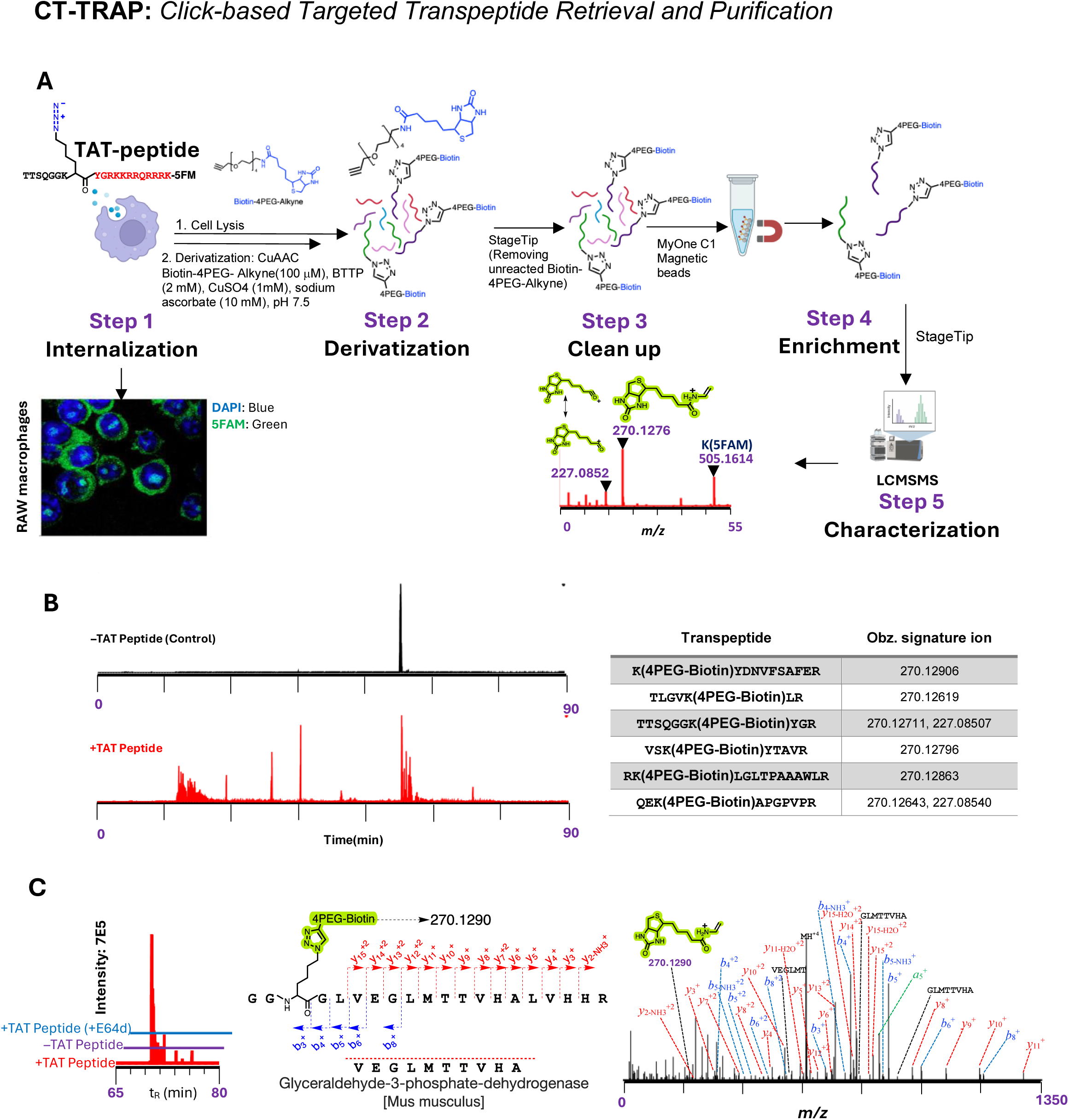
Identification of cis/transpeptides in live cells. (A) Scheme of the CT-TRAP method using a lysosomotropic TAT peptide containing CT-TRAP probe (NH_2_-`TTSQGGK`(N₃)-TAT-5FAM)) and its cellular uptake to identify cis/transpeptides in the RAW264.7 cell line. This method incorporated the inclusion of specific signature ions allowing the unequivocal identification of TAT-containing cis/transpeptides. (B) Peptides identified by LC-MS/MS containing the Lys-4PEG-Biotin signal at the *N*-terminal and in the middle of transpeptides as well as the processed but unligated CT-TRAP-probe, `TTSQGGK`(4-PEG-Biotin)`YGR`. None of the tagged peptides were seen in the absence of CT-TRAP probe. (C) Fusion product of the truncated probe with a sequence of endogenous GAPDH which was not observed in the presence of the E-64d.

The application of CT-TRAP followed by LC-MS/MS enabled the detection of probe-derived candidate species based on diagnostic signature fragment ions from the 4-PEG-Biotin tag (Fig. 6B; Fig. S78 and S79). We observed several peptides containing the Lys(4-PEG-Biotin) tag at different position of the fusion products indicating a complex processing of the original CT-TRAP probe at its *N*- and *C*-termini as well as subsequent hydrolysis steps of the fusion products. We also identified the partially processed probe-derived peptide TTSQGGK(4-PEG-Biotin)YGR, consistent with cellular uptake of the probe, partial lysosomal processing (YGR↓KK), retention of the Lys(N₃) handle for CuAAC derivatization, and the stability of the probe throughout enrichment and mass spectrometric workflows. This peptide was not detected in assays lacking the probe (Fig. 6B). In an additional experiment using E-64d as control for eliminating cysteine cathepsin activities, we identified in the absence of E-64d a `Lys`-(4-PEG-Biotin)-tagged peptide fused to a cellular protein, glyceraldehyde-3-phosphate dehydrogenase (GAPDH) indicating that cysteine cathepsins are responsible for generating this transpeptide within the endolysosomal compartment (Fig. 6C).

The primary objective of this experiment was to establish CT-TRAP as a chemical biology platform for detection and enrichment of probe-involved ligation products in cells rather than to comprehensively catalogue endogenous cis/transpeptides. These data demonstrate that ligation chemistry involving the synthetic probe can be captured under cellular lysosomal conditions, while the extent to which analogous ligation events occur between endogenous proteins remains to be determined.

## Discussion

This study resolves a long-standing knowledge gap in reverse proteolysis by demonstrating several mechanistic aspects of how cysteine cathepsins are capable of catalyzing the formation of cis/transpeptides. The observed ligation reactions occur through the intrinsic acyl-enzyme chemistry of these proteases and can compete measurably with hydrolysis under defined experimental conditions.

The fact that transpeptides, initially identified using small biotinylated peptides (e.g., **`SLEVIK`**`TTSQGGK`-Biotin), were also detected using full-length proteins (e.g., cispeptide **`SLEVIK`**`LIATLK`), demonstrated that tagged peptides can serve as effective tools for cis/transpeptide discovery under controlled experimental conditions. The incorporation of tags like biotin markedly increases confidence in peptide identification by LC-MS/MS, allowing for the unambiguous detection of rare cis/transpeptides in complex mixtures. This strategy enables high-confidence discovery under simplified conditions while capturing ligation events that can also be reproduced using full-length protein substrates.

Using this experimental approach, we showed that hCatS can catalyze ligation at the C-insulin peptide and an IAPP junction (`ELGGG-NAVE,` generating a hybrid insulin peptide that matches the sequence of a hybrid insulin peptide (HIP) shown to be recognized by CD4^+^ T cells from T1D patients ^5^. Our data show that hCatS is capable of generating a peptide sequence identical to a previously described HIP under defined *in vitro* conditions. In addition, we also identified several yet unknown transpeptides with high affinity binding to HLA-DQ8. Some of these novel transpeptides contain fragments from the SARS-CoV-2 spike protein and human insulin, highlighting the enzymatic capacity of hCatS to generate diverse fusion sequences. While earlier work focused largely on T1D models, our findings regarding the formation of hPF4-based cis/transpeptides suggest that this mechanism could also warrant consideration in other autoimmune diseases. Furthermore, our findings suggest that cathepsin-mediated reverse proteolysis may represent a biochemical mechanism capable of contributing to peptide diversity in different biological and disease context.

One criticism regarding the relevance of reverse proteolysis has been the assumption that it may only form negligible amounts of cis/transpeptides ^6^. However, in our model reaction using small peptides, one hybrid peptide transiently accounted for up to 4.5% of the major hydrolysis product ^5^. This yield is notable in the context of enzyme-catalyzed reactions and demonstrates that peptide ligation can compete measurably with hydrolysis under defined conditions. Although direct quantitative comparison to cellular systems is not possible, peptide–MHC biology indicates that low-abundance peptide species can be biologically relevant, suggesting that cis/transpeptides should be considered when evaluating how enzymatic processes influence peptide diversity ^34,35^.

Cathepsin-mediated transpeptidase activity is pH-dependent, with activity increasing when the pH value shifts from 4.5 to 7.5. This is particularly relevant because lysosomal dysfunction, often accompanied by alkalinization, is a hallmark of several autoimmune diseases ^36^. Since transpeptidase activity is optimal at neutral pH, the extracellular generation of larger fusion protein fragments by specific cathepsins appears plausible, which may increase the diversity of peptide species generated under extracellular conditions. The detection of high molecular fusion products identified in our biotin-avidin assays supports this possibility (Fig. 3F, 4D). In contrast, the diverse cathepsin content in the endolysosomal compartment may favour linear, short antigenic peptides suitable for MHC class II presentation.

Also of high relevance is our discovery that post-translational modifications, such as citrullination, can dramatically increase the efficiency of these transpeptidation reactions. This finding adds a new biochemical dimension to how citrullination may influence antigenic peptide diversity, suggesting that beyond altering individual peptide sequences, this PTM can enhance the formation of cis/trans fusion peptides with potential relevance to autoimmune contexts such as rheumatoid arthritis.

The sequence of cis/transpeptides depends on the specificity of individual cathepsins. Since their expression profile differs between the central and peripheral immune system ^16,37,38^, this may lead to the generation of different sets of cis/transpeptides. Differences in cathepsin expression across tissues may influence the repertoire of cis/transpeptides generated, raising the possibility that such peptides could contribute to antigenic diversity encountered by T cells in peripheral tissues, exemplified by the previously shown increased hCatS expression in peripheral DCs contributing to systemic lupus erythematosus ^39^. Future work should also consider that transpeptides may also interact with regulatory T cells and may have a significant effect on immune regulation.

Finally, application of the CT-TRAP approach enabled the capture of probe-derived cis/transpeptides under cellular conditions in antigen-presenting cells. These data indicate that cysteine protease–dependent fusion and hydrolysis events can occur in lysosome-targeted environments involving the synthetic probe. While these observations demonstrate the feasibility of capturing ligation chemistry in cells, determining the extent to which analogous processes between endogenous proteins influence antigen presentation or disease contexts will require further investigation.

In summary, our research establishes reverse proteolysis by lysosomal cysteine cathepsins as a fundamental mechanism for generating higher-order cis/transpeptides through iterative cycles of hydrolysis and ligation. This expands the traditional view of antigen processing beyond simple degradation, revealing an enzymatic pathway capable of generating fusion peptides from self- and non-self proteins that are compatible with current models of neoantigen diversity. This hypothesis suggests that self-derived peptides, for which T cells develop tolerance, do not remain constant throughout life and that PTMs may alter the peptide repertoire encountered by the immune system ^40^. In this context, we can consider cis/transpeptidation as a specific PTM. Cis/transpeptidation may represent an underexplored biochemical source of non-genomically templated peptides that could significantly expand the pool of antigens considered in autoimmune contexts. These cis/transpeptides represent non-genomically templated host-derived sequences that may be perceived as structurally distinct from canonical self peptides within antigen presentation pathways. By linking hCatS and other cathepsins to the enzymatic formation of a previously described disease-associated hybrid epitope and additional fusion peptides with strong HLA-binding compatibility, our study provides a biochemical framework for considering how such non-genomically templated sequences could arise during antigen processing. Furthermore, these findings raise the possibility that modulation of transpeptidation activity could influence the composition of the peptide repertoire generated during antigen processing. The challenge of future studies will be the identification of disease-associated T cells specifically recognizing these cis/transpeptide-based neoantigens and exploring methods to selectively block the formation of these hybrid peptides without interfering with the proteolytic function of cathepsins. Moreover, future studies should explore the potential relevance of reverse proteolysis across diverse biological contexts, including aging, chronic infection, and cancer, where changes in proteolytic environments may alter the cis/transpeptide diversity. It will also be important to define how cellular factors, such as lysosomal pH and post-translational modifications, regulate the balance between hydrolysis and transpeptidation under physiological conditions.

## Materials and Methods

### Materials

Native human PF4 protein (ABChem Inc., MA, USA; Cat. No. AB81754, Lot No. 1054529-1), rhProinsulin, recombinant human proinsulin >95% (*E.* coli-derived), Cat. No. 1336-PN, LOT# MJL0825011, R&D Systems, Minneapolis, MN, USA. Recombinant human insulin (Roche, Germany; Cat. No. 11376497001, yeast-derived), recombinant SARS-CoV-2 Spike protein S2 (RayBiotech Life Inc., GA, USA; Cat. No. 230-01103-250, Lot No. 02G10231), and recombinant SARS-CoV-2 Spike protein S1 (RayBiotech Life Inc.; Cat. No. 230-01101-250, Lot No. 09G21221) were used as protein substrates. Avidin–fluorescein conjugate (Invitrogen, Thermo Fisher Scientific, OR, USA; Ref. A821, Lot No. 2652967 and Lot:2831364) was employed for biotin detection. Mass spectrometry grade Trypsin-ultra™ was obtained from New England Biolabs (Cat. No. P8101S, Lot No. 10189739). The biotinylated peptides NH_2_-`TTSQGGK`-Biotin, NH_2_-`NAVEGGK`-Biotin, and NH_2_-`DLPGGK`-Biotin; synthetic standards Ac-`AE`(EDANS)`FR`, Ac-`AE`(EDANS)`FRTTSQGGK`-Biotin, NH_2_-`TTTTTSQGGK`-Biotin, NH_2_-`TTSQGGK`-Dabcyl, and NH_2_-ELGGGNAVEGGK-Biotin; and FRET-based substrates Ac-`AE`(EDANS)`FRTTSQGGK`-Dabcyl and Ac-`AE`(EDANS)`FCitTTSQGGK`-Dabcyl were all custom synthesized to >98% purity by Biomatik Co. (ON, Canada). Dynabeads™ MyOne™ Streptavidin C1 (Invitrogen, Thermo Fisher Scientific, USA; Cat. No. 65001, Lot No. 3165522) were used for peptide enrichment. Biotin-PEG4-alkyne (BroadPharma, USA; Cat. No. BP-22684, Lot No. B130-175). BTTP (BroadPharma, USA; Cat. No. BP-26133, Lot No. 2024051). The peptide NH_2_-`TTSQGGK`(N₃)YGRKKRRQRRRK-5FAM (purity 97.7%) was synthesized by CPC Scientific Inc. (CA, USA; Product No. 890350, Lot Nos. 05-01668 and CX-09-01087).

### Enzyme sources

Human cathepsins K and V were expressed in *Pichia pastoris* and purified as described in (*1-3*) and human cathepsin S was expressed in E. coli BL21(DE3) *(4)*. All enzymes were titrated with E64 prior to use (*3*). Human recombinant cathepsin B was kindly provided by J. Mort^†^ from the Shriner’s Hospital for Sick Children (Montreal, QC, Canada). Human liver cathepsin L was purchased from Millipore (Cat. No. 219402-25UG, Lot No. 4080104).

### Enzyme Kinetics using FRET substrates

Peptides Ac-`AE`(EDANS)`FRTTSQGGK`(Dabcyl) and Ac-`AE`(EDANS)`FCitTTSQGGK`(Dabcyl) were used as FRET substrates. Assays were performed with 30 nM cathepsins in 0.1 M sodium phosphate (pH 6.5) containing 2.5 mM dithiothreitol (DTT) at 25 °C. Substrate concentrations ranged from 1–50 µM. Fluorescence was monitored on a Synergy H1 microplate reader (BioTek; excitation 336 nm, emission 490 nm). Michaelis–Menten parameters (K_m_, *k*_cat_) were obtained by nonlinear regression (SigmaPlot or GraphPad Prism). Assays were performed in triplicate.

### Enzymatic reactions using FRET substrates and modified donor peptides for LC-MS/MS analysis

Reaction mixtures contained 20 µM FRET substrate (e.g., Ac-`AE`(EDANS)`FRTTSQGGK`-Dabcyl), 200 µM biotinylated peptide, and 2.5 mM DTT in 0.1 M sodium phosphate buffer (pH 6.5), 1 mM biotinylated or Dabcyl containing donor peptide (NH_2_-`TTSQGGK`-Biotin) or NH_2_-`TTSQGGK`-Dabcyl)) and 50 nM cathepsin. Reactions were incubated at 25 °C for 2 h in the dark and quenched with E-64 (10× of molar enzyme concentration). Cathepsin were precipitated by adding methanol (>80% v/v), stored at −20 °C for 2 h, and centrifuged (15 min). Supernatants were collected, dried by SpeedVac, reconstituted by TFA 0.5% and desalted using StageTips prior to LC-MS/MS analysis (*5*). For time-course experiments, aliquots were removed at 10 min, 30 min, 1 h, 2 h, and 5 h, inhibited with E-64, subjected to methanol precipitation, and cleaned on StageTips as above prior to LC-MS/MS.

### Enzymatic reactions with synthetic biotinylated peptides

Enzymatic reactions with synthetic biotinylated peptides were performed and analyzed by LC–MS/MS. Independent reaction mixtures contained Spike subunits (5 µg each of S1 and S2) combined with either hPF4 (10 µg), insulin (25 µg), or proinsulin (10 µg). Each mixture also contained 2.5 mM DTT in 0.1 M sodium phosphate buffer (pH 6.5), 1 mM biotinylated peptide NH_2_-`TTSQGGK`-Biotin, NH_2_-`NAVEGGK`-Biotin, or NH_2_-`DLPGGK`-Biotin and 50 nM cathepsins. Reactions were incubated at 25 °C for 2 h in the dark and quenched with E-64 (10-fold molar excess relative to enzyme). Proteins were precipitated by addition of methanol (>80% v/v), stored at −20 °C for 2 h, and centrifuged at 14,000 × g for 15 min. Supernatants were collected, dried in a SpeedVac, reconstituted in 0.5% TFA, and desalted using StageTips prior to LC–MS/MS analysis.

### Enzymatic reactions for protein–protein analyzed by LC–MS/MS

Reaction mixtures contained Spike subunits (5 µg each of S1 and S2) together with either hPF4 (10 µg) or insulin (25 µg) in 0.1 M sodium phosphate buffer (pH 6.5) supplemented with 2.5 mM DTT and 50 nM cathepsin. Reactions were incubated at 25 °C for 2 h in the dark and quenched with E-64 (10-fold molar excess relative to enzyme). Proteins were precipitated by addition of methanol (>80% v/v), stored at −20 °C for 2 h, and centrifuged at 14,000 × g for 15 min. Supernatants were collected, dried in a SpeedVac, reconstituted in 0.5% TFA, and desalted using StageTips prior to LC–MS/MS analysis.

### General data processing and analysis for insulin/hPF4 and Spike cis/transpeptide discovery

A custom Python pipeline was developed to analyze trans-spliced peptides from mass spectrometry data exported from PEAKS Studio. The analysis began by importing CSV data from three control and three experimental replicates. An initial quality control step retained only high-confidence entries by filtering for peptides with an Analytical Confidence (ALC) score greater than 90%. To ensure accuracy and mitigate false-positive identifications that often arise from isobaric amino acid mismatches, specifically between Leucine (L) and Isoleucine (I), a standardization protocol was implemented to create a canonical sequence for each peptide. A critical component of the pipeline was a manual verification step, which involved reanalyzing selected candidate peptide in FreeStyle software (Thermo Fisher Scientific). This process entailed extracting the monoisotopic peptide mass and meticulously examining the corresponding tandem mass spectra to confirm the peptide’s identity and structural integrity, thereby significantly increasing the fidelity of the identified peptides.

### Transpeptidase assays using biotinylated peptides as donors and proteins as acceptors

Reactions contained 50 nM enzyme, 16 µg protein substrate (e.g., bovine serum albumin (BSA), spike subunits, S1 or S2), and 1 mM biotinylated peptide (e.g., NH_2_-`TTSQGGK`-Biotin in 5% DMSO) in various pH buffer systems: 0.1 M sodium acetate (pH 4.5, 5.5), 0.1 M sodium phosphate (pH 6.5), or 0.1 M HEPES (pH 7.5) each containing 2.5 mM DTT. Incubations were performed at room temperature for 2 h in the dark. Reactions were terminated with 10 µM E64. For controls, enzymes were pre-incubated with E-64 for 5 min prior to substrate addition. Final DMSO concentrations were <1%.

Reaction products diluted by SDS sample buffer to the final concentration of 1.2% SDS, 1% glycerol, 0.001% bromophenol blue, 0.03 M Tris-HCl, pH 6.8, and 2.5% (v/v) 2-mercaptoethanol) heated at 100 °C for 5 min. Samples resolved by SDS-PAGE (15% polyacrylamide gels; 10–30 µg protein per well) and transferred onto nitrocellulose membranes (Amersham™ Protran™, GE Healthcare) using a Mini-PROTEAN Tetra Cell (Bio-Rad; 90 mA/gel, 120 min) in ice-cold transfer buffer (2.9 g glycine, 5.8 g Tris, 200 mL methanol per 1 L, pre-chilled at –20 °C ≥2 h). Membranes were blocked overnight at 4 °C in 1% (w/v) skimmed milk in Tris-buffered saline with 0.1% Tween 20 (TBST) (50 mM Tris, 150 mM NaCl, 0.1% Tween-20, pH 7.5), washed in TBST (4 × 5 min) and Phosphate-buffered salin (PBS) (2 × 5 min), and probed with avidin–FITC (25 µg/mL in PBS, 1 h, room temperature with gentle rocking). Excess probe was removed by PBS + 1% Tween-20 wash (1 × 5 min), followed by PBS wash (2 × 5 min). Fluorescence was imaged on an Amersham Typhoon scanner (excitation 488 nm).

### LC-MS/MS analysis

#### Liquid Chromatography

Samples were injected and separated on-line using an Easy-nLC 1200 (Thermo Fisher Scientific) with Aurora Series analytical column, (25cm x 75μm 1.6μm C18; Ion Opticks, Parkville, Victoria, Australia). The analytical column was heated to 40°C using an integrated column oven (PRSO-V2, Sonation, Biberach, Germany). Buffer A consisted of 0.1% aqueous formic acid and 2% acetonitrile in water, and buffer B consisted of 0.1% aqueous formic acid and 80% acetonitrile in water. A standard 60 min gradient was run from 2% B to 20% B over 46 min, then to 32% B over 15 min, then to 50% B from 61 to 66min, then to 95% B over 5 min, held at 95% B for 8 min, then dropped to 3% B over 2 min, held at 3% B for 6 min. The analysis was performed at 0.25 μL/min flow rate. The Easy-nLC thermostat temperature was set at 7°C.

#### MS/MS Acquisition Method

The peptides were analyzed with an Orbitrap Exploris 480 mass spectrometer (Orbitrap Exploris^TM^ 480, Thermo Fisher Scientific). The Nanospray Flex^TM^ ion source was operated at 1900 V spray voltage and ion transfer tubes were heated to 290°C. During analysis, the Orbitrap Exploris 480 was operated in a data-dependent (DDA) mode with 20 dependent scans. The MS and MS/MS spectra were collected in positive mode. Full MS resolution was set to 60,000 with a normalized automatic gain control (AGC) target of 100%, 50% RF lens and 20 ms maximum injection time. Scan range from m/z 375 Th to m/z 1200 Th, and charge state from 2-5 were included. Dynamic exclusion was enabled to exclude after 1 time for 5 s. For MS/MS scans, the resolution was set to 15,000, with normalized AGC target at 50%, normalized HCD collision energy at 28%, and isolation windows at m/z 2 Th.

#### MS/MS data analysis

Raw LC–MS/MS data were processed in PEAKS Studio 12.5 (Bioinformatics Solutions Inc.) under CID–DDA mode. Peptide lists were exported as .csv files and analyzed with custom Python scripts (pandas, numpy, scipy). Peptides were at ALC ≥ 90%, and restricted to sequences ≥4 amino acids. Sequences were canonicalized for I/L isobaric equivalence and matched against precompiled human proinsulin/PF4 and SARS-CoV-2 Spike fragments. Peptides were classified as single fragments, cispeptides (non-contiguous within one protein), or transpeptides (insulin/PF4–Spike hybrids), and further designated as first- or second-generation by number of junctions. To ensure data integrity and minimize false-positive peptide assignments arising from algorithmic misidentification, all candidate peptides underwent stringent manual validation. Reanalysis was performed in FreeStyle (Thermo Fisher Scientific) by extracting the monoisotopic peptide mass and critically inspecting the corresponding tandem mass spectra. Validation included extracted ion chromatograms (±5 ppm), and for tagged peptides, diagnostic fragment ions were specifically monitored: biotinylated peptides at m/z 227.0845 and 310.158, EDANS at m/z 333.0904, Dabcyl at m/z 252.1131, and 4PEG-Biotin with a precursor ion at m/z 483.2152 and characteristic fragment ions at m/z 270.1276 and 227.08 (Fig. S1).

### *In vitro* binding assay to HLA-DQ8

The binding of selected fusion peptides to the HLA-DQ8 molecule (DQA1*03:01–DQB1*03:02) was evaluated by ProImmune Ltd. (Oxford, UK) using the ProImmune REVEAL™ MHC–peptide proprietory binding assay platform. The cis/transpeptides identified *in vitro* by LC–MS/MS defined the fusion junctions between peptide fragments but were shorter than the length typically used for HLA class II binding assays. Therefore, peptide sequences were extended according to the corresponding parent protein sequences to generate 15-mer peptides centered around the identified fusion junctions. The resulting peptides (GQVELGGGNAVEVLK, GIVEQCCTNAVEVLK, HLVEALYLNAVEVLK, EALYLVCGNAVEVLK, IAVEQDKNNAVEVLK, and YNENGTITNAVEVLK) were synthesized and evaluated for binding to HLA-DQ8. Peptide binding to recombinant HLA molecules was assessed by monitoring the formation of correctly folded peptide–HLA complexes using a labeled conformation-dependent antibody that recognizes the native structure of the complex. Binding signals generated by test peptides were quantified relative to a reference control peptide and reported as a percentage of the control signal according to the manufacturer’s criteria.

### RAW 264.7 mouse macrophages

RAW 264.7 cells were maintained in RPMI-1640 medium supplemented with 10% fetal bovine serum (FBS) and 1% penicillin-streptomycin at 37 °C in 5% CO₂. Cells were detached with 0.05% trypsin-EDTA (5 min, 37 °C), neutralized with complete medium, pelleted (300 × g, 5 min), and resuspended in fresh medium. Subcultures were performed every 2–3 days at 80–90% confluence. For experiments, 1.5 × 10⁵ RAW 264.7 cells were seeded per well in 24-well plates and allowed to adhere overnight. Cells were then treated with 10 µM fluorogenic TAT peptide (NH_2_-`TTSQGGK`(N_3_)YGRKKRRQRRRK-5-FAM) in the presence or absence of the cysteine protease inhibitor E-64d (10 µM) and incubated for 4 h. Two experimental sets were prepared: one for immunofluorescence staining and the other for mass spectrometry analysis. For mass spectrometry, cells were lysed according to CT-TRAP workflow. For immunofluorescence, cells were washed three times with PBS (2 min each), fixed with 4% formaldehyde for 10 min, washed again, mounted, and imaged using a Leica SP5 laser scanning confocal inverted microscope (Leica microsystems, Wetzlar, Germany). Negative controls were included to normalize and assess peptide uptake.

### CT-TRAP workflow

Cells were lysed in 8 M urea, 50 mM HEPES (pH 7.5) supplemented with an EDTA-free protease inhibitor cocktail. Lysates were sonicated in a water bath for 10 min, clarified by centrifugation, and diluted to 0.8 M urea prior to copper-catalyzed azide–alkyne cycloaddition (CuAAC). Click reactions were initiated with 100 µM 4PEG-alkyne, followed by addition of a premixed solution of 1 mM CuSO₄ and 2 mM Benzotriazolylmethyl-tris(2-pyridylmethyl)amine (BTTP), and immediate addition of 10 mM sodium ascorbate. Reactions were incubated for 1 h at room temperature in the dark and quenched with 500 µM Ethylenediaminetetraacetic acid (EDTA). Samples were concentrated by SpeedVac, desalted using StageTips (70% acetonitrile elution), redissolved in PBS (150 µL), and incubated with MyOne C1 streptavidin beads (40 µL, 1 h). Beads were washed five times with PBS, and bound material was eluted with 6 M guanidinium–HEPES (pH 7.5, 200 µL) at 50 °C for 1 min. Eluates were acidified (final pH <3) and subjected to StageTip cleanup (40% acetonitrile elution) prior to LC-MS/MS.

## Author contributions

SYTD and DB designed the study, validated the results, and oversaw project administration. SYTD performed enzyme kinetics and all LC-MS/MS analyses, while JR, OH, and LF carried out sample preparation and instrumental work for LC-MS/MS. SYTD designed and developed the CT-TRAP method. PP and OH were responsible for all cell culture and confocal microscopy experiments. EM produced and purified recombinant human CatK, and SYTD produced and purified human CatS. SYTD and DB wrote the original draft of the manuscript. All authors reviewed and revised the final manuscript. DB acquired funding for the study.

## Acknowledgements

This work was supported by Canadian Institutes of Health Research grants (PJT-155979, CPG-158275), Collaborative Health Research Project (CHRP 523434-18) and the NSERC discovery grant (LJGP GR003266). DB was supported by Canada Research Chair award funding. We are thankful to the proteomics core facility at UBC under the leadership of L. Foster and the technical support by J. Rogalski to generate the raw LC-MS/MS data.

## Conflict of interest statement

No disclosures and all authors have no conflicts of interest.

## Notes

### Competing Interest Statement

The authors have declared no competing interest.

### Summary of Updates

The structure of Introduction, Discussion and results has been revised. The binding prediction to HLA molecules has been removed. The peptide discovery in mice splenocyes has been removed.

