## Supplementary material for "Reverse Proteolysis Uncovers a Hidden Dimension of the Peptidome": Supplimentary Info

S. Yasin Tabatabaei Dakhili *et al.*

#### Signature Ions table

| Conjugated Amino acid | Molecular Structure of Marker Ions with Calculated *m/z* |
|---|---|
| Lysine-Biotin | 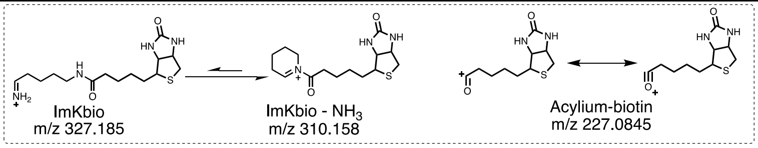  |
| Glutamic acid -EDANS | 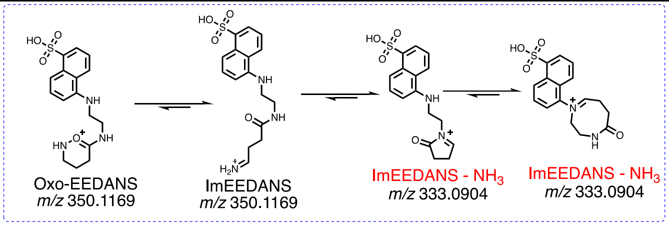  |
| Lysine-Dabcyl | 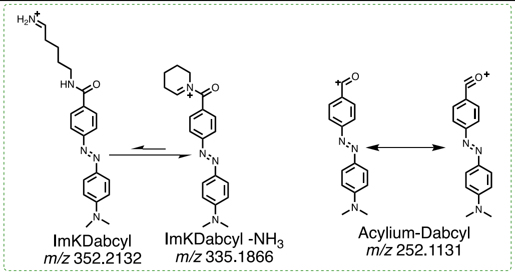  |

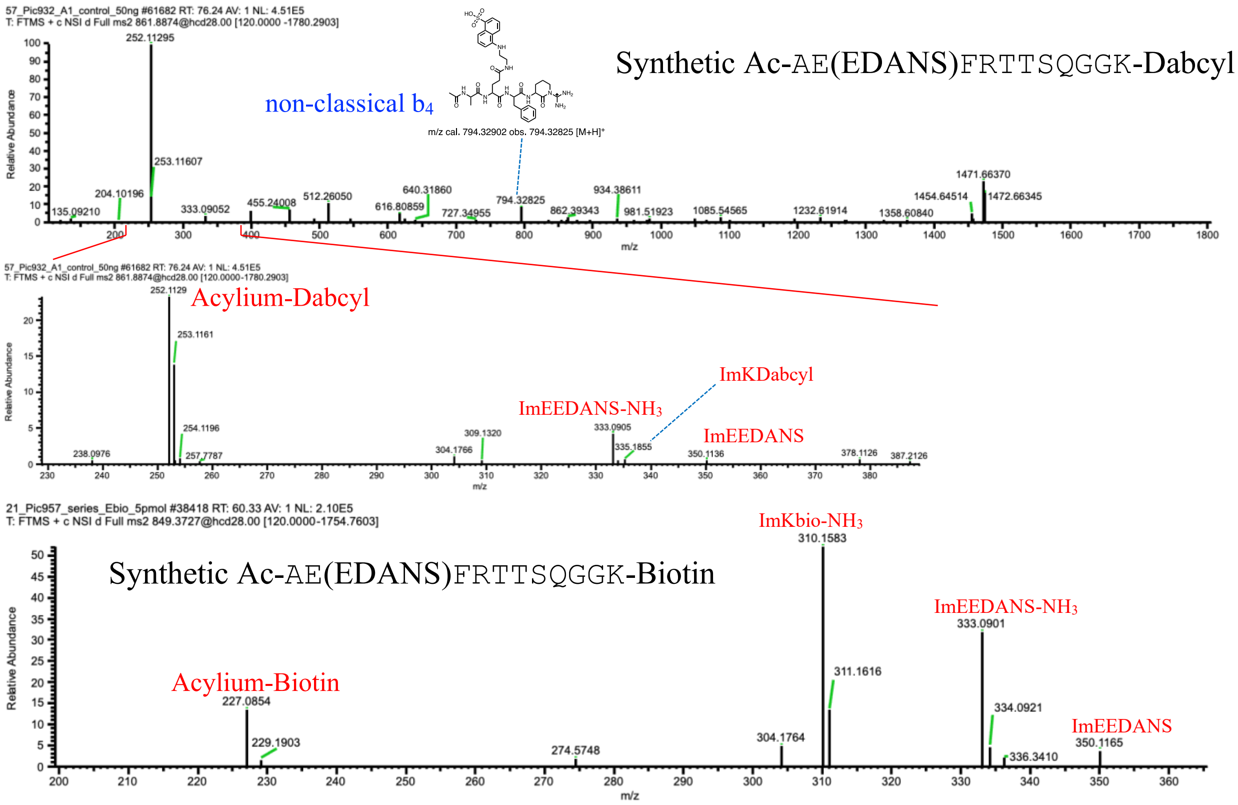

**Fig. S1:** The table presents the molecular structures of molecular ions of conjugated peptide-tags—EDANS, Dabcyl, and Biotin—along with the theoretical *m/z* values of their major marker ions. The graphs below show tandem mass spectrum of the corresponding synthetic peptides, with major observed marker ions highlighted in red (*5, 6*).

#### Characterization of EDANS–Dabcyl cis/trans-peptides

##### Ac-AE(EDANS)FRTTTTSQGGK(Biotin)

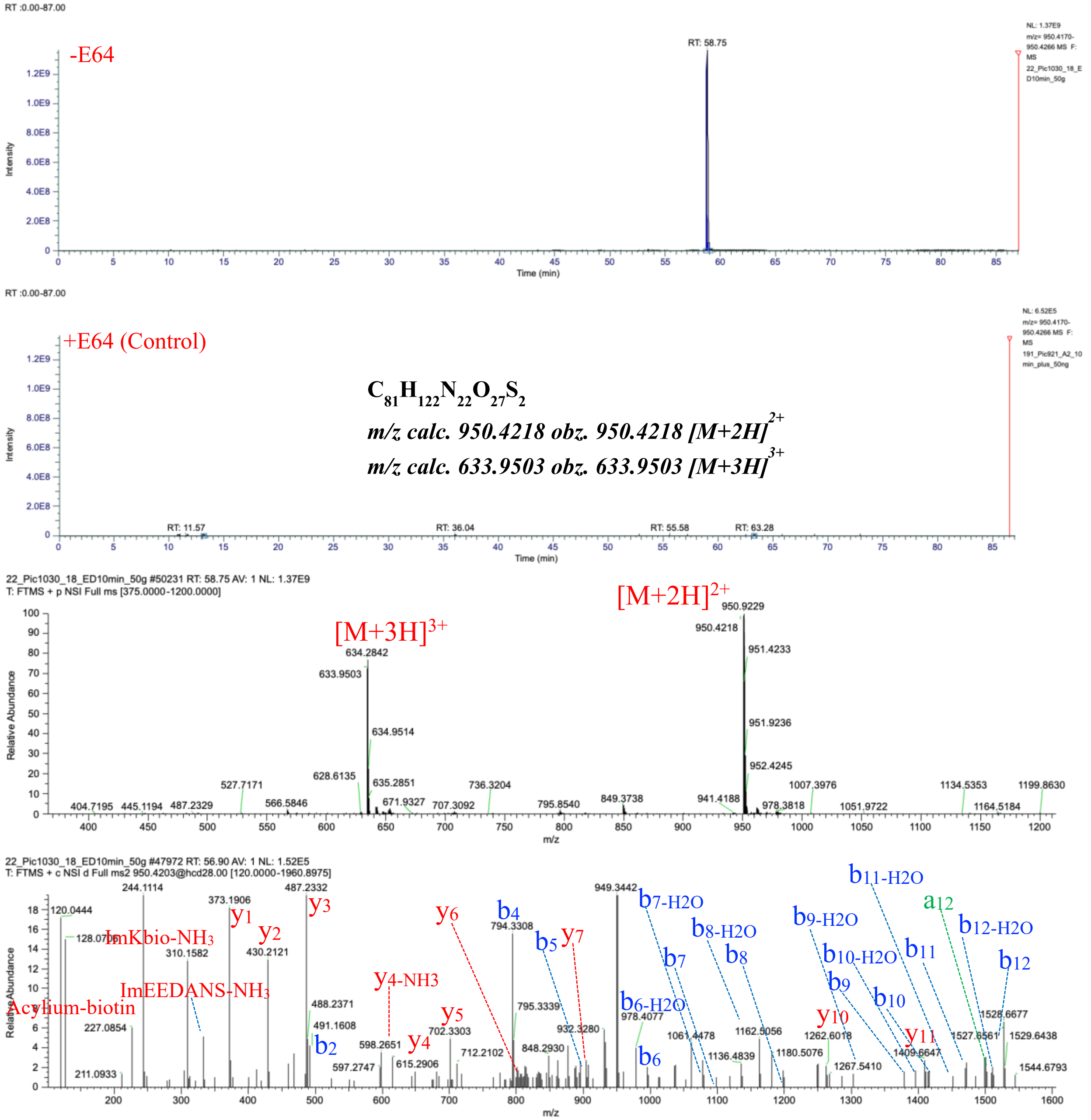

**Fig. S2.** Ac-AE(EDANS)FRTTTTSQGGK(Biotin) characterized by LC-MS/MS as a product of hCatS-catalyzed transpeptidation reaction. The figure displays a graphical representation of the extracted ion chromatogram (EIC), mass spectrum, and tandem mass spectrum of Ac-AE(E)FRTTTTSQGGK(Biotin), identified from a reaction between Ac-AE(E)FRTTSQGGK(D) and TTSQGGK(Biotin) in the presence of active hCatS. LC-MS/MS analysis was performed on samples treated with active hCatS (–E64) and compared to a control in which the enzyme was inhibited by E64 (+E64).

Ac-AE(EDANS)FRTTSQGGK(Biotin)

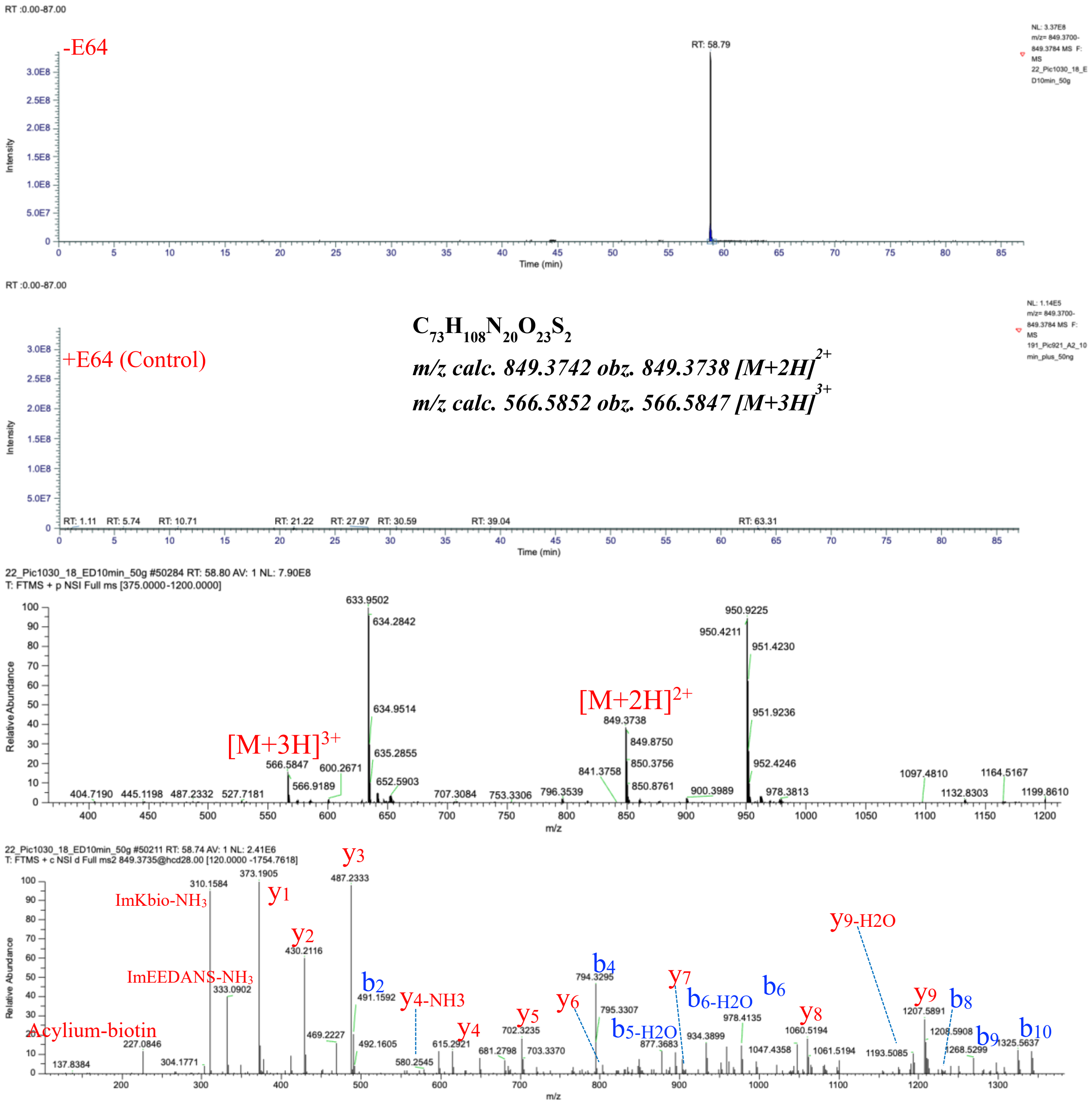

***Fig. S3***. *Ac-AE(EDANS)FRTTSQGGK(Biotin)* characterized by LC-MS/MS as a product of hCatS-catalyzed transpeptidation reaction. The figure displays a graphical representation of the extracted ion chromatogram (EIC), mass spectrum, and tandem mass spectrum of Ac-AE(E)FRTTSQGGK(Biotin), identified from a reaction between Ac-AE(E)FRTTSQGGK(D) and TTSQGGK(Biotin) in the presence of active hCatS. LC-MS/MS analysis was performed on samples treated with active hCatS (–E64) and compared to a control in which the enzyme was inhibited by E64 (+E64).

*Ac-AE(EDANS)FRTTTSQGGK(Biotin)*

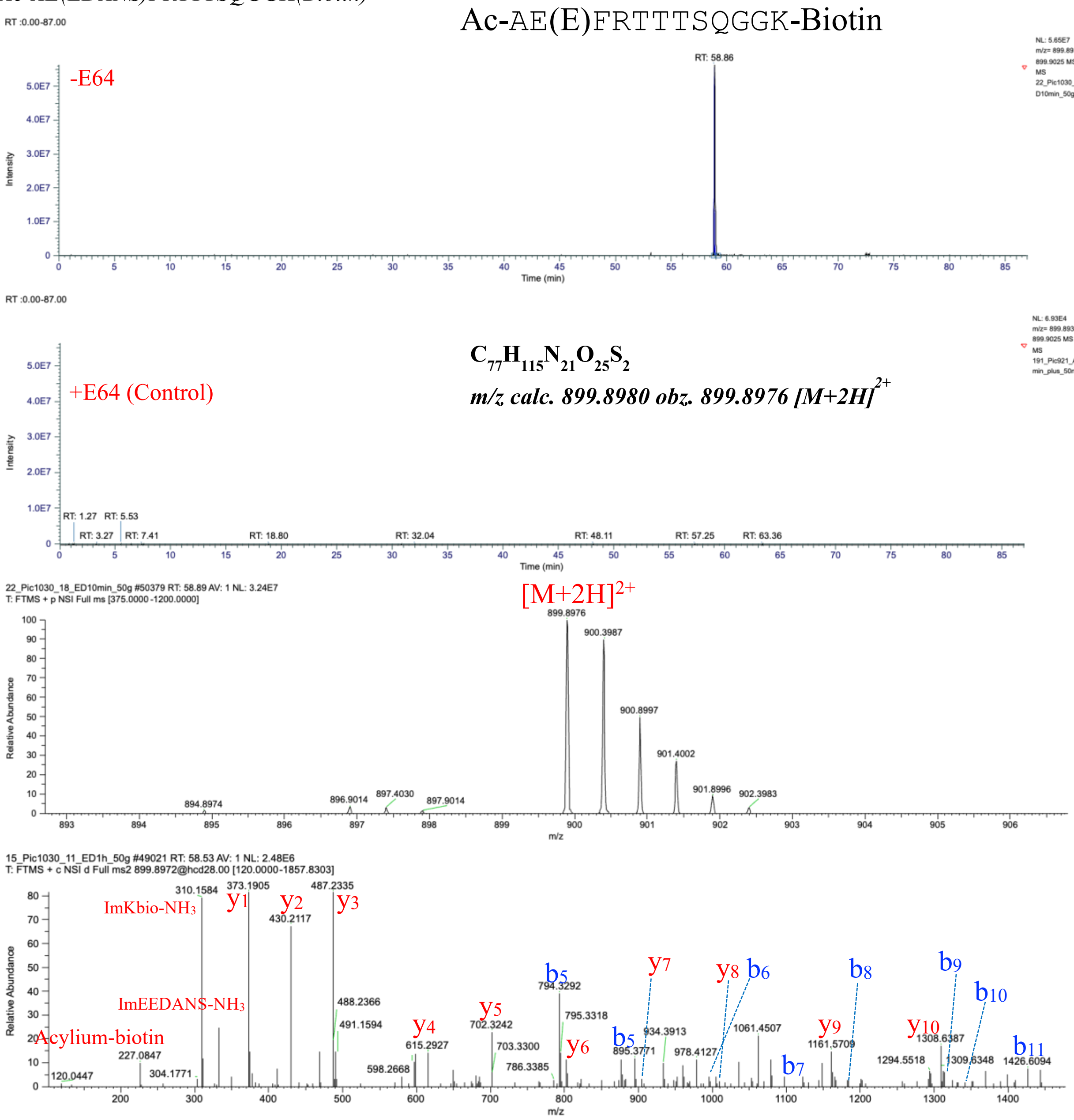

***Fig. S4***. *Ac-AE(EDANS)FRTTTSQGGK(Biotin)* characterized by LC-MS/MS as a product of hCatS-catalyzed transpeptidation reaction. The figure displays a graphical representation of the extracted ion chromatogram (EIC), mass spectrum, and tandem mass spectrum of Ac-AE(E)FRTTTSQGGK(Biotin), identified from a reaction between Ac-AE(E)FRTTSQGGK(D) and TTSQGGK(Biotin) in the presence of active hCatS. LC-MS/MS analysis was performed on samples treated with active hCatS (–E64) and compared to a control in which the enzyme was inhibited by E64 (+E64).

*Ac-AE(EDANS)FRTTTTTSQGGK(Biotin)*

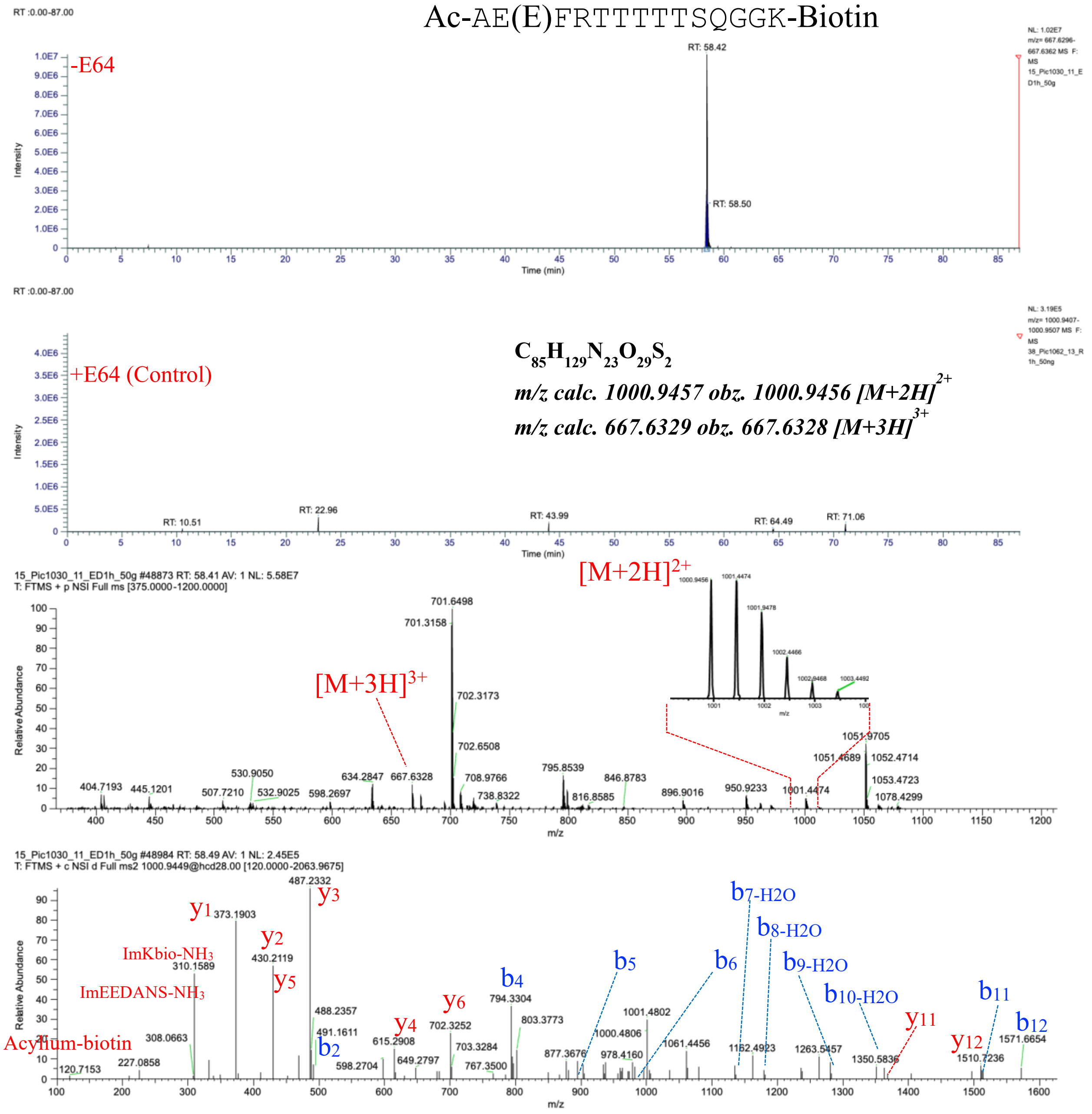

***Fig. S5***. *Ac-AE(EDANS)FRTTTTTSQGGK(Biotin)* characterized by LC-MS/MS as a product of hCatS-catalyzed transpeptidation reaction. The figure displays a graphical representation of the extracted ion chromatogram (EIC), mass spectrum, and tandem mass spectrum of Ac-AE(E)FRTTTTTSQGGK(Biotin), identified from a reaction between Ac-AE(E)FRTTSQGGK(D) and TTSQGGK(Biotin) in the presence of active hCatS. LC-MS/MS analysis was performed on samples treated with active hCatS (–E64) and compared to a control in which the enzyme was inhibited by E64 (+E64).

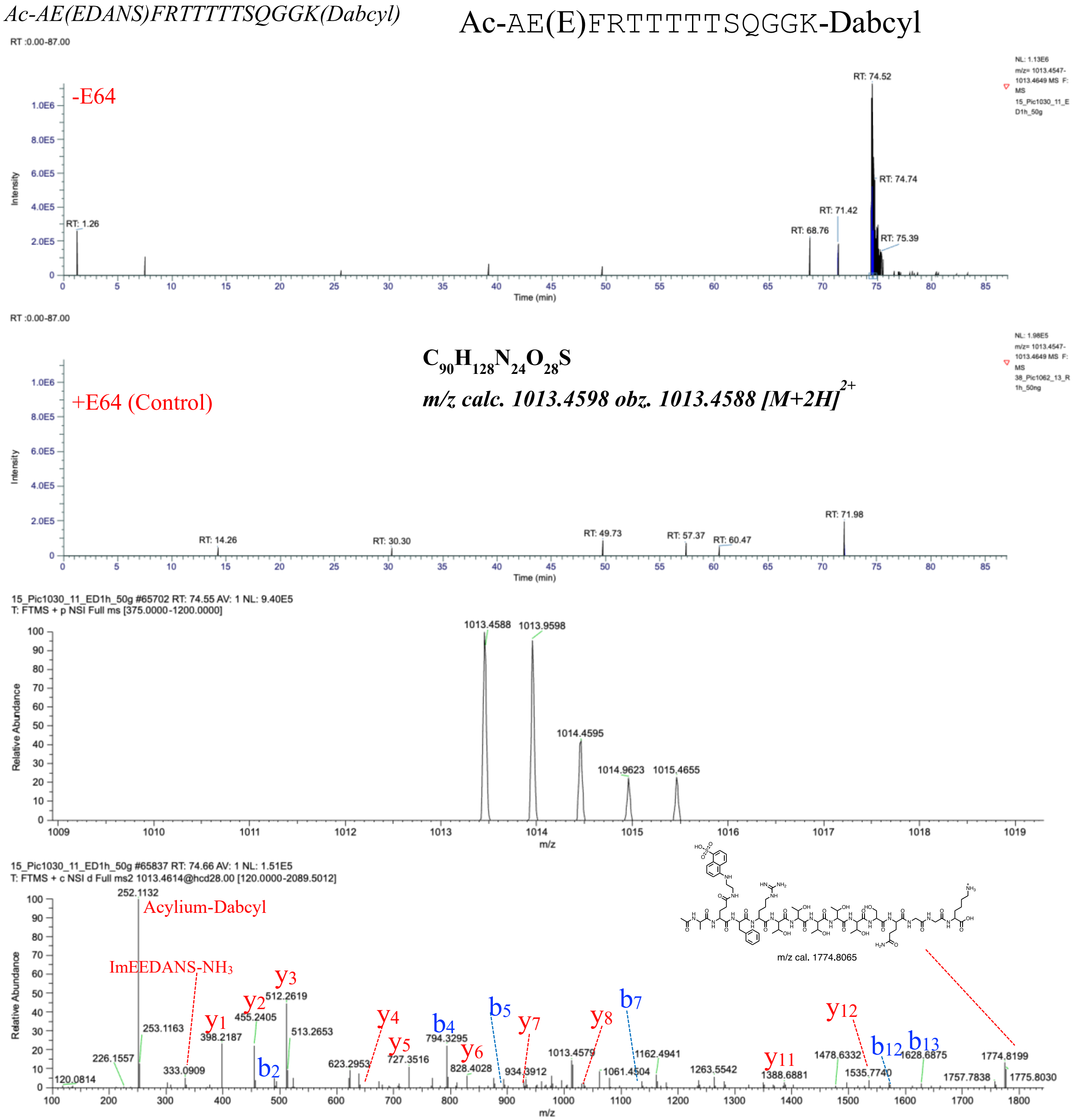

***Fig. S6***. *Ac-AE(EDANS)FRTTTTTSQGGK(Dabcyl)* characterized by LC-MS/MS as a product of hCatS-catalyzed transpeptidation reaction. The figure displays a graphical representation of the extracted ion chromatogram (EIC), mass spectrum, and tandem mass spectrum of Ac-AE(E)FRTTTTTSQGGK(Dabcyl), identified from a reaction between Ac-AE(E)FRTTSQGGK(D) and TTSQGGK(Biotin) in the presence of active hCatS. LC-MS/MS analysis was performed on samples treated with active hCatS (–E64) and compared to a control in which the enzyme was inhibited by E64 (+E64).

Ac-AE(EDANS)FRTTTTSQGGK(Dabcyl)

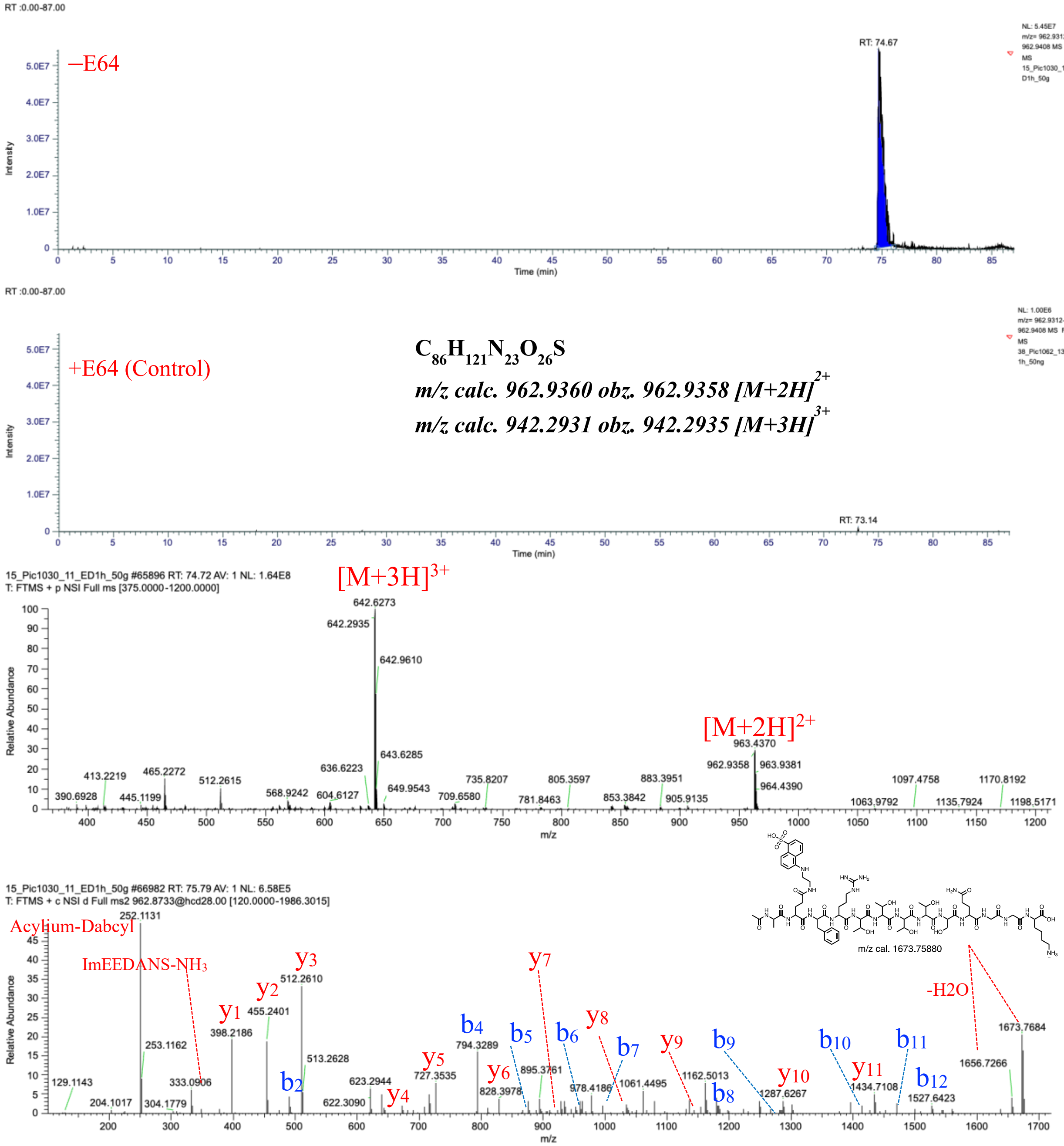

**Fig. S7.** Ac-AE(EDANS)FRTTTTSQGGK(Dabcyl) characterized by LC-MS/MS as a product of hCatS-catalyzed transpeptidation reaction. The figure displays a graphical representation of the extracted ion chromatogram (EIC), mass spectrum, and tandem mass spectrum of Ac-AE(E)FRTTTTSQGGK(Dabcyl), identified from a reaction between Ac-AE(E)FRTTSQGGK(D) and TTSQGGK(Biotin) in the presence of active hCatS. LC-MS/MS analysis was performed on samples treated with active hCatS (–E64) and compared to a control in which the enzyme was inhibited by E64 (+E64).

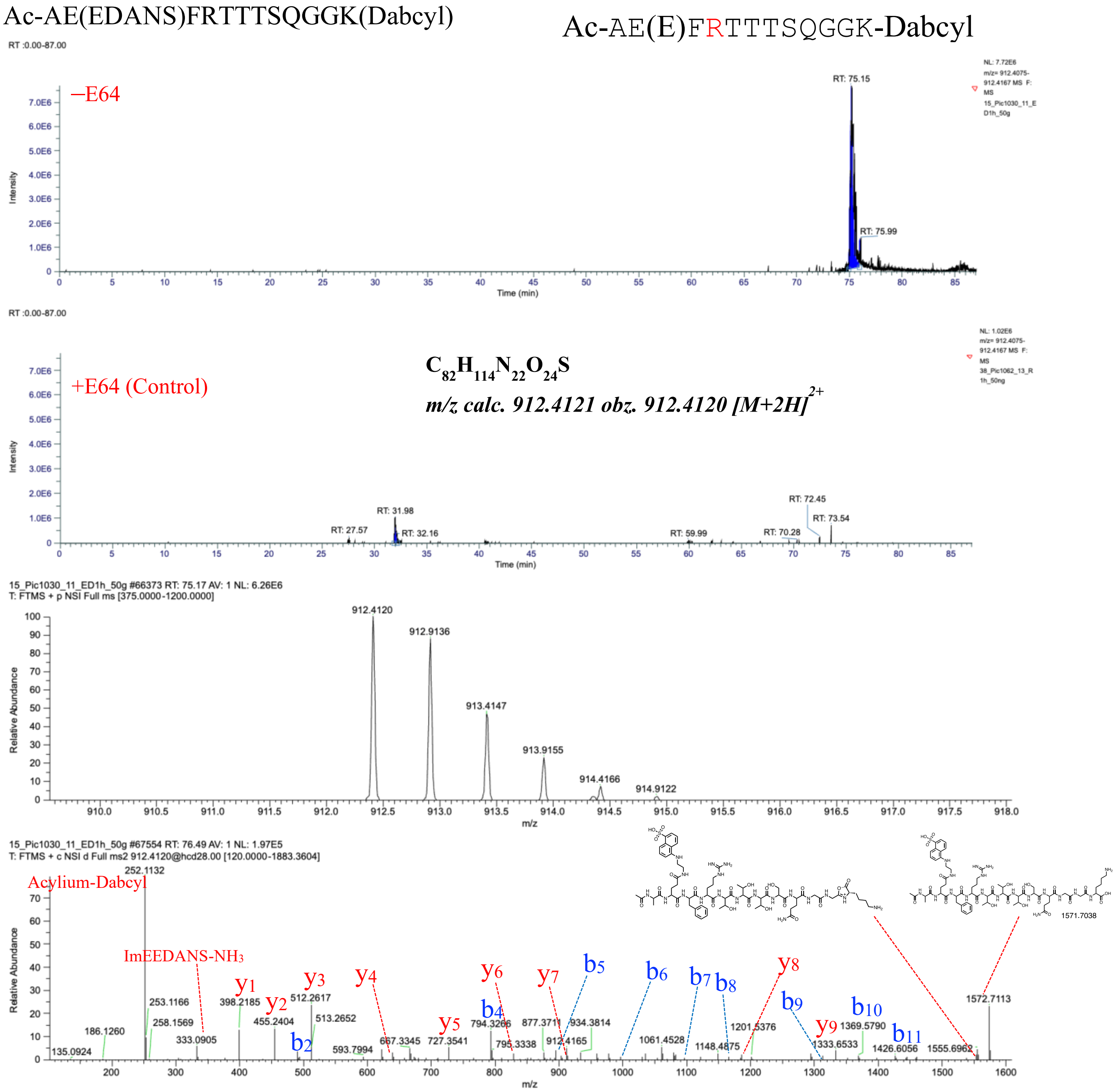

**Fig. S8.** Ac-AE(EDANS)FRTTTSQGGK(Dabcyl) characterized by LC-MS/MS as a product of hCatS-catalyzed transpeptidation reaction. The figure displays a graphical representation of the extracted ion chromatogram (EIC), mass spectrum, and tandem mass spectrum of Ac-AE(E)FRTTTSQGGK(Dabcyl), identified from a reaction between Ac-AE(E)FRTTSQGGK(D) and TTSQGGK(Biotin) in the presence of active hCatS. LC-MS/MS analysis was performed on samples treated with active hCatS (–E64) and compared to a control in which the enzyme was inhibited by E64 (+E64).

TTTSQGGK(Biotin)

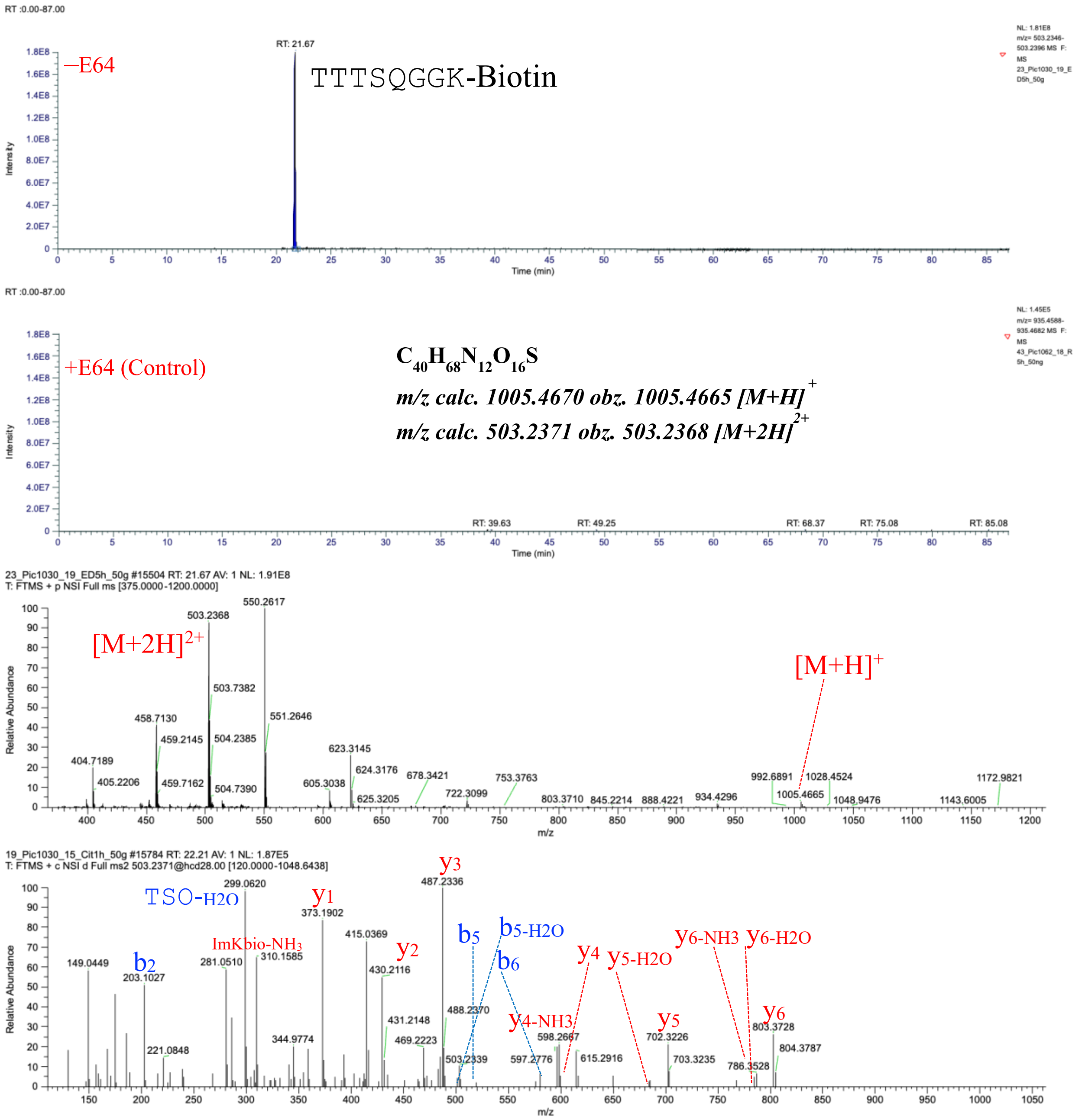

**Fig. S9.** TTTSQGGK(Biotin) characterized by LC-MS/MS as a product of hCatS-catalyzed transpeptidation reaction. The figure displays a graphical representation of the extracted ion chromatogram (EIC), mass spectrum, and tandem mass spectrum of TTTSQGGK(Biotin), identified from a reaction between Ac-AE(E)FRTTSQGGK(D) and TTSQGGK(Biotin) in the presence of active hCatS. LC-MS/MS analysis was performed on samples treated with active hCatS (–E64) and compared to a control in which the enzyme was inhibited by E64 (+E64).

TTTSQGGK(Dabcyl)

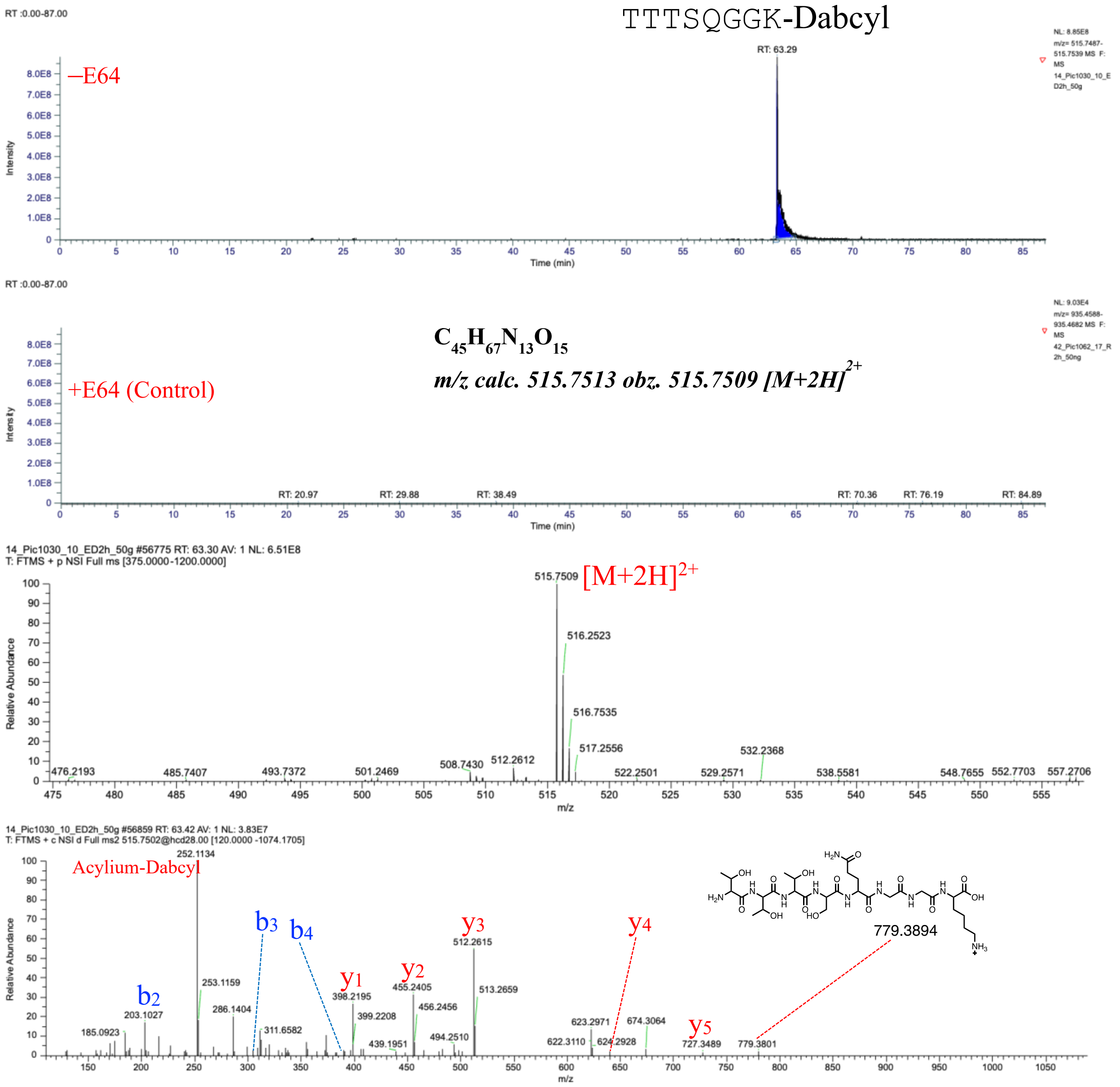

**Fig. S10.** TTTSQGGK(Dabcyl) characterized by LC-MS/MS as a product of hCatS-catalyzed transpeptidation reaction. The figure displays a graphical representation of the extracted ion chromatogram (EIC), mass spectrum, and tandem mass spectrum of TTTSQGGK(Dabcyl), identified from a reaction between Ac-AE(E)FRTTSQGGK(D) and TTSQGGK(Biotin) in the presence of active hCatS. LC-MS/MS analysis was performed on samples treated with active hCatS (–E64) and compared to a control in which the enzyme was inhibited by E64 (+E64).

TTTTSQGGK(Dabcyl)

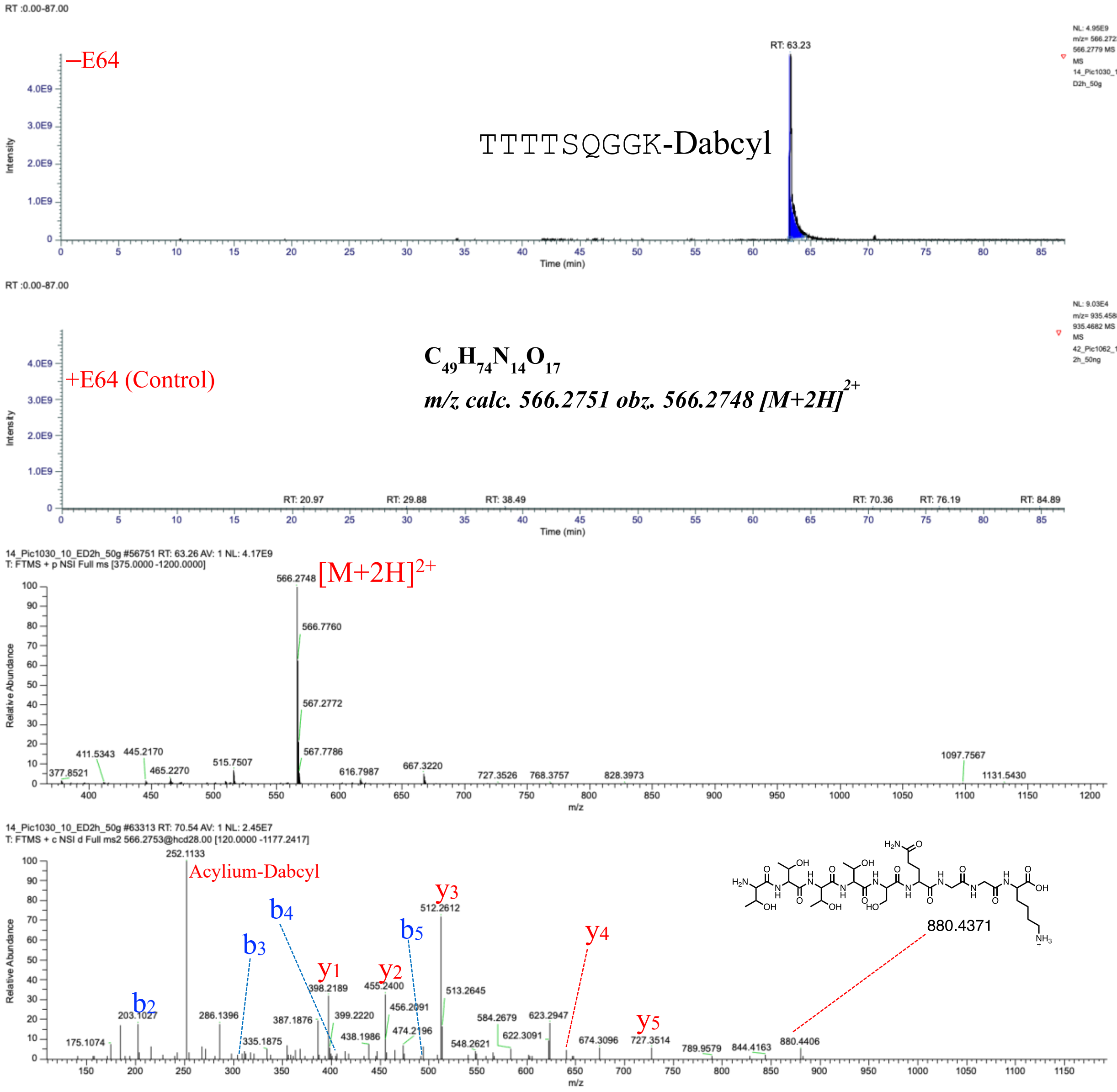

**Fig. S11.** TTTTSQGGK(Dabcyl) characterized by LC-MS/MS as a product of hCatS-catalyzed transpeptidation reaction. The figure displays a graphical representation of the extracted ion chromatogram (EIC), mass spectrum, and tandem mass spectrum of TTTTSQGGK(Dabcyl), identified from a reaction between Ac-AE(E)FRTTSQGGK(D) and TTSQGGK(Biotin) in the presence of active hCatS. LC-MS/MS analysis was performed on samples treated with active hCatS (–E64) and compared to a control in which the enzyme was inhibited by E64 (+E64).

TTTTTSQGGK(Dabcyl)

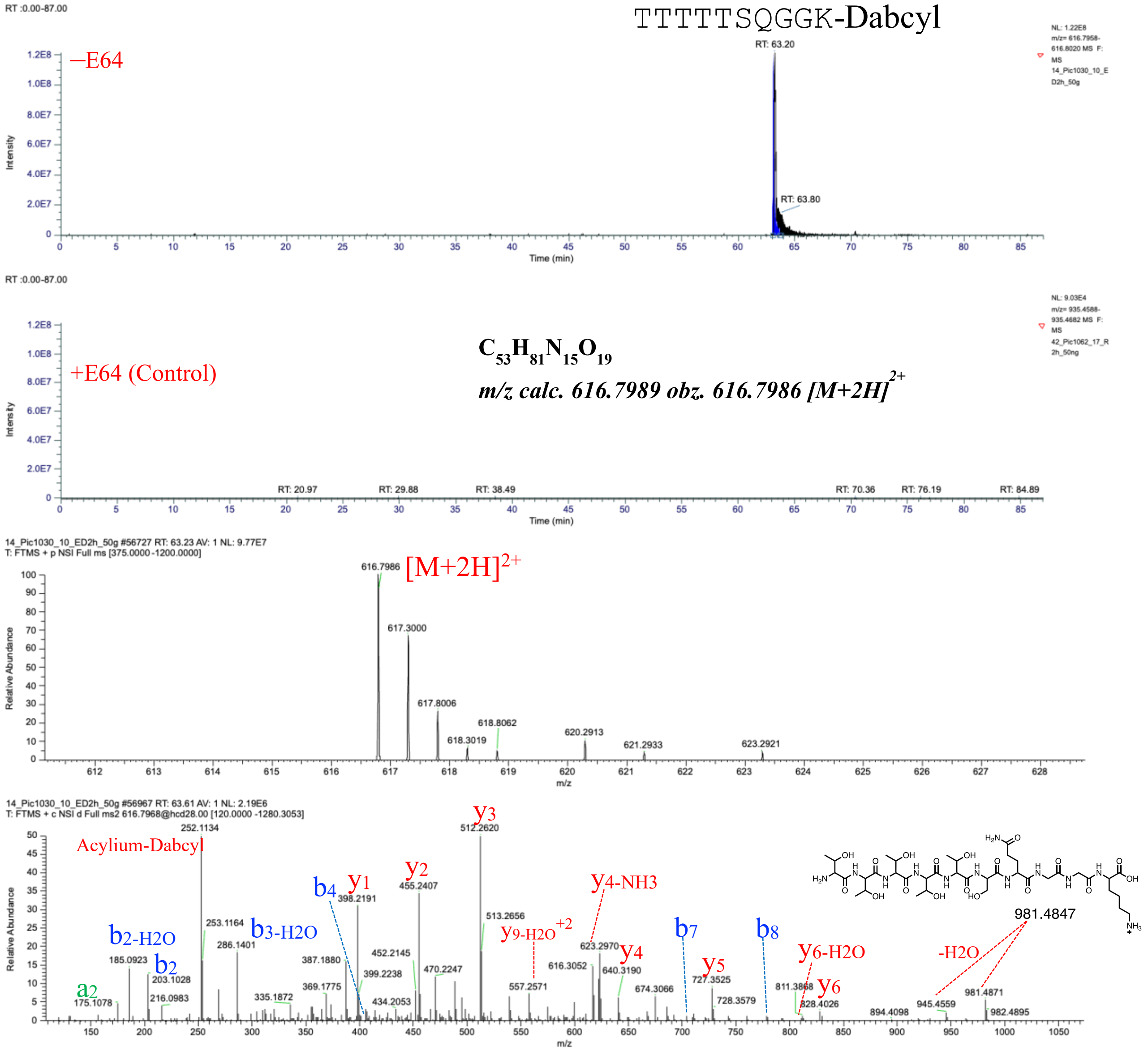

**Fig. S12.** TTTTTSQGGK(Dabcyl) characterized by LC-MS/MS as a product of hCatS-catalyzed transpeptidation reaction. The figure displays a graphical representation of the extracted ion chromatogram (EIC), mass spectrum, and tandem mass spectrum of TTTTTSQGGK(Dabcyl), identified from a reaction between Ac-AE(E)FRTTSQGGK(D) and TTSQGGK(Biotin) in the presence of active hCatS. LC-MS/MS analysis was performed on samples treated with active hCatS (–E64) and compared to a control in which the enzyme was inhibited by E64 (+E64).

#### Comparing chemically synthesized and enzyme catalyzed Ac-AE(E)FRTTSQGGK-Biotin

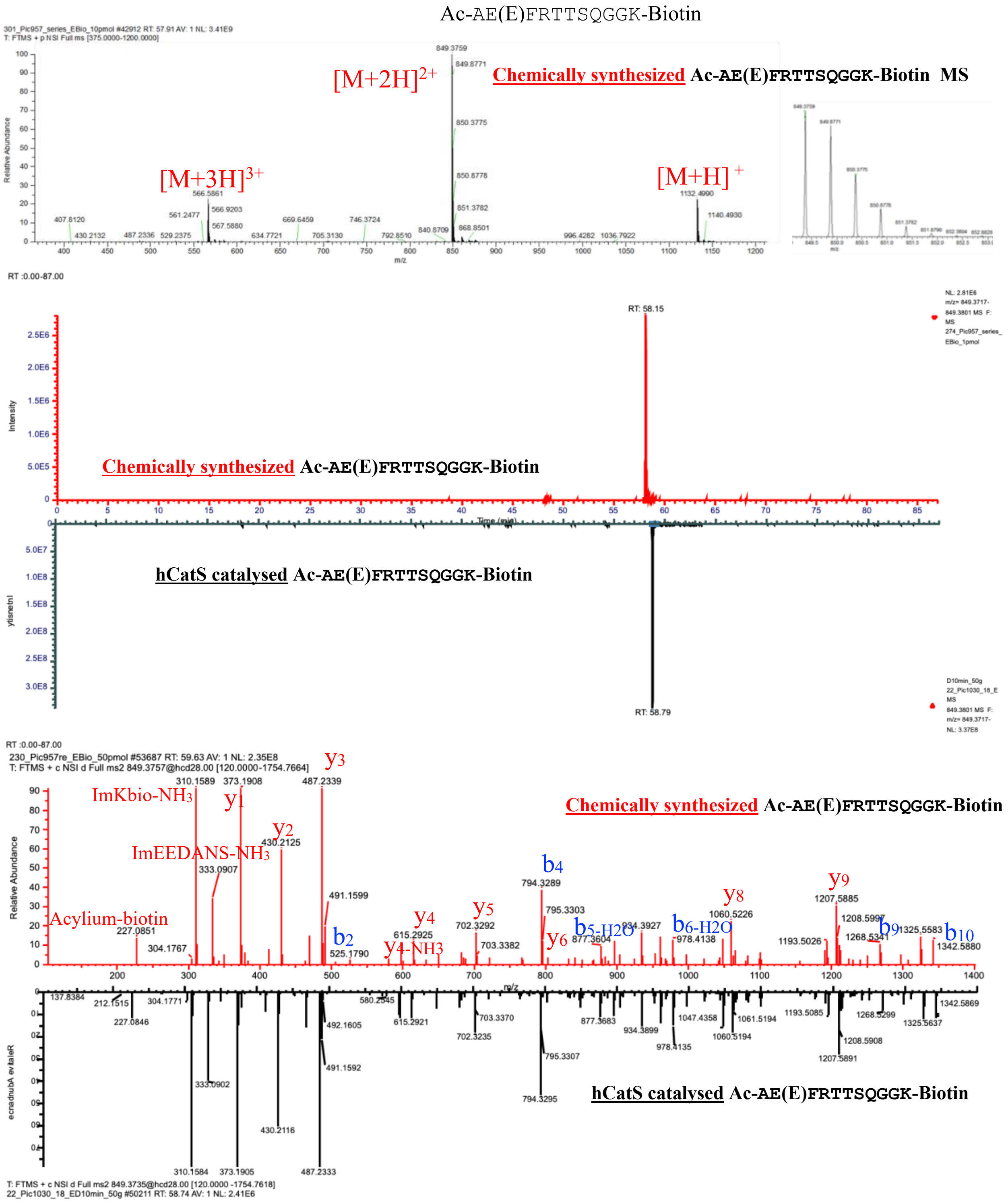

**Fig. S13.** The figure presents a comparison of the graphical representations of the extracted ion chromatogram (EIC), mass spectrum, and tandem mass spectrum between synthetic Ac-AE(E)FRTTSQGGK(Biotin) and the enzymatically produced counterpart.

#### Comparing chemically synthesized and enzyme catalyzed TTTTTSQGGK-Biotin

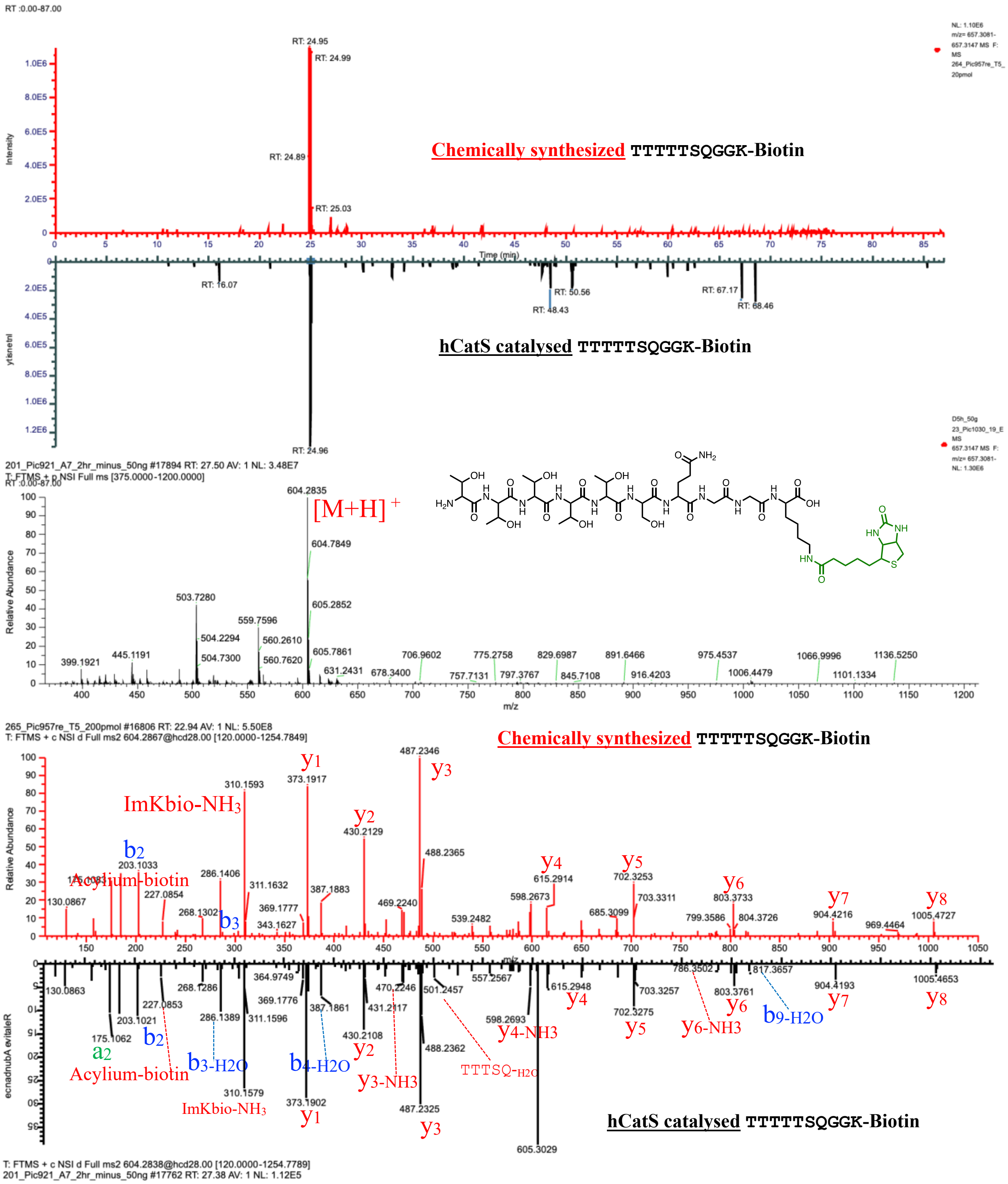

**Fig. S14.** The figure presents a comparison of the graphical representations of the extracted ion chromatogram (EIC), mass spectrum, and tandem mass spectrum between synthetic TTTTTSQGGK(Biotin) and the enzymatically produced counterpart.

Ac-AE(EDANS)FCitTTTTSQGGK(Dabcyl)

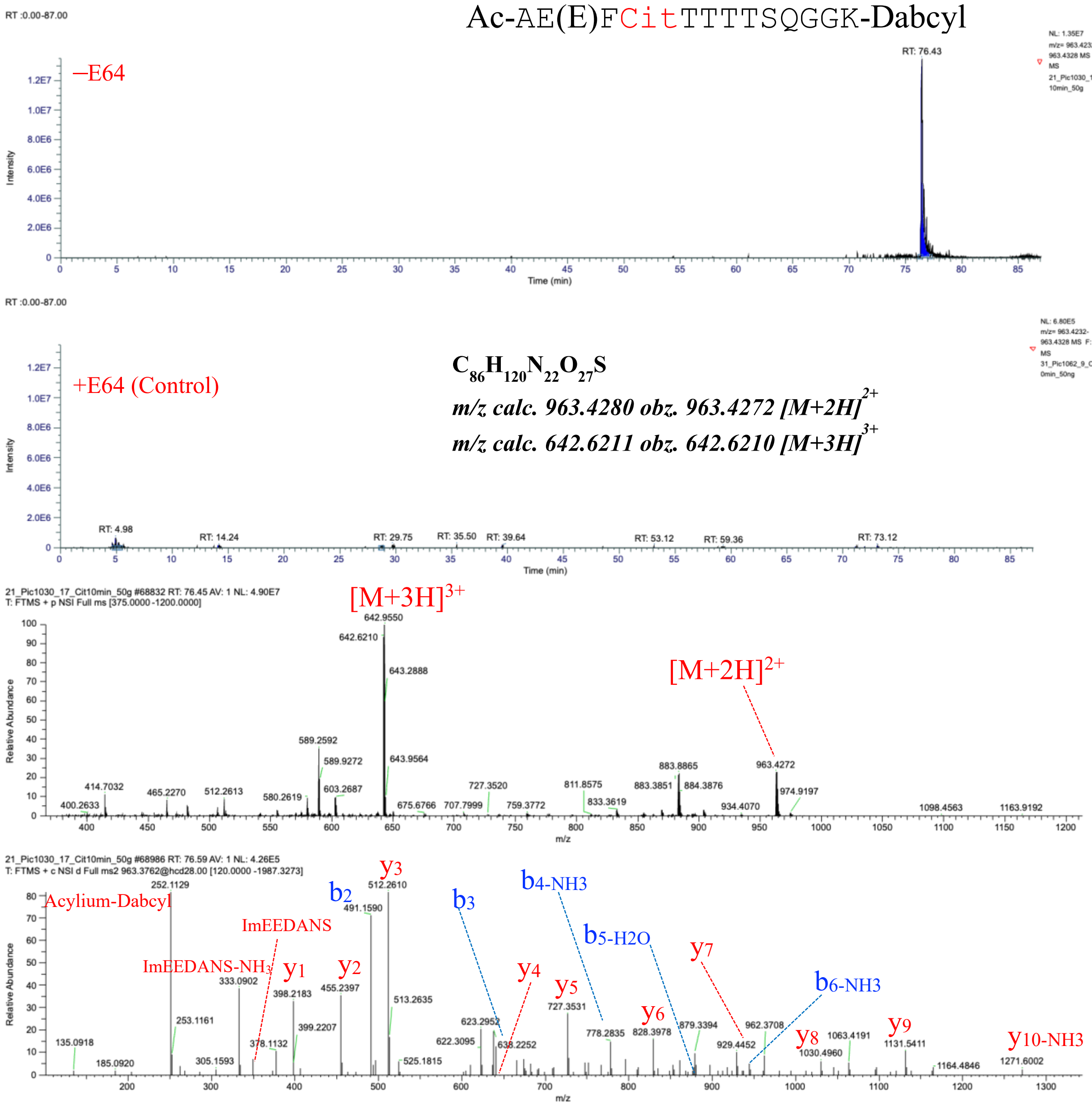

**Fig. S15.** Ac-AE(EDANS)FCitTTTTSQGGK(Dabcyl) characterized by LC-MS/MS as a product of hCatS-catalyzed transpeptidation reaction. The figure displays a graphical representation of the extracted ion chromatogram (EIC), mass spectrum, and tandem mass spectrum of Ac-AE(E)FCitTTTTSQGGK(Dabcyl), identified from a reaction between Ac-AE(E)FCitTTSQGGK(D) and TTSQGGK(Biotin) in the presence of active hCatS. LC-MS/MS analysis was performed on samples treated with active hCatS (–E64) and compared to a control in which the enzyme was inhibited by E64 (+E64).

Ac-AE(EDANS)FCitTTTSQGGK(Dabcyl)

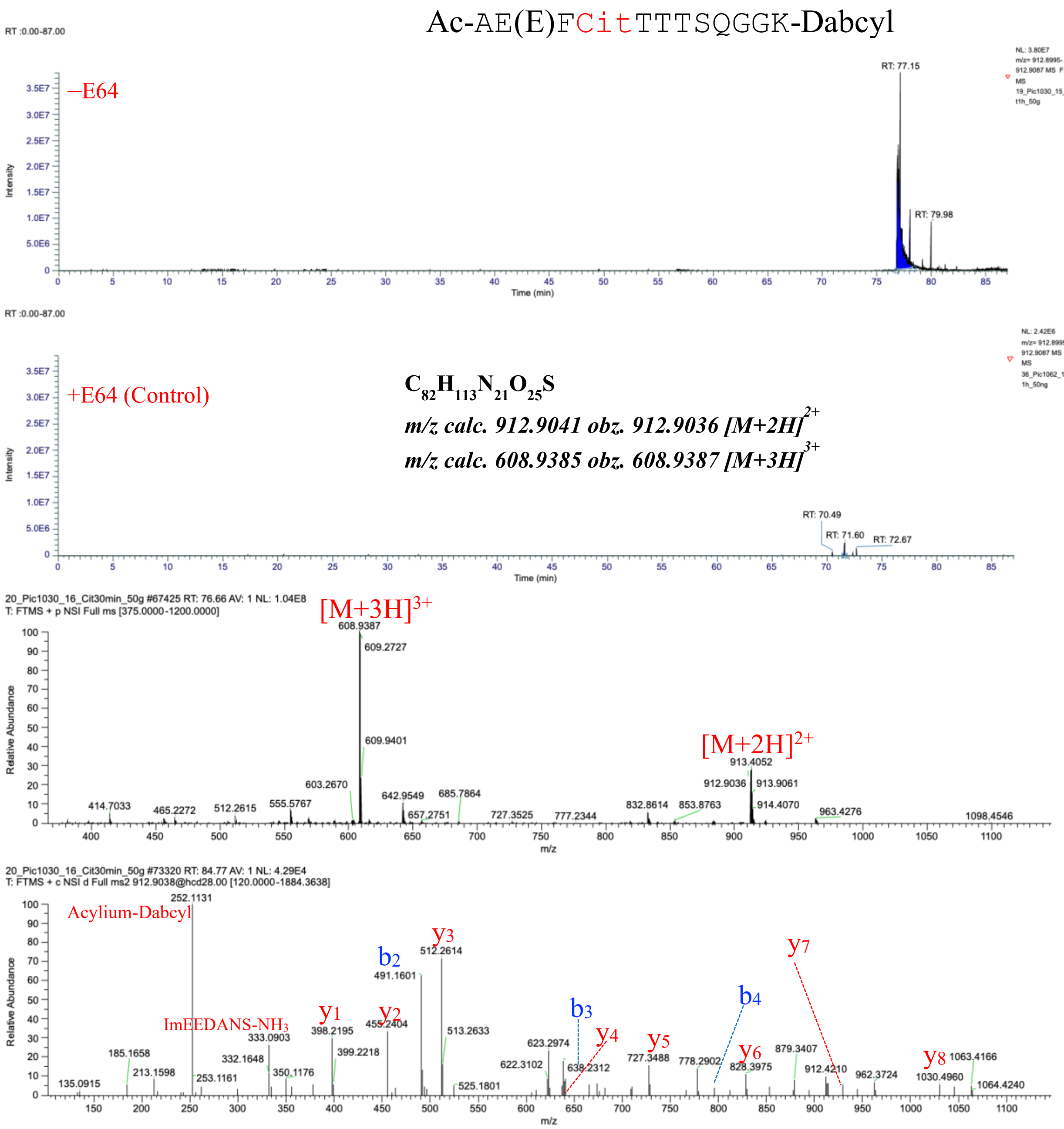

**Fig. S16.** Ac-AE(EDANS)FCitTTTSQGGK(Dabcyl) characterized by LC-MS/MS as a product of hCatS-catalyzed transpeptidation reaction. The figure displays a graphical representation of the extracted ion chromatogram (EIC), mass spectrum, and tandem mass spectrum of Ac-AE(E)FCitTTTSQGGK(Dabcyl), identified from a reaction between Ac-AE(E)FCitTTSQGGK(D) and TTSQGGK(Biotin) in the presence of active hCatS. LC-MS/MS analysis was performed on samples treated with active hCatS (–E64) and compared to a control in which the enzyme was inhibited by E64 (+E64).

#### Ac-AE(EDANS)FCitTTSQGGK(Biotin)

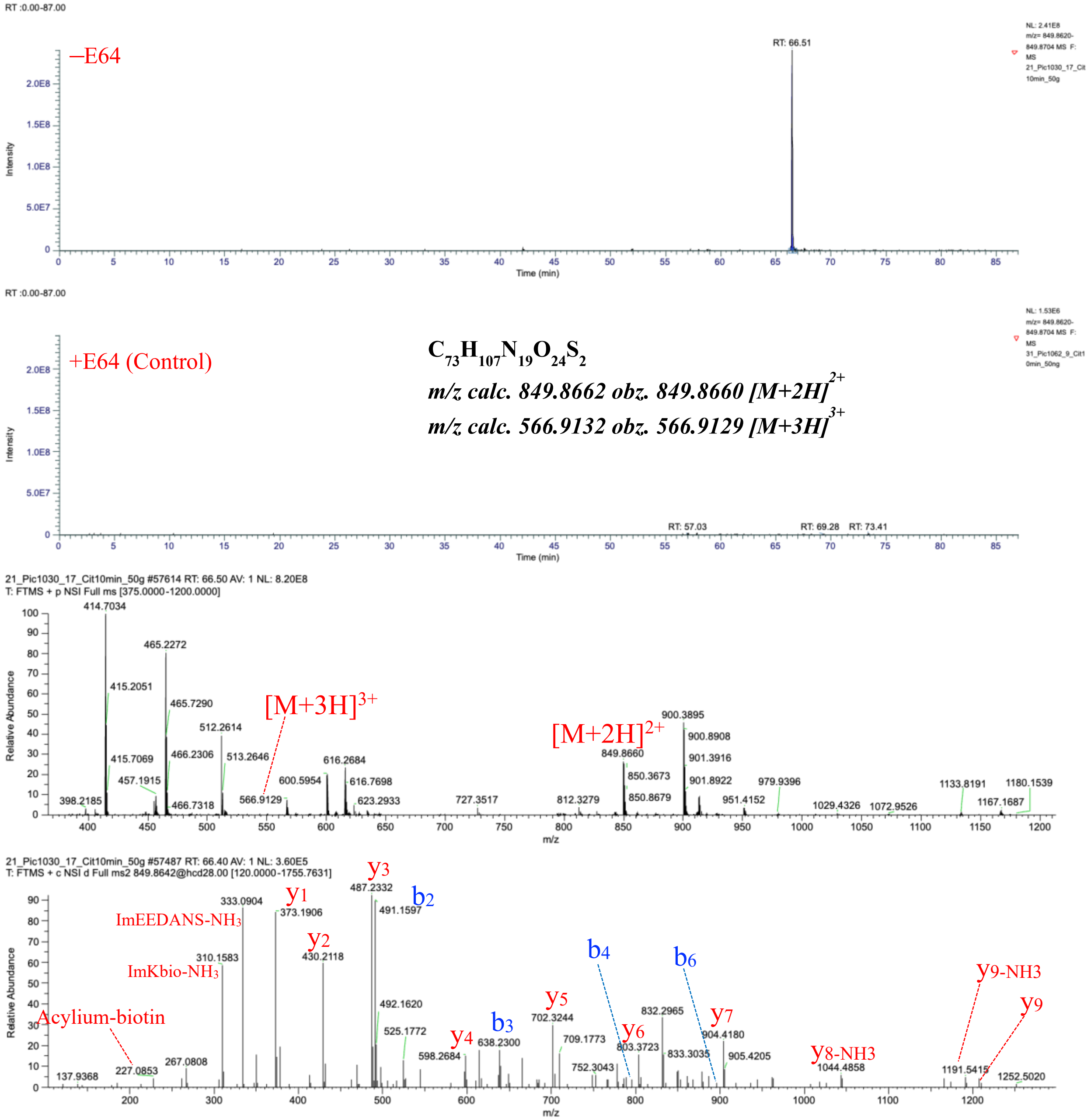

**Fig. S17.** Ac-AE(EDANS)FCitTTSQGGK(Biotin) characterized by LC-MS/MS as a product of hCatS-catalyzed transpeptidation reaction. The figure displays a graphical representation of the extracted ion chromatogram (EIC), mass spectrum, and tandem mass spectrum of Ac-AE(E)FCitTTSQGGK(Biotin), identified from a reaction between Ac-AE(E)FCitTTSQGGK(D) and TTSQGGK(Biotin) in the presence of active hCatS. LC-MS/MS analysis was performed on samples treated with active hCatS (–E64) and compared to a control in which the enzyme was inhibited by E64 (+E64).

Ac-AE(EDANS)FCitTTTTSQGGK(Biotin)

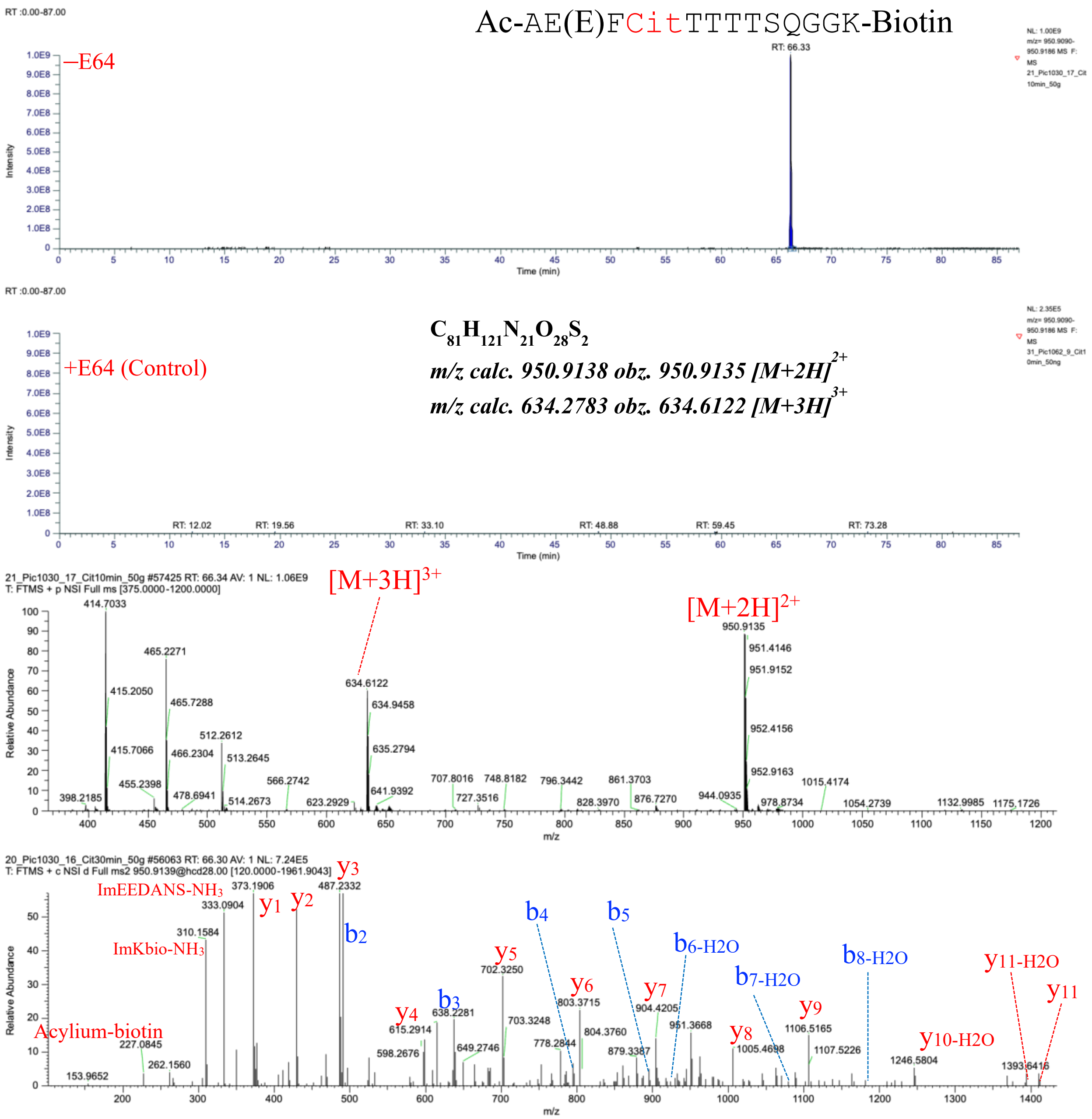

**Fig. S18.** Ac-AE(EDANS)FCitTTTTSQGGK(Biotin) characterized by LC-MS/MS as a product of hCatS-catalyzed transpeptidation reaction. The figure displays a graphical representation of the extracted ion chromatogram (EIC), mass spectrum, and tandem mass spectrum of Ac-AE(E)FCitTTTTSQGGK(Biotin), identified from a reaction between Ac-AE(E)FCitTTSQGGK(D) and TTSQGGK(Biotin) in the presence of active hCatS. LC-MS/MS analysis was performed on samples treated with active hCatS (–E64) and compared to a control in which the enzyme was inhibited by E64 (+E64).

Ac-AE(EDANS)FCitTTTTTSQGGK(Biotin)

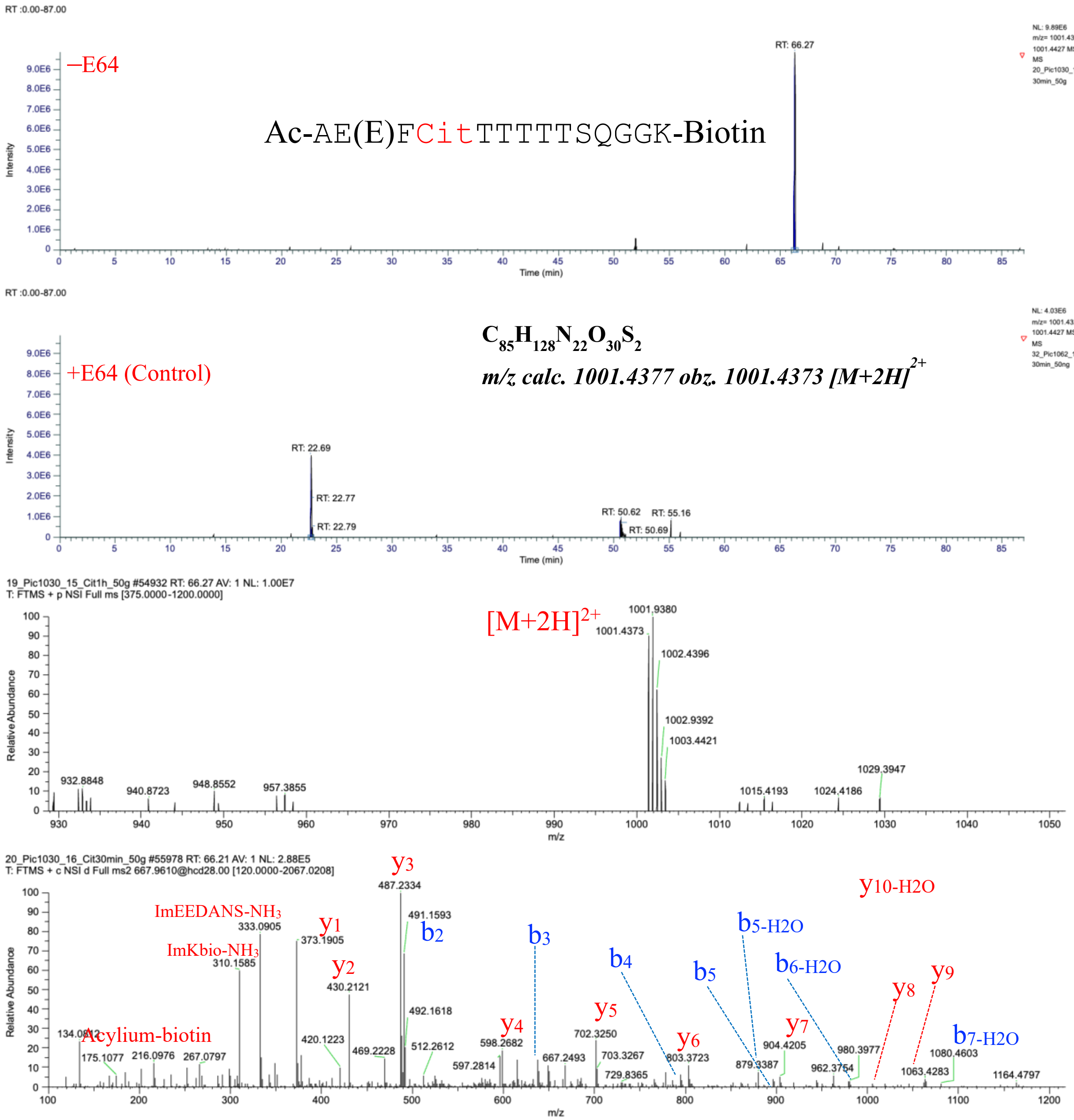

**Fig. S19.** Ac-AE(EDANS)FCitTTTTTSQGGK(Biotin) characterized by LC-MS/MS as a product of hCatS-catalyzed transpeptidation reaction. The figure displays a graphical representation of the extracted ion chromatogram (EIC), mass spectrum, and tandem mass spectrum of Ac-AE(E)FCitTTTTTSQGGK(Biotin), identified from a reaction between Ac-AE(E)FCitTTSQGGK(D) and TTSQGGK(Biotin) in the presence of active hCatS. LC-MS/MS analysis was performed on samples treated with active hCatS (–E64) and compared to a control in which the enzyme was inhibited by E64 (+E64).

Ac-AE(EDANS)FCitTTTSQGGK(Biotin)

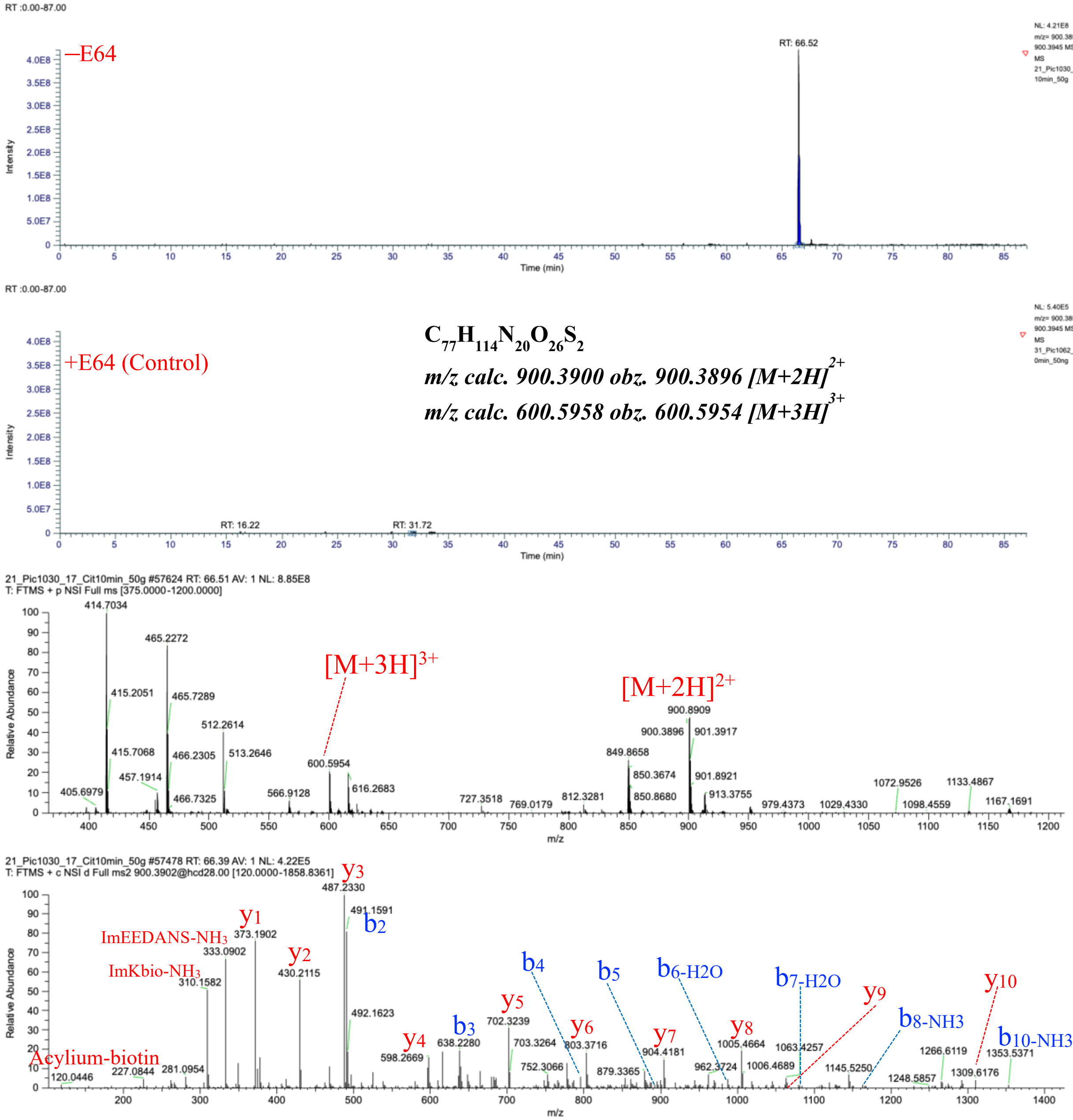

**Fig. S20.** Ac-AE(EDANS)FCitTTTSQGGK(Biotin) characterized by LC-MS/MS as a product of hCatS-catalyzed transpeptidation reaction. The figure displays a graphical representation of the extracted ion chromatogram (EIC), mass spectrum, and tandem mass spectrum of Ac-AE(E)FCitTTTSQGGK(Biotin), identified from a reaction between Ac-AE(E)FCitTTSQGGK(D) and TTSQGGK(Biotin) in the presence of active hCatS. LC-MS/MS analysis was performed on samples treated with active hCatS (–E64) and compared to a control in which the enzyme was inhibited by E64 (+E64).

#### Comparing the peak area and rate of formation of cispeptides (citrullinated vs non-citrullinated)

A

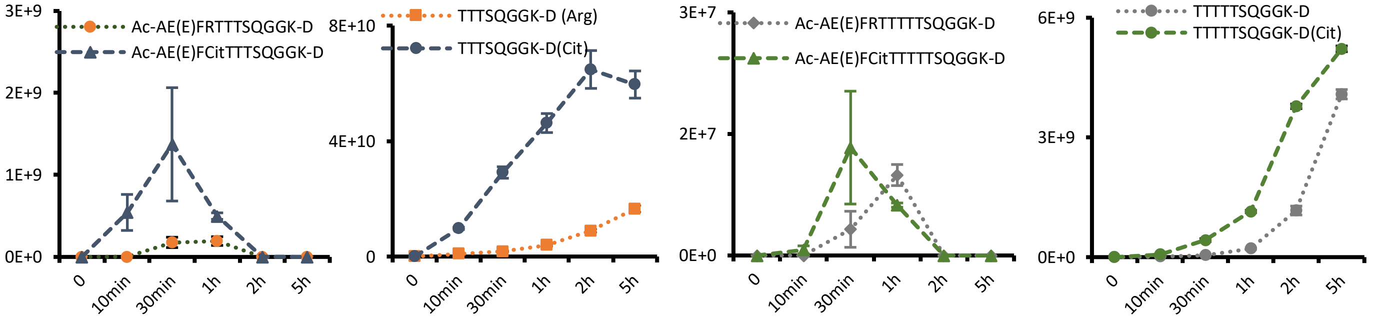

B

| Parent peptide | Rate (PA/min) | Hydrolysis Product | Rate (PA/min) |
|---|---|---|---|
| Ac-AE(EDANS)FRTTSQGGK-Biotin | 12.7 ± 1 (x10$^{8}$) | TTSQGGK-Biotin | N/A |
| Ac-AE(EDANS)FCitTTSQGGK-Biotin | 10.9 ± 1.3 (x10$^{8}$) | TTSQGGK-Biotin | N/A |
| Ac-AE(EDANS)FRTTTSQGGK-Biotin | 2.3 ± 0.1 (x10$^{8}$) | TTTSQGGK-Biotin | 0.9 ± 0.2 (x10$^{8}$) |
| Ac-AE(EDANS)FCitTTTSQGGK-Biotin | 22.4 ± 4 (x10$^{8}$) | TTTSQGGK-Biotin | 19.4 ± 1.5 (x10$^{8}$) |
| Ac-AE(EDANS)FRTTTTSQGGK-Biotin | 7.3 ± 0.8 (x10$^{9}$) | TTTTSQGGK-Biotin | 1.5 ± 0.4 (x10$^{9}$) |
| Ac-AE(EDANS)FCitTTTTSQGGK-Biotin | 5.2 ± 0.6 (x10$^{9}$) | TTTTSQGGK-Biotin | 2.7 ± 0.3 (x10$^{9}$) |
| Ac-AE(EDANS)FRTTTTTSQGGK-Biotin | 2.6 ± 0.5 (x10$^{5}$) | TTTTTSQGGK-Biotin | 5.2 ± 0.4 (x10$^{6}$) |
| Ac-AE(EDANS)FCitTTTTTSQGGK-Biotin | 7.2 ± 0.6 (x10$^{5}$) | TTTTTSQGGK-Biotin | 17.2 ± 0.3 (x10$^{6}$) |
| Ac-AE(EDANS)FRTTTSQGGK-Dabcyl | 59 ± 21 (x10$^{5}$) | TTTSQGGK-Dabcyl | 65 ± 6 (x10$^{7}$) |
| Ac-AE(EDANS)FCitTTTSQGGK-Dabcyl | 457 ± 230 (x10$^{5}$) | TTTSQGGK-Dabcyl | 770 ± 54 (x10$^{7}$) |
| Ac-AE(EDANS)FRTTTTSQGGK-Dabcyl | 8.6 ± 1.6 (x10$^{7}$) | TTTTSQGGK-Dabcyl | 5.1 ± 0.1 (x10$^{8}$) |
| Ac-AE(EDANS)FCitTTTTSQGGK-Dabcyl | 2.4.4 ± 0.8 (x10$^{7}$) | TTTTSQGGK-Dabcyl | 2.1 ± 0.1 (x10$^{8}$) |
| Ac-AE(EDANS)FRTTTTTSQGGK-Dabcyl | 10.7 ± 7.4 (x10$^{4}$) | TTTTTSQGGK-Dabcyl | 9.7 ± 0.9 (x10$^{6}$) |
| Ac-AE(EDANS)FCitTTTTTSQGGK-Dabcyl | 44.3 ± 23 (x10$^{4}$) | TTTTTSQGGK-Dabcyl | 31.4 ± 0.4 (x10$^{6}$) |

**Fig S21:** (A) Peak areas of cispeptides with corresponding hydrolysis products generated by hCatS in reactions with Ac-AE(E)FRTTSQGGK(D) or Ac-AE(E)FCitTTSQGGK(D) as substrate.(B) summarizes the formation and hydrolysis rates of all identified cis and trans fusion and follow-up hydrolysis products from the Arg and Cit-containing FRET based substrates.

#### Characterization of Insulin derived biotinylated (NAVEGGK-Biotin conjugates) transpeptides

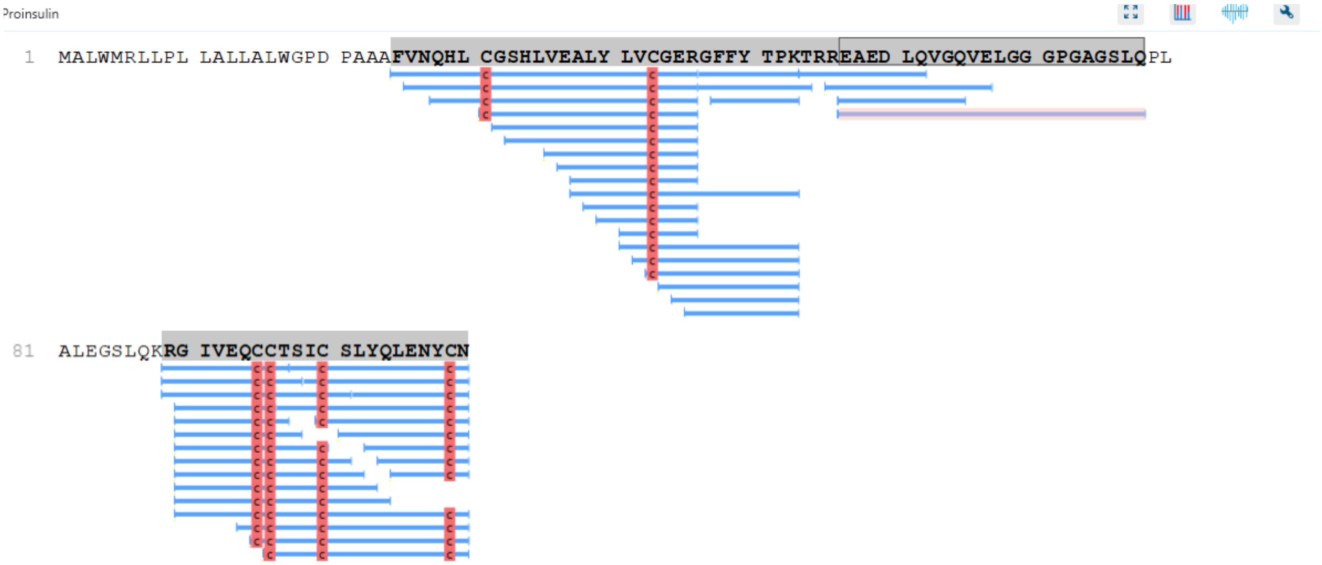

**Fig S22**: Sequence coverage of human proinsulin from LC–MS/MS analysis of trypsin-digested mature insulin using PEAKS Studio. Mature insulin (**Recombinant human insulin (Roche, Germany; Cat. No. 11376497001, yeast-derived)** was digested with trypsin and analyzed by LC–MS/MS, and MS/MS spectra were searched in PEAKS Studio using the human proinsulin sequence as the database with digestion mode set to Auto, reflecting parallel cathepsin S digestion experiments. The full proinsulin sequence is shown, with identified peptides underlined in blue. In addition to the expected A and B chain peptides, multiple peptides mapping to the C peptide region were detected, indicating contamination of the insulin preparation with residual proinsulin or partially processed intermediates.

Comparing chemically synthesized and enzyme catalyzed ELGGGNAVEGGK-Biotin

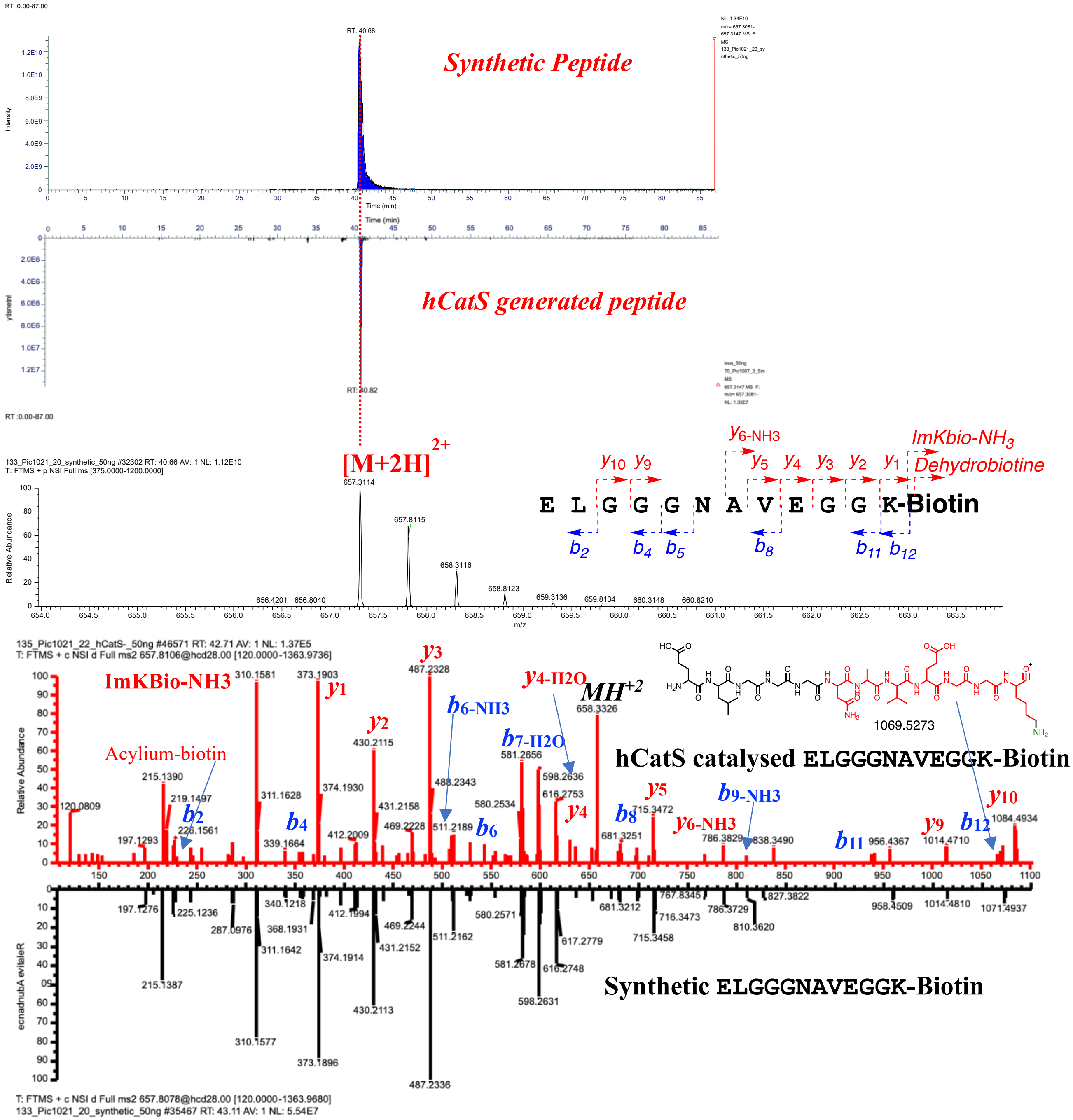

**Fig. S23.** The figure presents a comparison of the graphical representations of the extracted ion chromatogram (EIC), mass spectrum, and tandem mass spectrum between synthetic ELGGGNAVEGGK(Biotin) and the enzymatically produced counterpart.

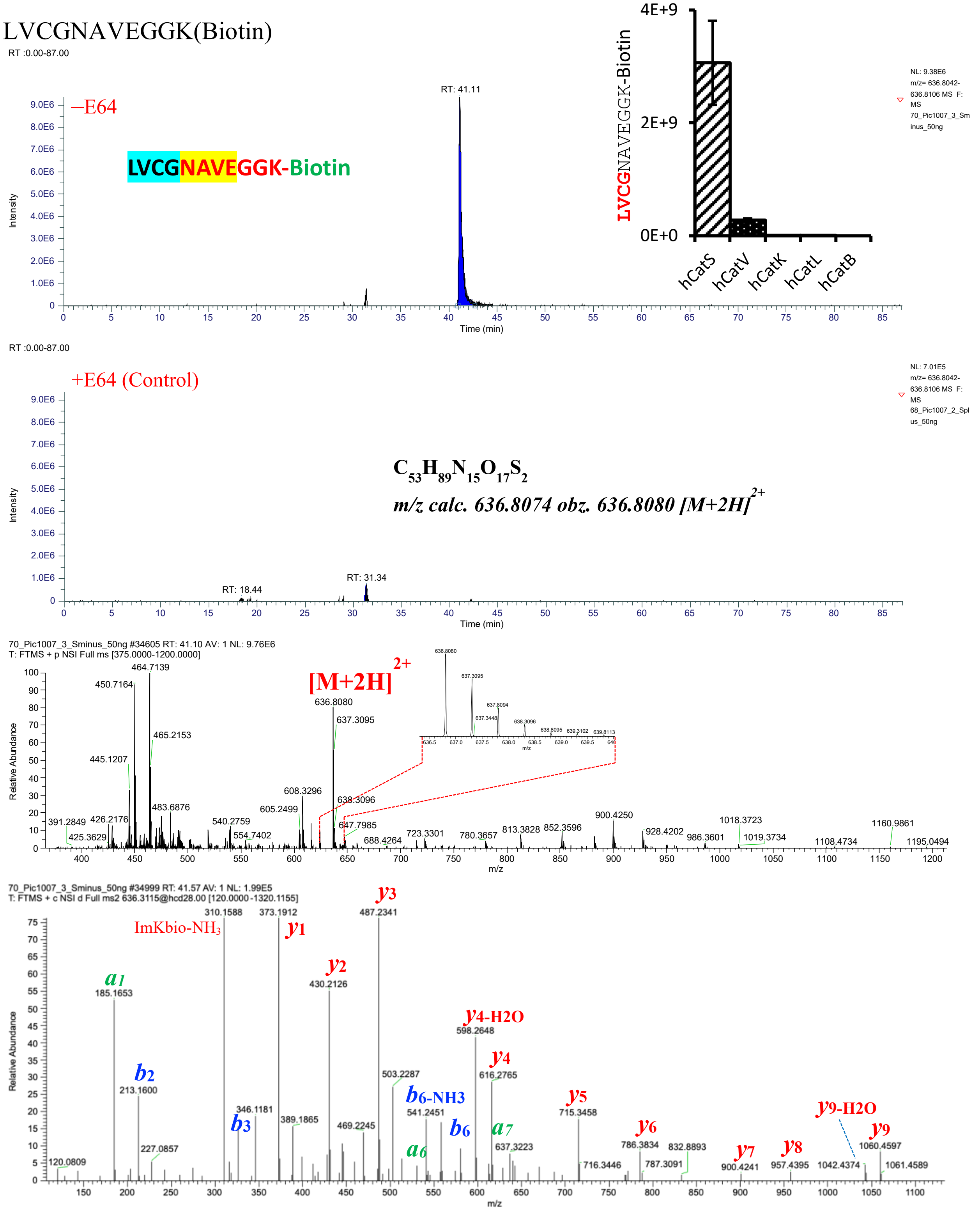

**Fig. S24.** LVCGNAVEGGK(Biotin) characterized by LC-MS/MS as a product of hCatS-catalyzed transpeptidation reaction. The figure displays a graphical representation of the extracted ion chromatogram (EIC), mass spectrum, and tandem mass spectrum of LVCGNAVEGGK(Biotin), identified from a reaction between NAVEGGK(Biotin) and Human Proinsulin in the presence of active hCatS.

#### ALYLNAVEGGK(Biotin)

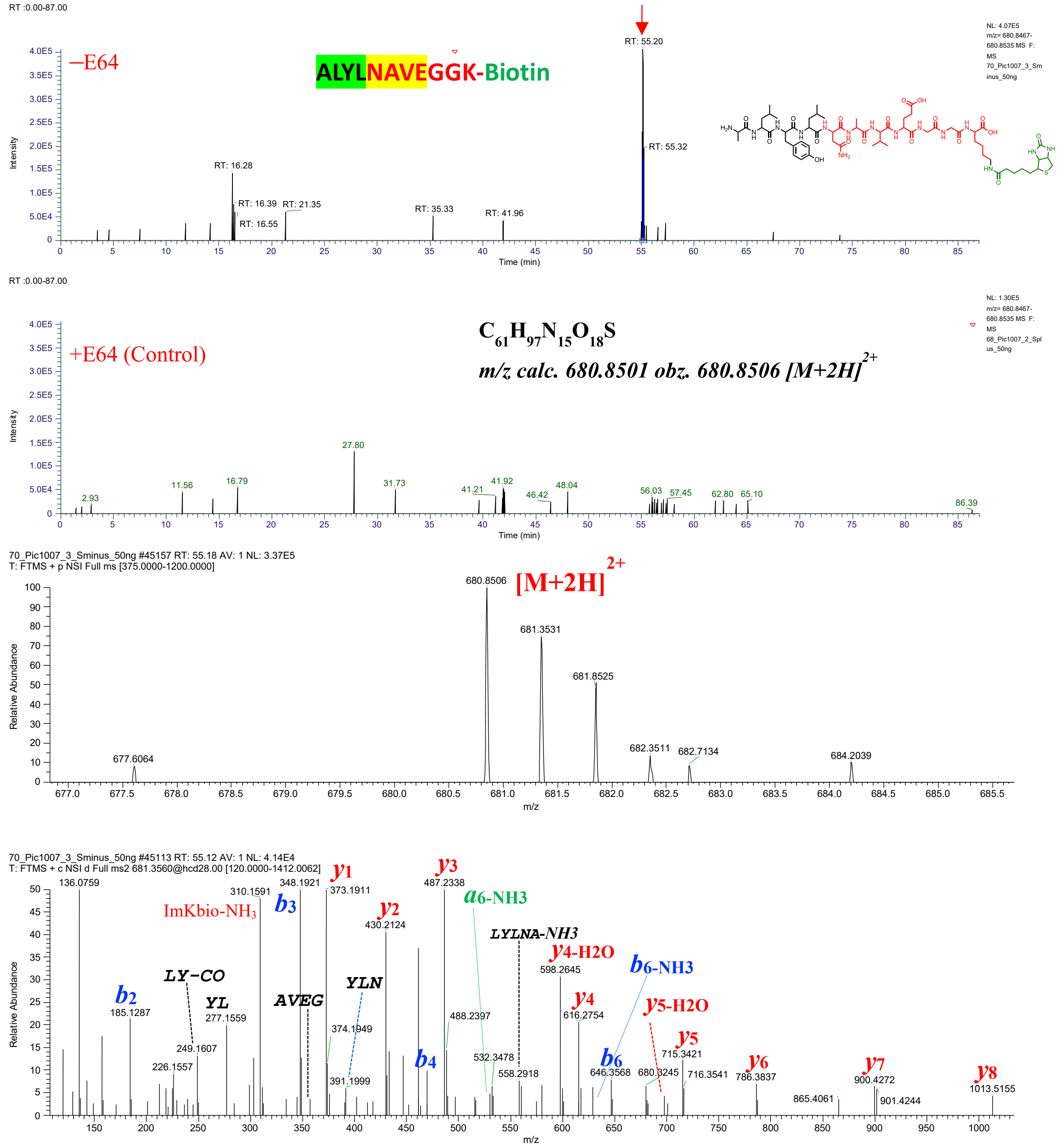

**Fig. S25.** ALYLNAVEGGK(Biotin) characterized by LC-MS/MS as a product of hCatS-catalyzed transpeptidation reaction. The figure displays a graphical representation of the extracted ion chromatogram (EIC), mass spectrum, and tandem mass spectrum of ALYLNAVEGGK(Biotin), identified from a reaction between NAVEGGK(Biotin) and Human Proinsulin in the presence of active hCatS. LC-MS/MS analysis was performed on samples treated with active hCatS (–E64) and compared to a control in which the enzyme was inhibited by E64 (+E64).

FVNQHLCGNAVEGGK(Biotin)

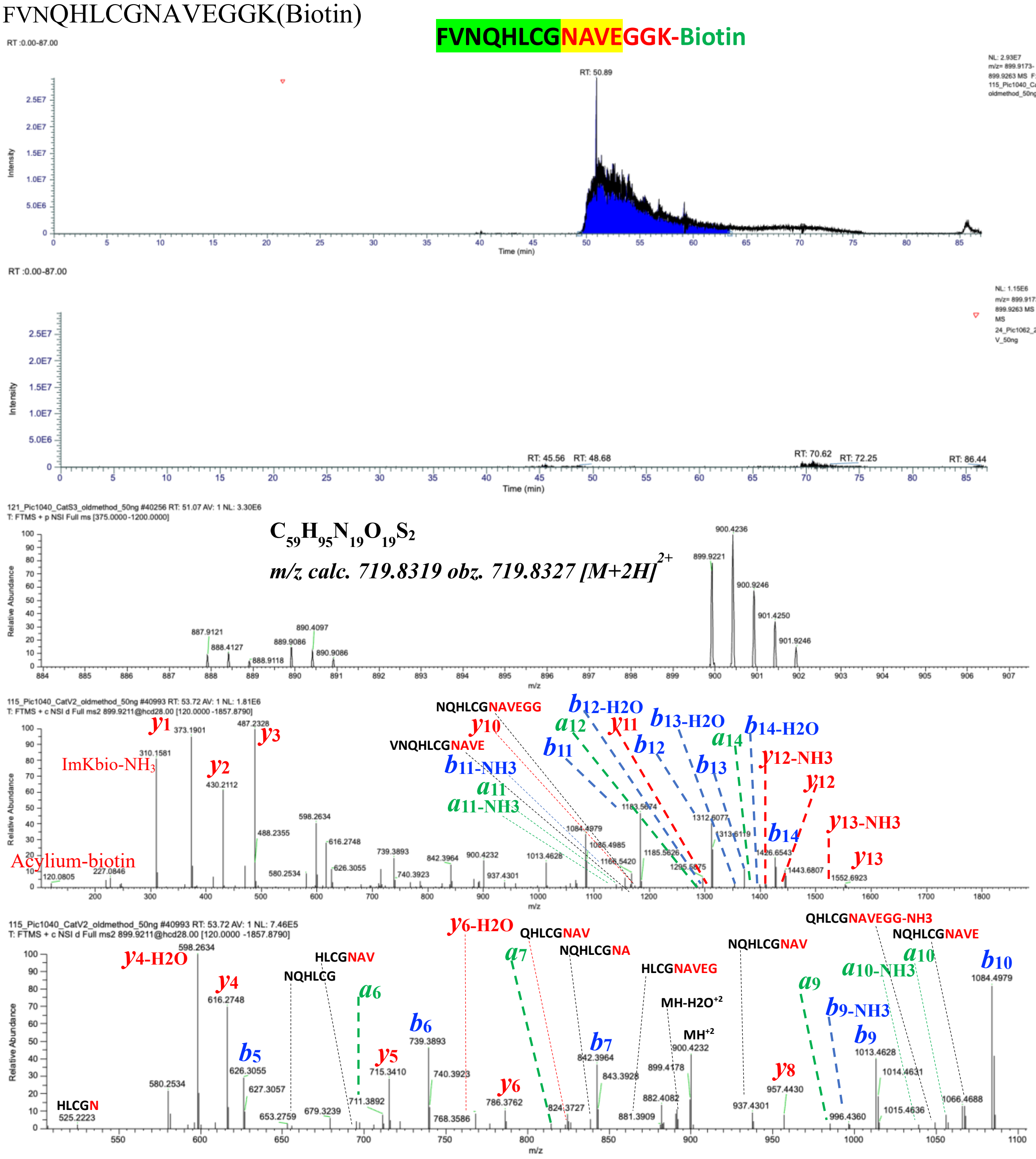

**Fig. S26.** FVNQHLCGNAVEGGK(Biotin) characterized by LC-MS/MS as a product of hCatS-catalyzed transpeptidation reaction. The figure displays a graphical representation of the extracted ion chromatogram (EIC), mass spectrum, and tandem mass spectrum of FVNQHLCGNAVEGGK(Biotin), identified from a reaction between NAVEGGK(Biotin) and Human Proinsulin in the presence of active hCatS. LC-MS/MS analysis was performed on samples treated with active hCatS (–E64) and compared to a control in which the enzyme was inhibited by E64 (+E64). The molecular ion was extracted based on its exact monoisotopic mass, and major fragment ions in the tandem mass spectrum were assigned

GIVEQCCTNAVEGGK(Biotin)

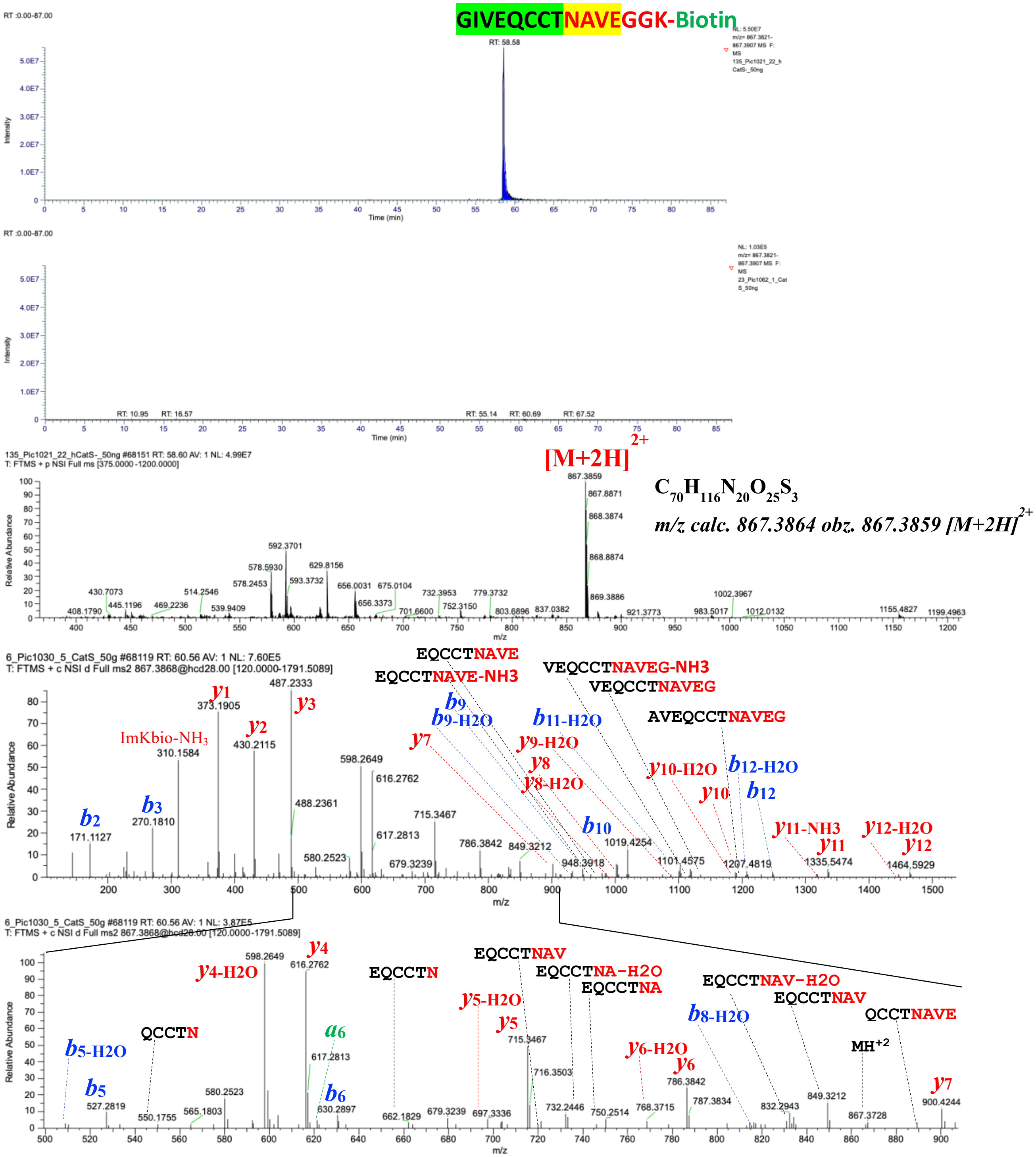

**Fig. S27.** GIVEQCCTNAVEGGK(Biotin) characterized by LC-MS/MS as a product of hCatS-catalyzed transpeptidation reaction. The figure displays a graphical representation of the extracted ion chromatogram (EIC), mass spectrum, and tandem mass spectrum of GIVEQCCTNAVEGGK(Biotin), identified from a reaction between NAVEGGK(Biotin) and Human Proinsulin in the presence of active hCatS.

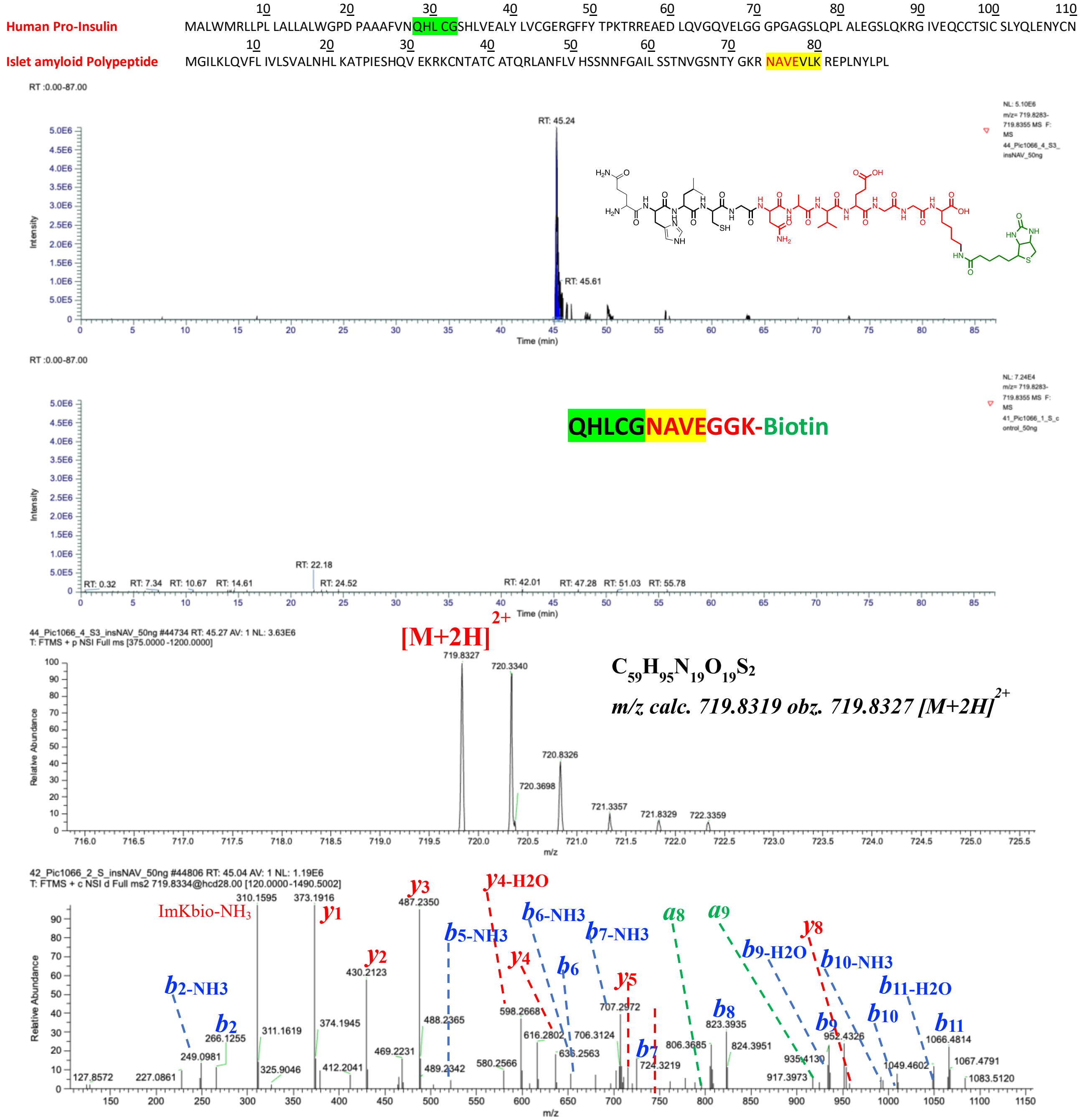

**Fig. S28.** QHLCGNAVEGGK(Biotin) characterized by LC-MS/MS as a product of hCatS-catalyzed transpeptidation reaction. The figure displays a graphical representation of the extracted ion chromatogram (EIC), mass spectrum, and tandem mass spectrum of QHLCGNAVEGGK(Biotin), identified from a reaction between NAVEGGK(Biotin) and Human Proinsulin in the presence of active hCatS.

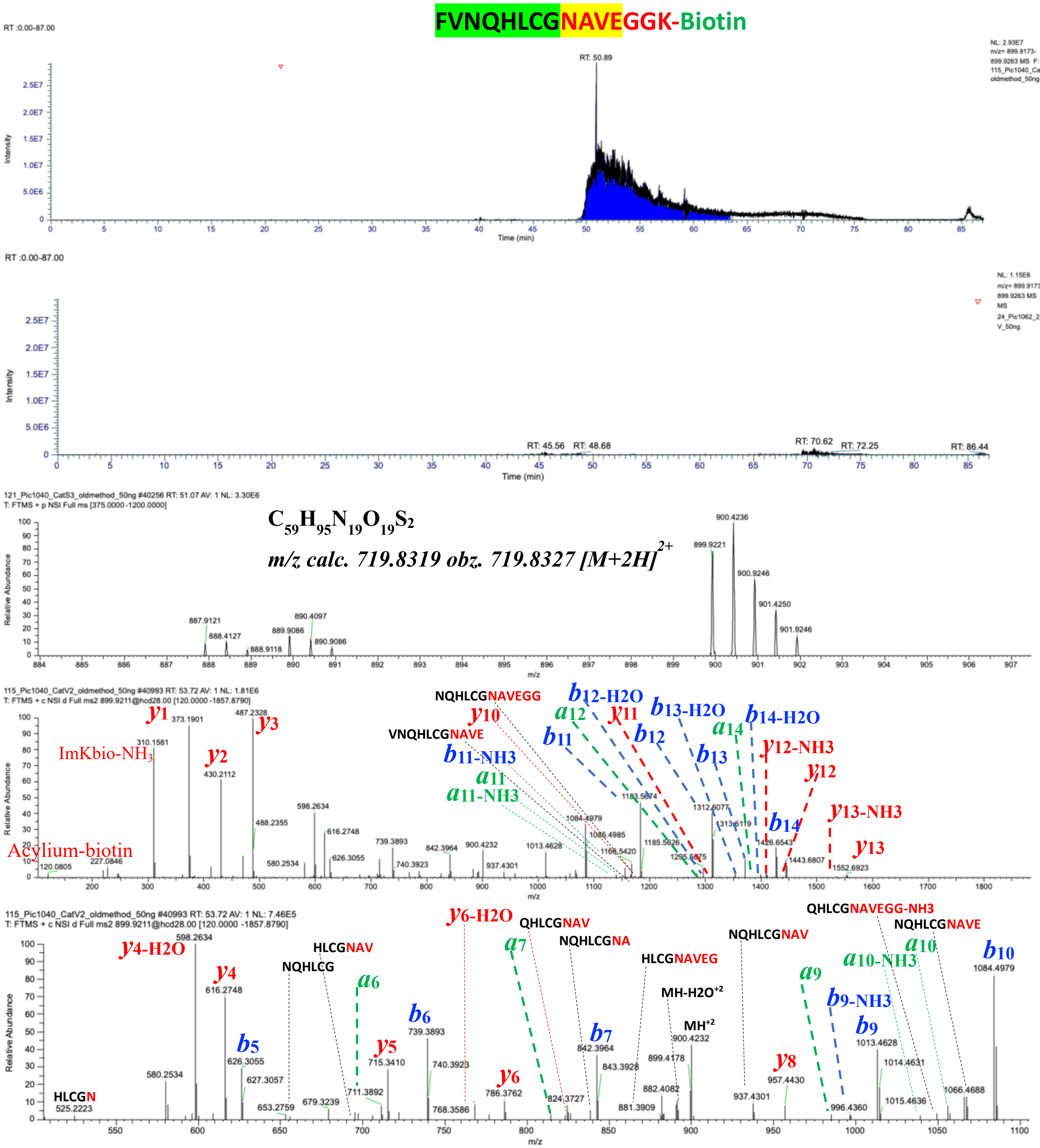

**Fig. S29.** FVNQHLCGNAVEGGK(Biotin) characterized by LC-MS/MS as a product of hCatS-catalyzed transpeptidation reaction. The figure displays a graphical representation of the extracted ion chromatogram (EIC), mass spectrum, and tandem mass spectrum of FVNQHLCGNAVEGGK(Biotin), identified from a reaction between NAVEGGK(Biotin) and Human Proinsulin in the presence of active hCatS.

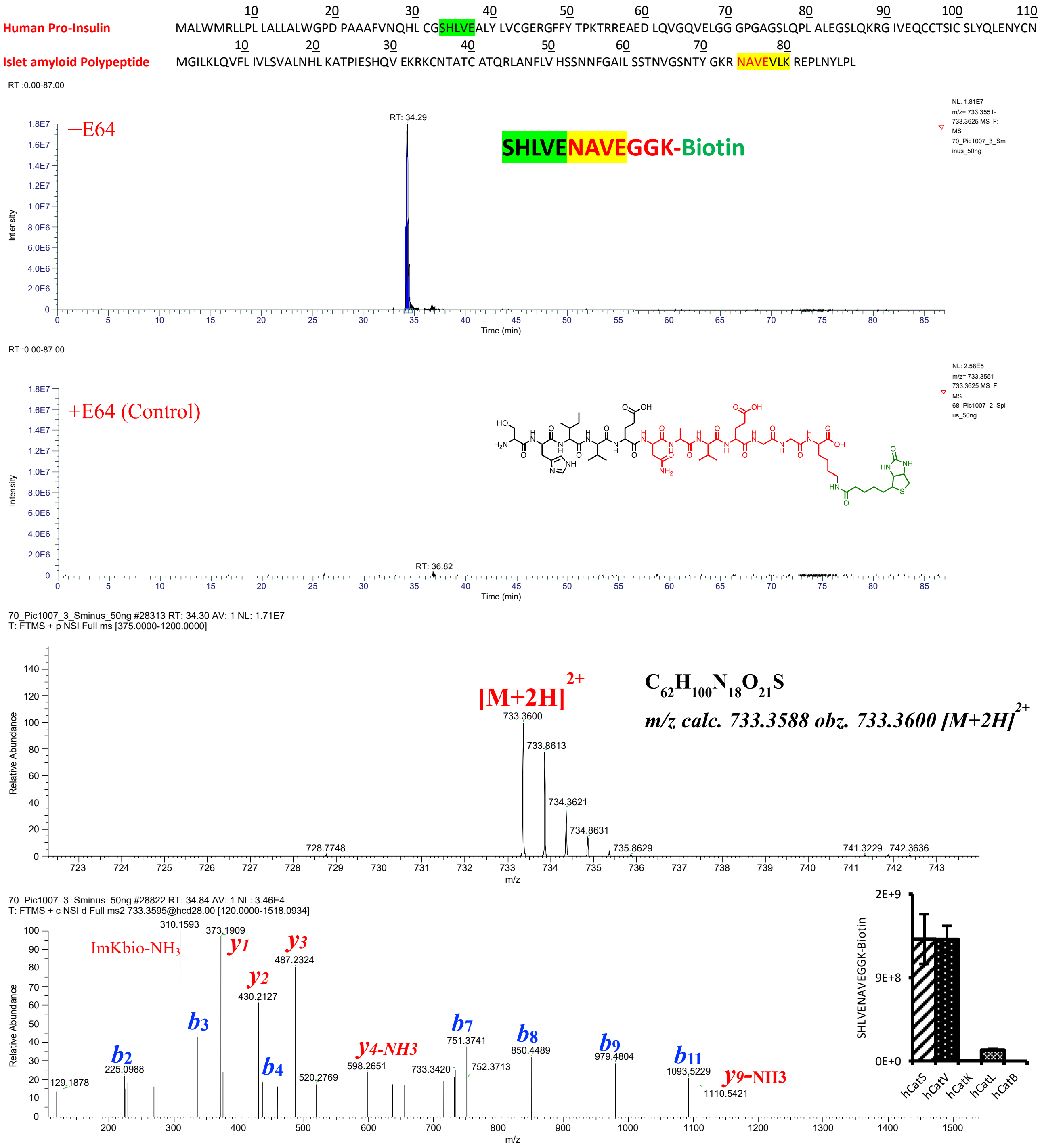

**Fig. S30.** SHLVENAVEGGK(Biotin) characterized by LC-MS/MS as a product of hCatS-catalyzed transpeptidation reaction. The figure displays a graphical representation of the extracted ion chromatogram (EIC), mass spectrum, and tandem mass spectrum of SHLVENAVEGGK(Biotin), identified from a reaction between NAVEGGK(Biotin) and Human Proinsulin in the presence of active hCatS.

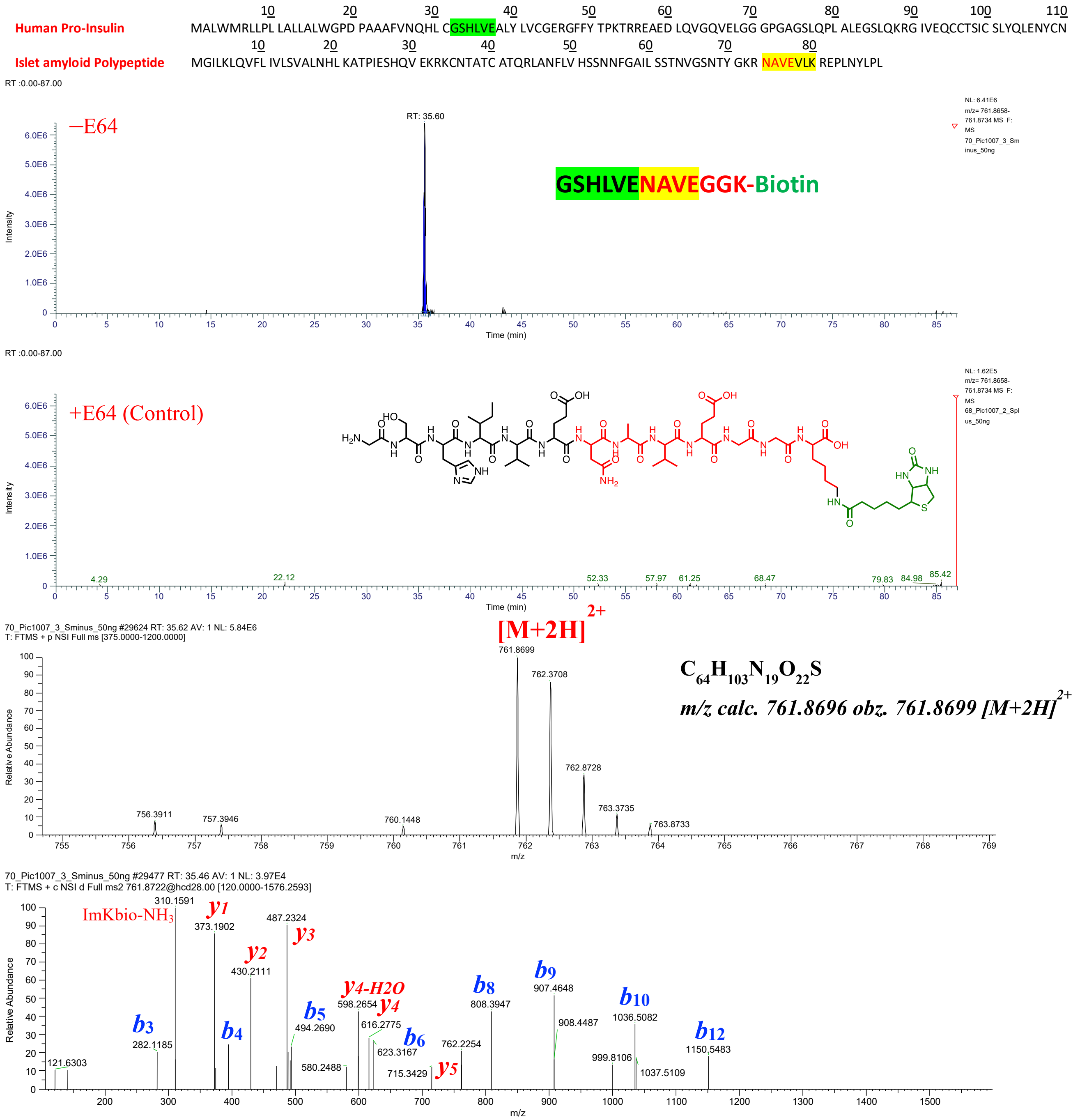

**Fig. S31.** GSHLVENAVEGGK(Biotin) characterized by LC-MS/MS as a product of hCatS-catalyzed transpeptidation reaction. The figure displays a graphical representation of the extracted ion chromatogram (EIC), mass spectrum, and tandem mass spectrum of GSHLVENAVEGGK(Biotin), identified from a reaction between NAVEGGK(Biotin) and Human Proinsulin in the presence of active hCatS.

| | |
|---|---|
| Human Pro-Insulin | MALWMRLLPL LALLALWGPD PAAAFVNQHL CGSHLVEALY LVCGERGFFY TPKTRREAED LQVGQVELGG GPGAGSLQPL ALEGSLQKRG IVEQCCTSIC SLYQLENYCN |
| Islet amyloid Polypeptide | MGILKLQVFL IVLSVALNHL KATPIESHQV EKRKCNTATC ATQRLANFLV HSSNNFGAIL SSTNVGSNTY GKR NAVEVLK REPLNYLPL |

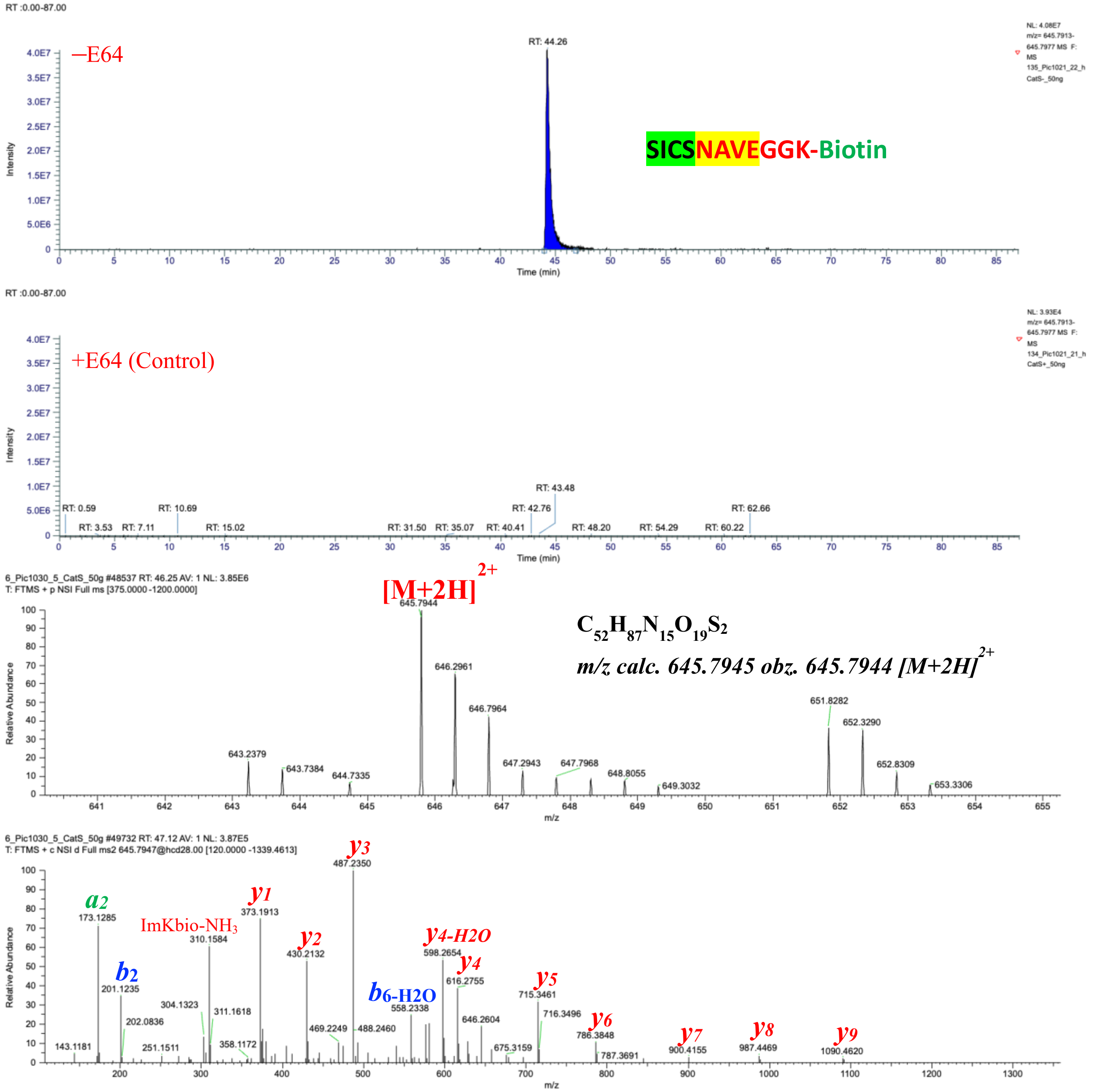

**Fig. S32.** SICSNAVEGGK(Biotin) characterized by LC-MS/MS as a product of hCatS-catalyzed transpeptidation reaction. The figure displays a graphical representation of the extracted ion chromatogram (EIC), mass spectrum, and tandem mass spectrum of SICSNAVEGGK(Biotin), identified from a reaction between NAVEGGK(Biotin) and Human Proinsulin in the presence of active hCatS.

#### Characterization of Spike derived biotinylated (NAVEGGK-Biotin conjugates) transpeptides

NGTITNAVEGGK(Biotin)

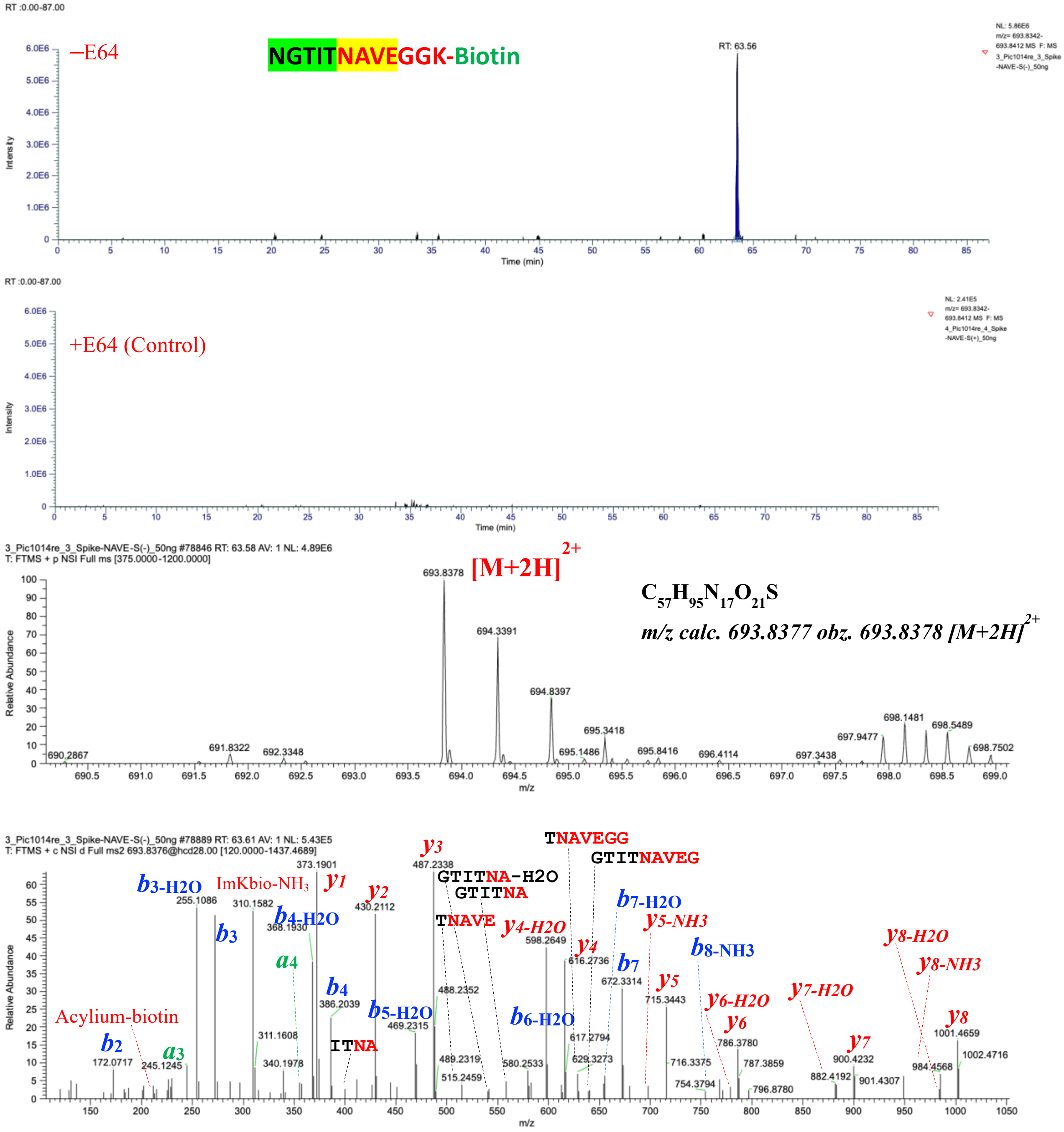

**Fig. S33.** NGTITNAVEGGK(Biotin) characterized by LC-MS/MS as a product of hCatS-catalyzed transpeptidation reaction. The figure displays a graphical representation of the extracted ion chromatogram (EIC), mass spectrum, and tandem mass spectrum of NGTITNAVEGGK(Biotin), identified from a reaction between NAVEGGK(Biotin) and Spike of SARS-CoV-2 in the presence of active hCatS. LC-MS/MS analysis was performed on samples treated with active hCatS (–E64) and compared to a control in which the enzyme was inhibited by E64 (+E64). The molecular ion was extracted based on its exact monoisotopic mass, and major fragment ions in the tandem mass spectrum were assigned.was inhibited by E64 (+E64).

SGWTFGNAVEGGK(Biotin)

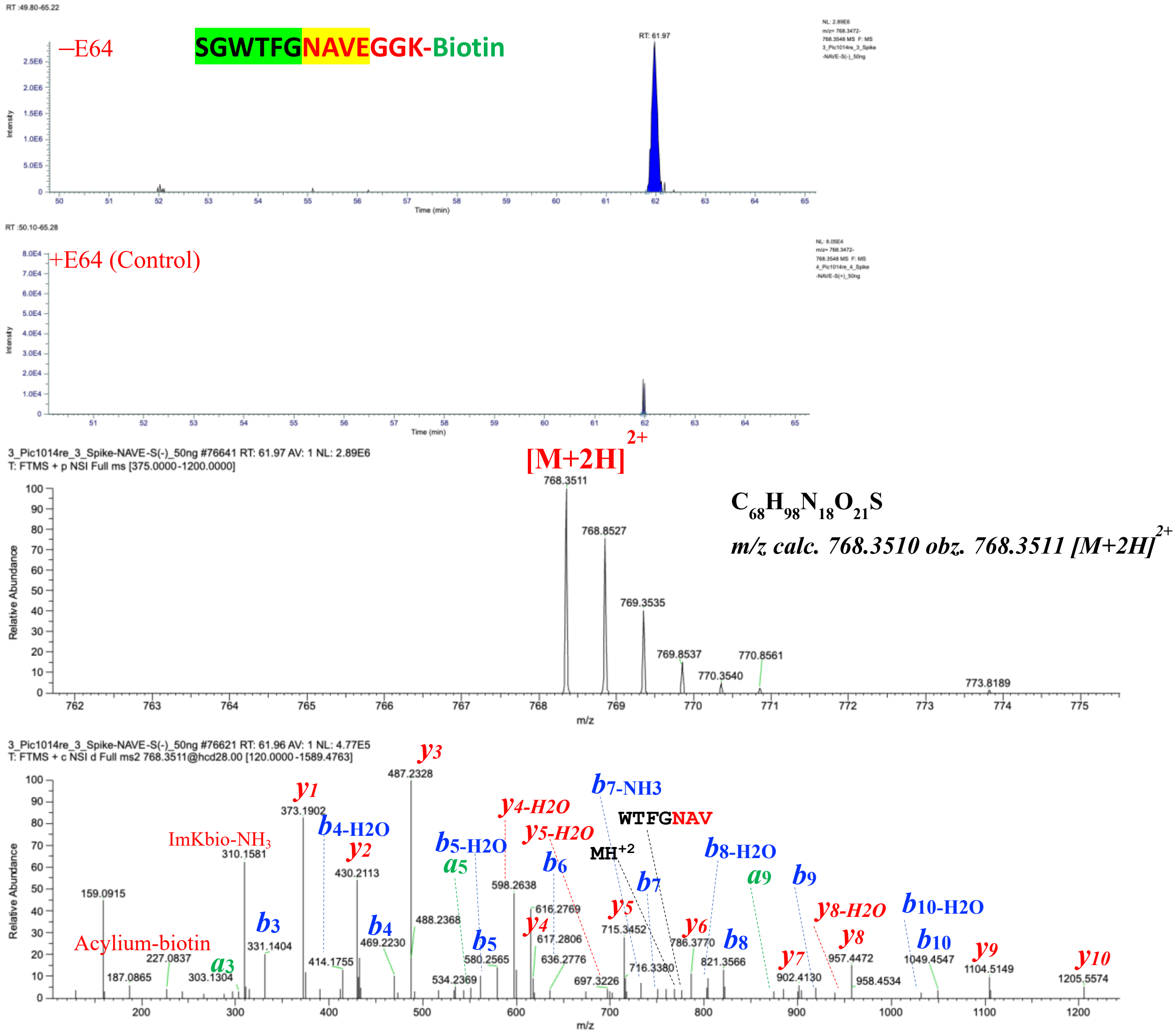

**Fig. S34.** SGWTFGNAVEGGK(Biotin) characterized by LC-MS/MS as a product of hCatS-catalyzed transpeptidation reaction. The figure displays a graphical representation of the extracted ion chromatogram (EIC), mass spectrum, and tandem mass spectrum of SGWTFGNAVEGGK(Biotin), identified from a reaction between NAVEGGK(Biotin) and Spike of SARS-CoV-2 in the presence of active hCatS. LC-MS/MS analysis was performed on samples treated with active hCatS (–E64) and compared to a control in which the enzyme was inhibited by E64 (+E64). The molecular ion was extracted based on its exact monoisotopic mass, and major fragment ions in the tandem mass spectrum were assigned.was inhibited by E64 (+E64).

SFPQSAPHGNAVEGGK(Biotin)

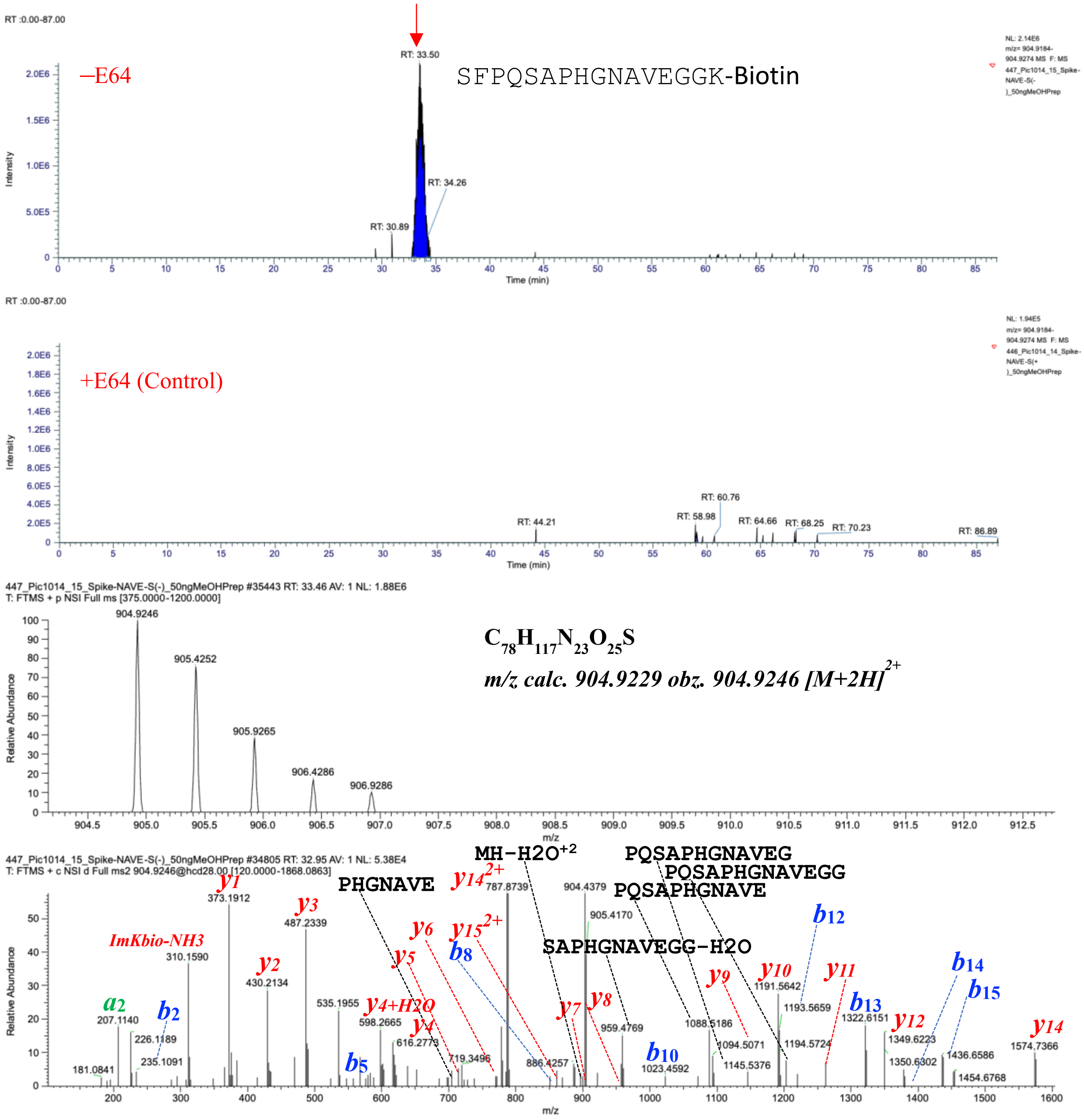

**Fig. S35.** SFPQSAPHGNAVEGGK(Biotin) characterized by LC-MS/MS as a product of hCatS-catalyzed transpeptidation reaction. The figure displays a graphical representation of the extracted ion chromatogram (EIC), mass spectrum, and tandem mass spectrum of SFPQSAPHGNAVEGGK(Biotin), identified from a reaction between NAVEGGK(Biotin) and Spike of SARS-CoV-2 in the presence of active hCatS. LC-MS/MS analysis was performed on samples treated with active hCatS (–E64) and compared to a control in which the enzyme was inhibited by E64 (+E64). The molecular ion was extracted based on its exact monoisotopic mass, and major fragment ions in the tandem mass spectrum were assigned.

LQYGNAVEGGK(Biotin)

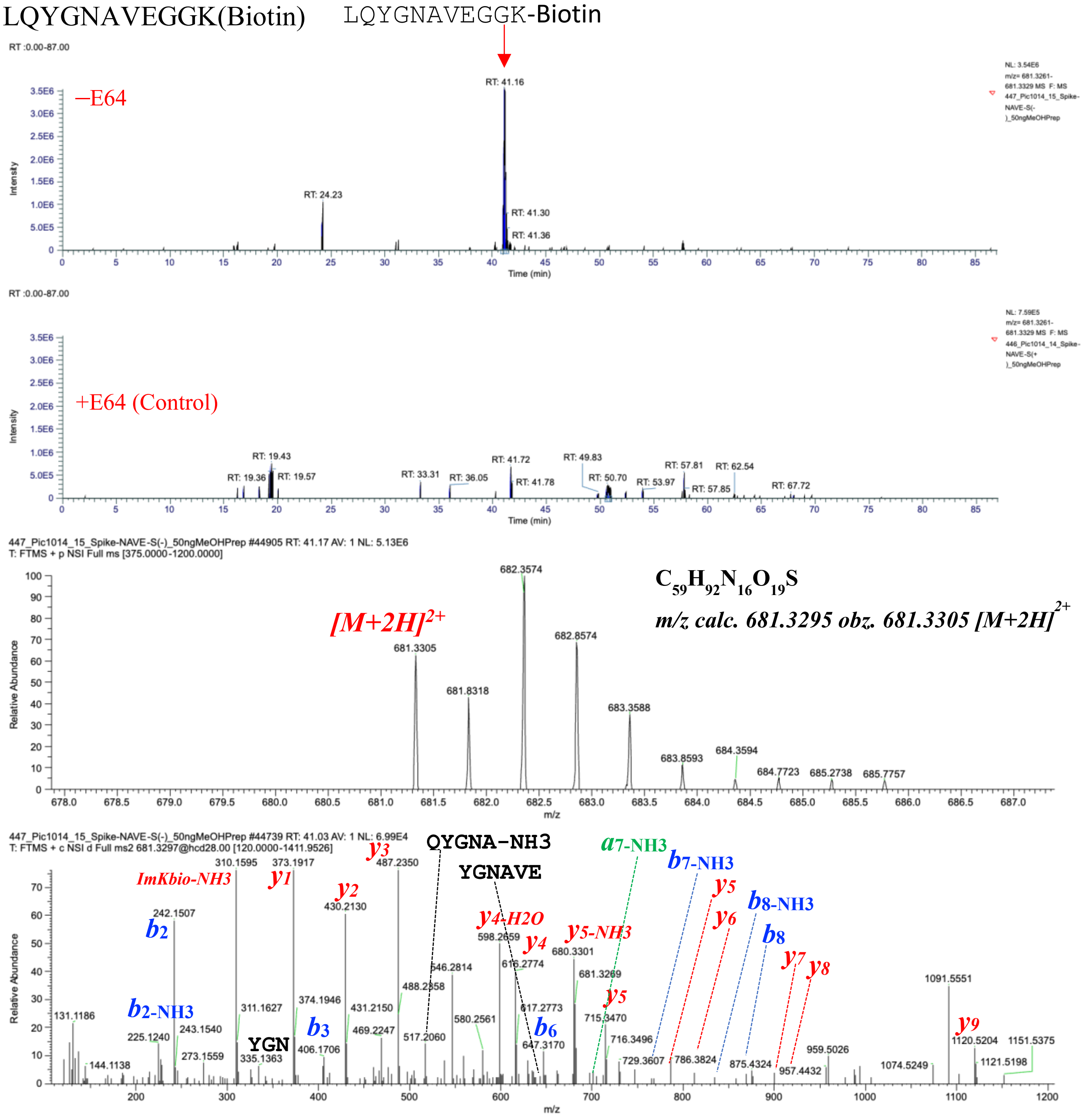

**Fig. S36.** LQYGNAVEGGK(Biotin) characterized by LC-MS/MS as a product of hCatS-catalyzed transpeptidation reaction. The figure displays a graphical representation of the extracted ion chromatogram (EIC), mass spectrum, and tandem mass spectrum of LQYGNAVEGGK(Biotin), identified from a reaction between NAVEGGK(Biotin) and Spike of SARS-CoV-2 in the presence of active hCatS. LC-MS/MS analysis was performed on samples treated with active hCatS (–E64) and compared to a control in which the enzyme was inhibited by E64 (+E64). The molecular ion was extracted based on its exact monoisotopic mass, and major fragment ions in the tandem mass spectrum were assigned.

#### Characterization of PF4 derived biotinylated (TTSQGGK-Biotin conjugates) transpeptides

SLEVIKTTSQGGK(Biotin)

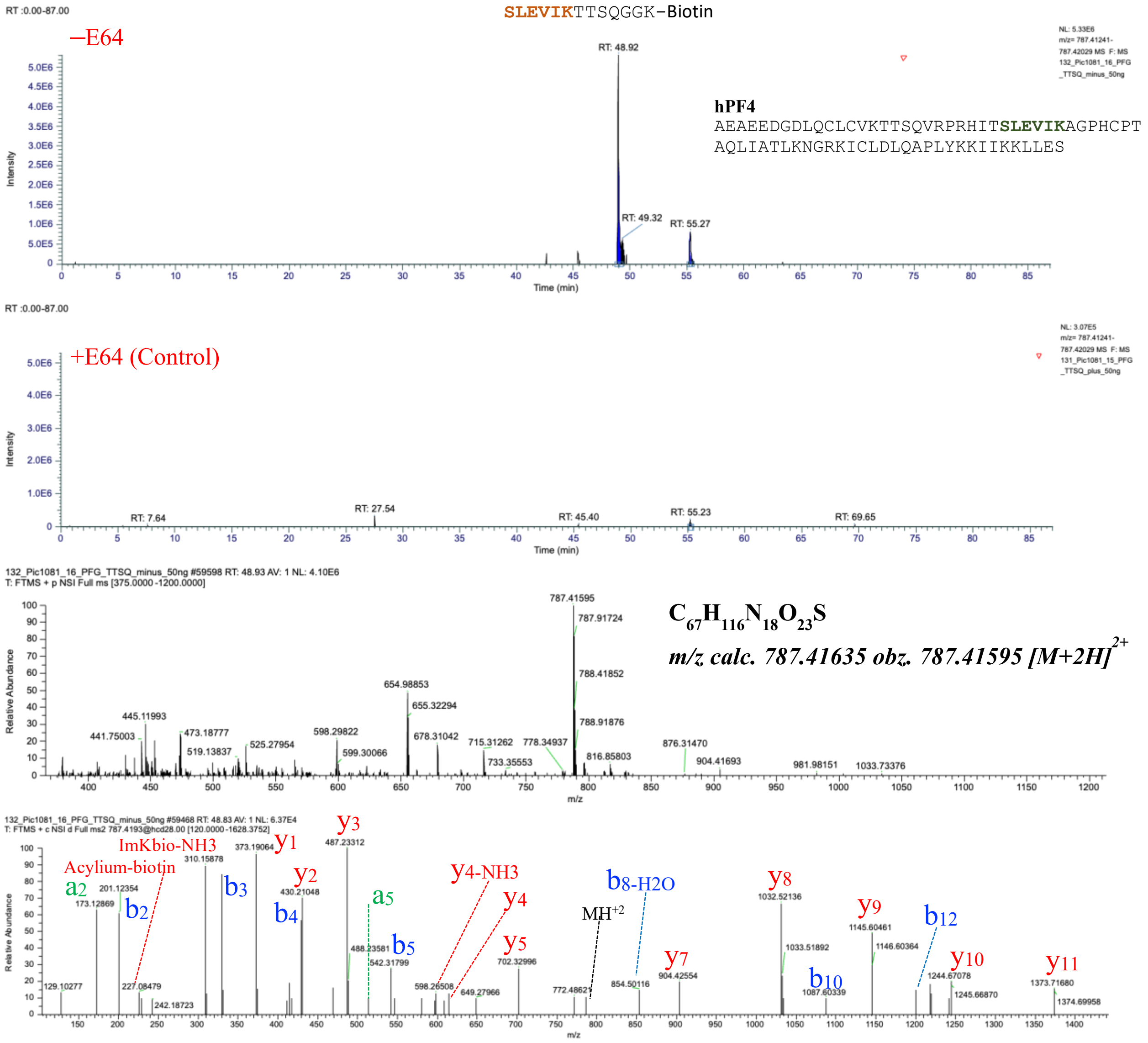

**Fig. S37.** SLEVIKTTSQGGK(Biotin) characterized by LC-MS/MS as a product of hCatS-catalyzed transpeptidation reaction. The figure displays a graphical representation of the extracted ion chromatogram (EIC), mass spectrum, and tandem mass spectrum of SLEVIKTTSQGGK(Biotin), identified from a reaction between TTSQGGK(Biotin) and Human PF4 in the presence of active hCatS. LC-MS/MS analysis was performed on samples treated with active hCatS (–E64) and compared to a control in which the enzyme was inhibited by E64 (+E64). The molecular ion was extracted based on its exact monoisotopic mass, and major fragment ions in the tandem mass spectrum were assigned.

**hPF4** AEAEEDGDLQCLCVKTTSQVRPRHITSLEVIKAGPHCPTAQLIATLKNGRKICLDLQAPLYKKIIKKLLES

### EAEEDGDLQTTSQGGK(Biotin)

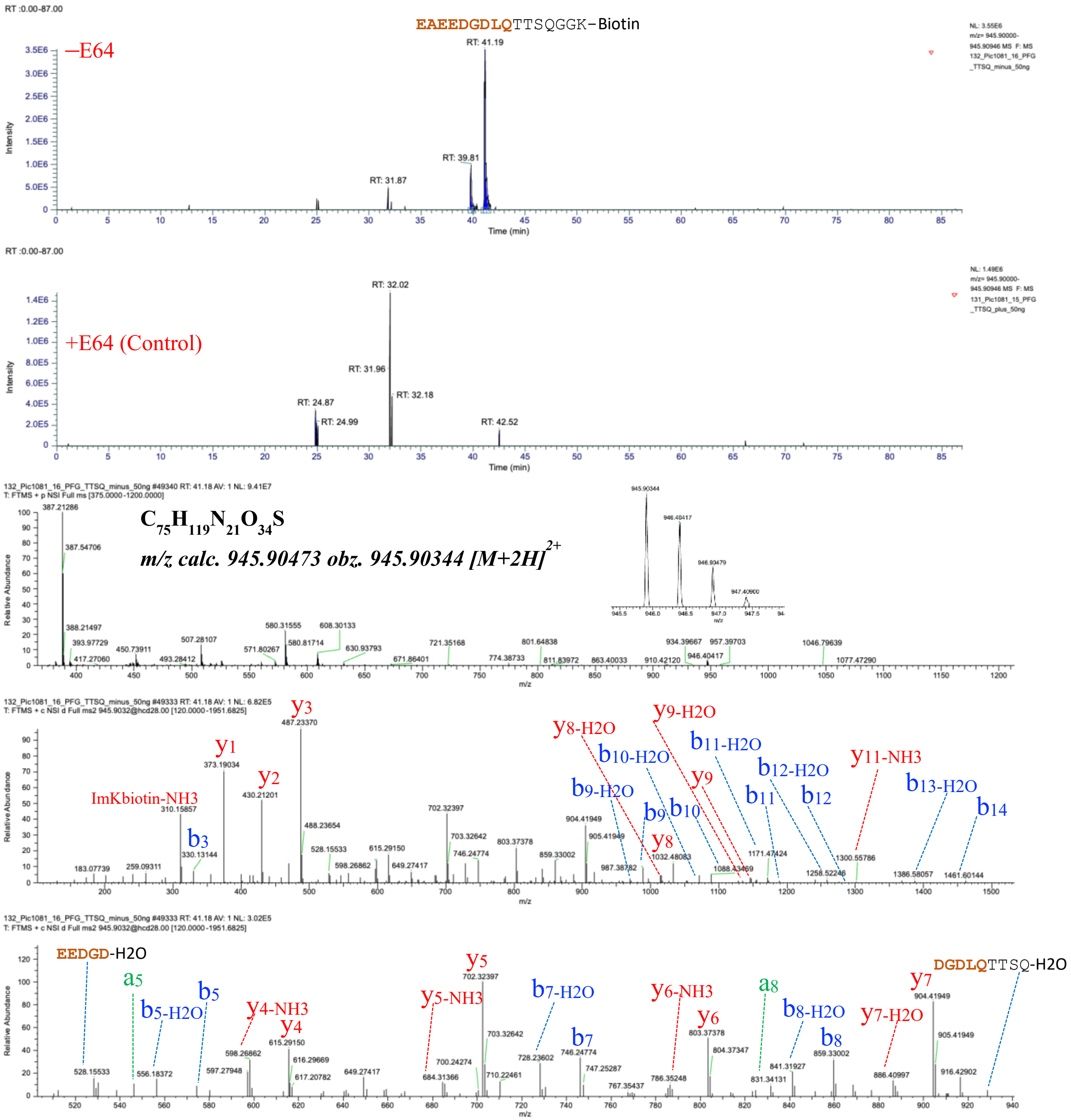

**Fig. S38.** EAEEDGDLQTTSQGGK(Biotin) characterized by LC-MS/MS as a product of hCatS-catalyzed transpeptidation reaction. The figure displays a graphical representation of the extracted ion chromatogram (EIC), mass spectrum, and tandem mass spectrum of EAEEDGDLQTTSQGGK(Biotin), identified from a reaction between TTSQGGK(Biotin) and Human PF4 in the presence of active hCatS. LC-MS/MS analysis was performed on samples treated with active hCatS (–E64) and compared to a control in which the enzyme was inhibited by E64 (+E64). The molecular ion was extracted based on its exact monoisotopic mass, and major fragment ions in the tandem mass spectrum were assigned.

**hPF4** AEAEEDGDLQCLCVKTTSQVRPRHITSLEVIKAGPHCPTAQLIATLKNGRKICLDLQAPLYKKIIKKLLES

### TTSTTTTTTSQGGK(Biotin)

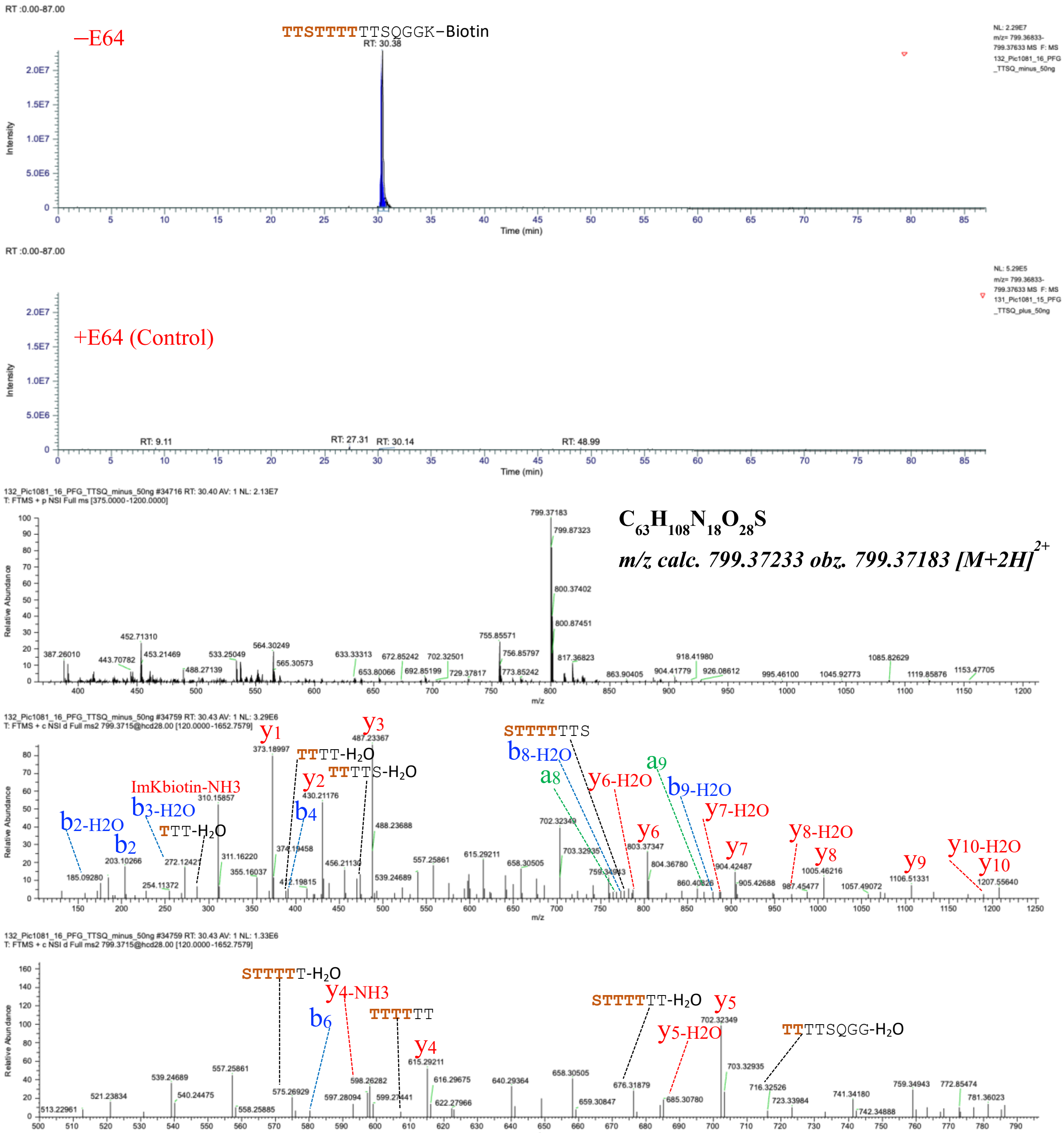

**Fig. S39.** TTSTTTTTTSQGGK(Biotin) characterized by LC-MS/MS as a product of hCatS-catalyzed transpeptidation reaction. The figure displays a graphical representation of the extracted ion chromatogram (EIC), mass spectrum, and tandem mass spectrum of TTSTTTTTTSQGGK(Biotin), identified from a reaction between TTSQGGK(Biotin) and Human PF4 in the presence of active hCatS. LC-MS/MS analysis was performed on samples treated with active hCatS (–E64) and compared to a control in which the enzyme was inhibited by E64 (+E64). The molecular ion was extracted based on its exact monoisotopic mass, and major fragment ions in the tandem mass spectrum were assigned.

VKTTTTSQGGK(Biotin)

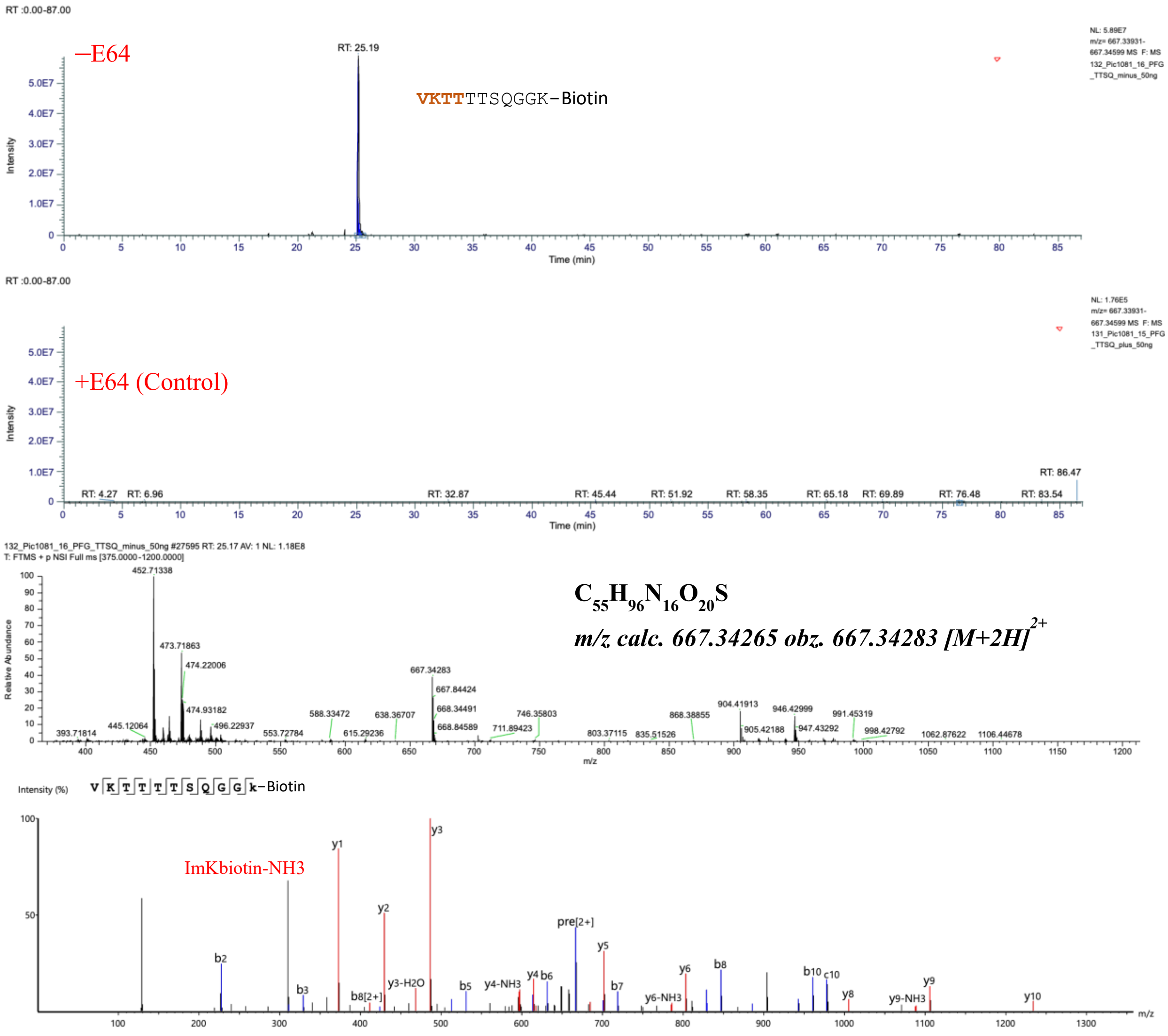

**Fig. S40.** VKTTTTSQGGK(Biotin) characterized by LC-MS/MS as a product of hCatS-catalyzed transpeptidation reaction. The figure displays a graphical representation of the extracted ion chromatogram (EIC), mass spectrum, and tandem mass spectrum of VKTTTTSQGGK(Biotin), identified from a reaction between TTSQGGK(Biotin) and Human PF4 in the presence of active hCatS. LC-MS/MS analysis was performed on samples treated with active hCatS (–E64) and compared to a control in which the enzyme was inhibited by E64 (+E64). The molecular ion was extracted based on its exact monoisotopic mass, and major fragment ions in the tandem mass spectrum were assigned.

VKTTTTTTSQGGK(Biotin)

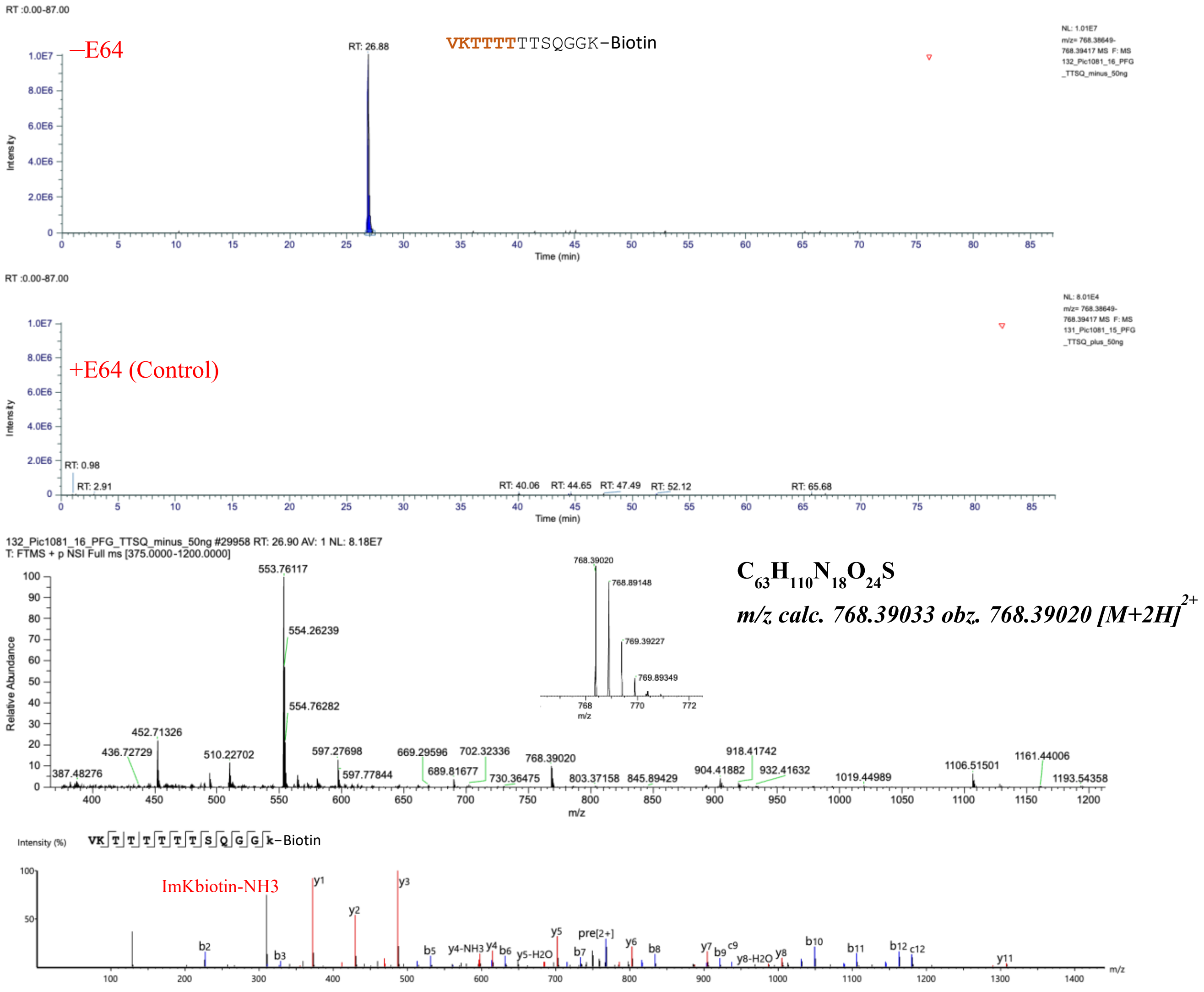

**Fig. S41.** VKTTTTTTSQGGK(Biotin) characterized by LC-MS/MS as a product of hCatS-catalyzed transpeptidation reaction. The figure displays a graphical representation of the extracted ion chromatogram (EIC), mass spectrum, and tandem mass spectrum of VKTTTTTTSQGGK(Biotin), identified from a reaction between TTSQGGK(Biotin) and Human PF4 in the presence of active hCatS. LC-MS/MS analysis was performed on samples treated with active hCatS (–E64) and compared to a control in which the enzyme was inhibited by E64 (+E64). The molecular ion was extracted based on its exact monoisotopic mass, and major fragment ions in the tandem mass spectrum were assigned.

TTTTTTSQGGK(Biotin)

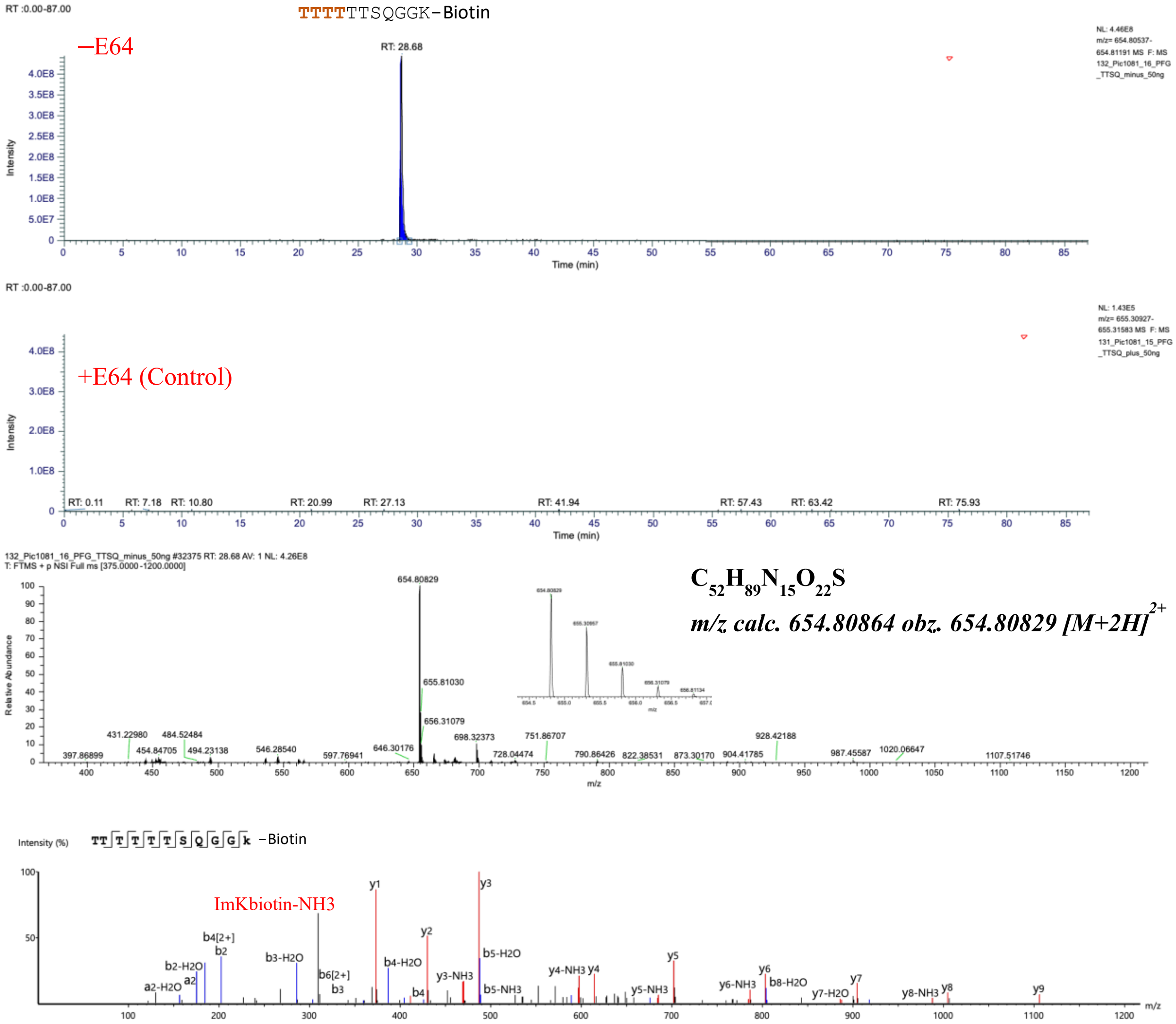

**Fig. S42.** TTTTTTSQGGK(Biotin) characterized by LC-MS/MS as a product of hCatS-catalyzed transpeptidation reaction. The figure displays a graphical representation of the extracted ion chromatogram (EIC), mass spectrum, and tandem mass spectrum of TTTTTTSQGGK(Biotin), identified from a reaction between TTSQGGK(Biotin) and Human PF4 in the presence of active hCatS. LC-MS/MS analysis was performed on samples treated with active hCatS (–E64) and compared to a control in which the enzyme was inhibited by E64 (+E64). The molecular ion was extracted based on its exact monoisotopic mass, and major fragment ions in the tandem mass spectrum were assigned.

#### VKTTSTTSQGGK(Biotin)

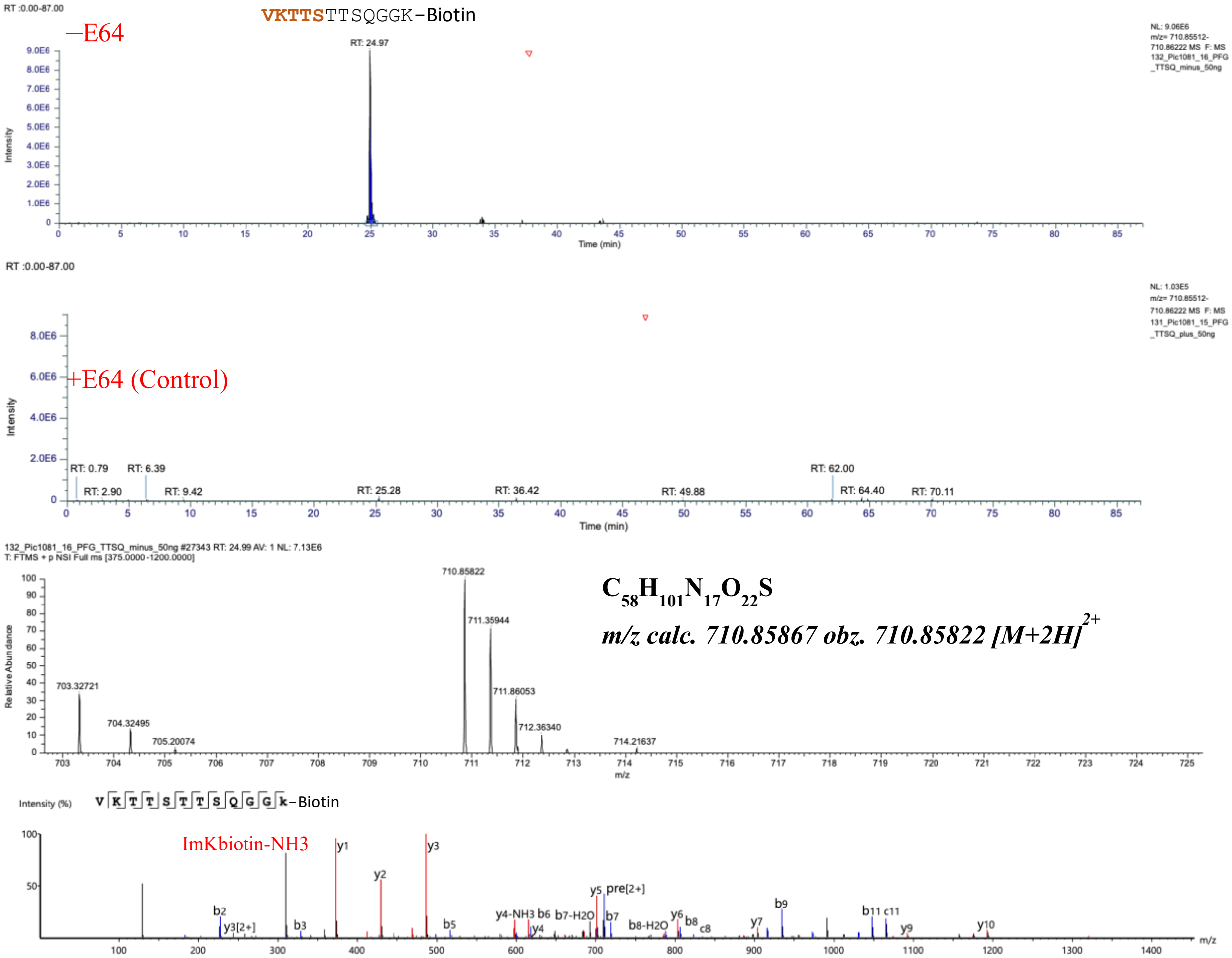

**Fig. S43.** VKTTSTTSQGGK(Biotin) characterized by LC-MS/MS as a product of hCatS-catalyzed transpeptidation reaction. The figure displays a graphical representation of the extracted ion chromatogram (EIC), mass spectrum, and tandem mass spectrum of VKTTSTTSQGGK(Biotin), identified from a reaction between TTSQGGK(Biotin) and Human PF4 in the presence of active hCatS. LC-MS/MS analysis was performed on samples treated with active hCatS (–E64) and compared to a control in which the enzyme was inhibited by E64 (+E64). The molecular ion was extracted based on its exact monoisotopic mass, and major fragment ions in the tandem mass spectrum were assigned.

APLYKKIIKTTSQGGK(Biotin)

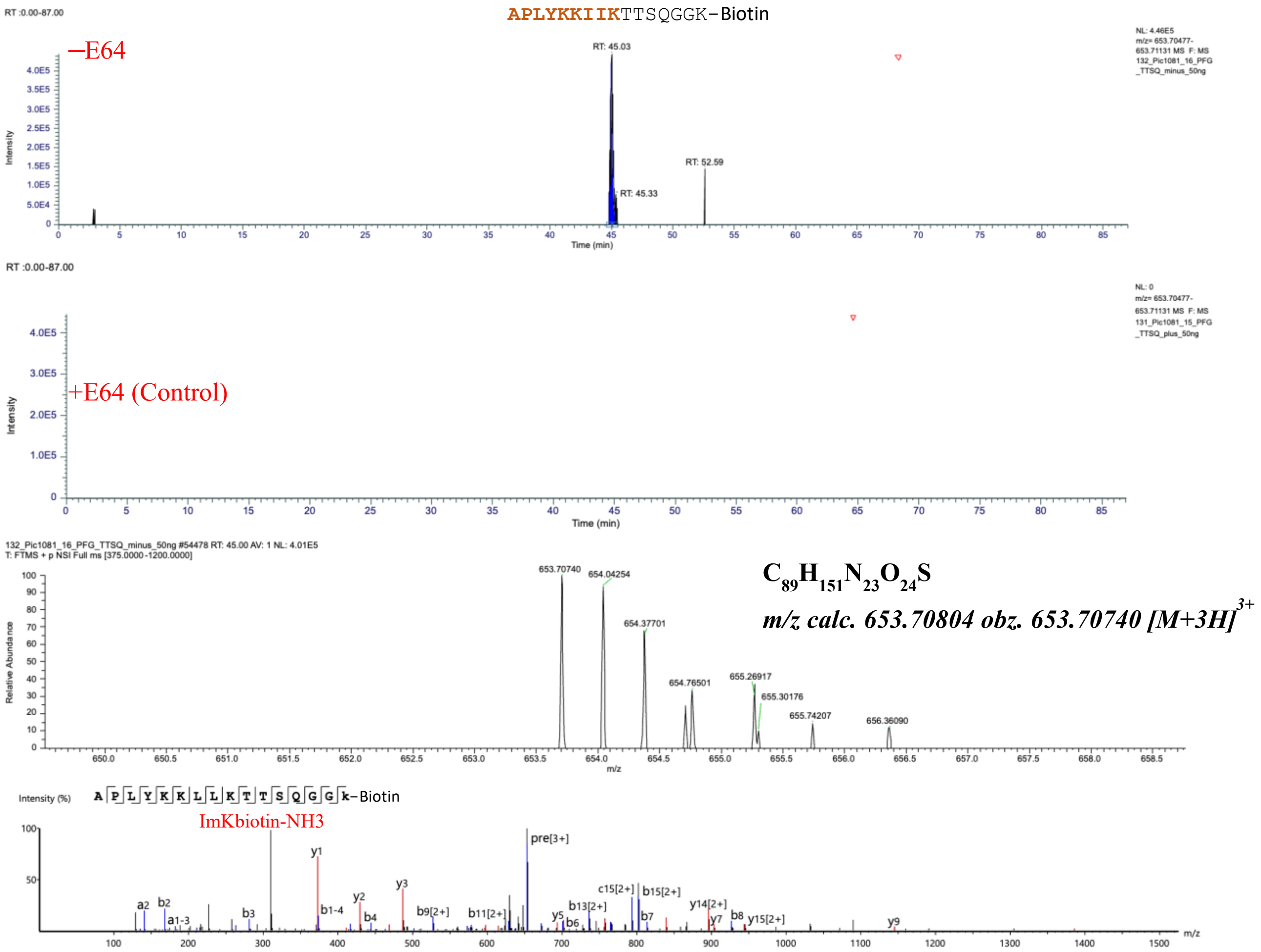

**Fig. S44.** APLYKKIIKTTSQGGK(Biotin) characterized by LC-MS/MS as a product of hCatS-catalyzed transpeptidation reaction. The figure displays a graphical representation of the extracted ion chromatogram (EIC), mass spectrum, and tandem mass spectrum of APLYKKIIKTTSQGGK(Biotin), identified from a reaction between TTSQGGK(Biotin) and Human PF4 in the presence of active hCatS. LC-MS/MS analysis was performed on samples treated with active hCatS (–E64) and compared to a control in which the enzyme was inhibited by E64 (+E64). The molecular ion was extracted based on its exact monoisotopic mass, and major fragment ions in the tandem mass spectrum were assigned.

APLYKTTSQGGK(Biotin)

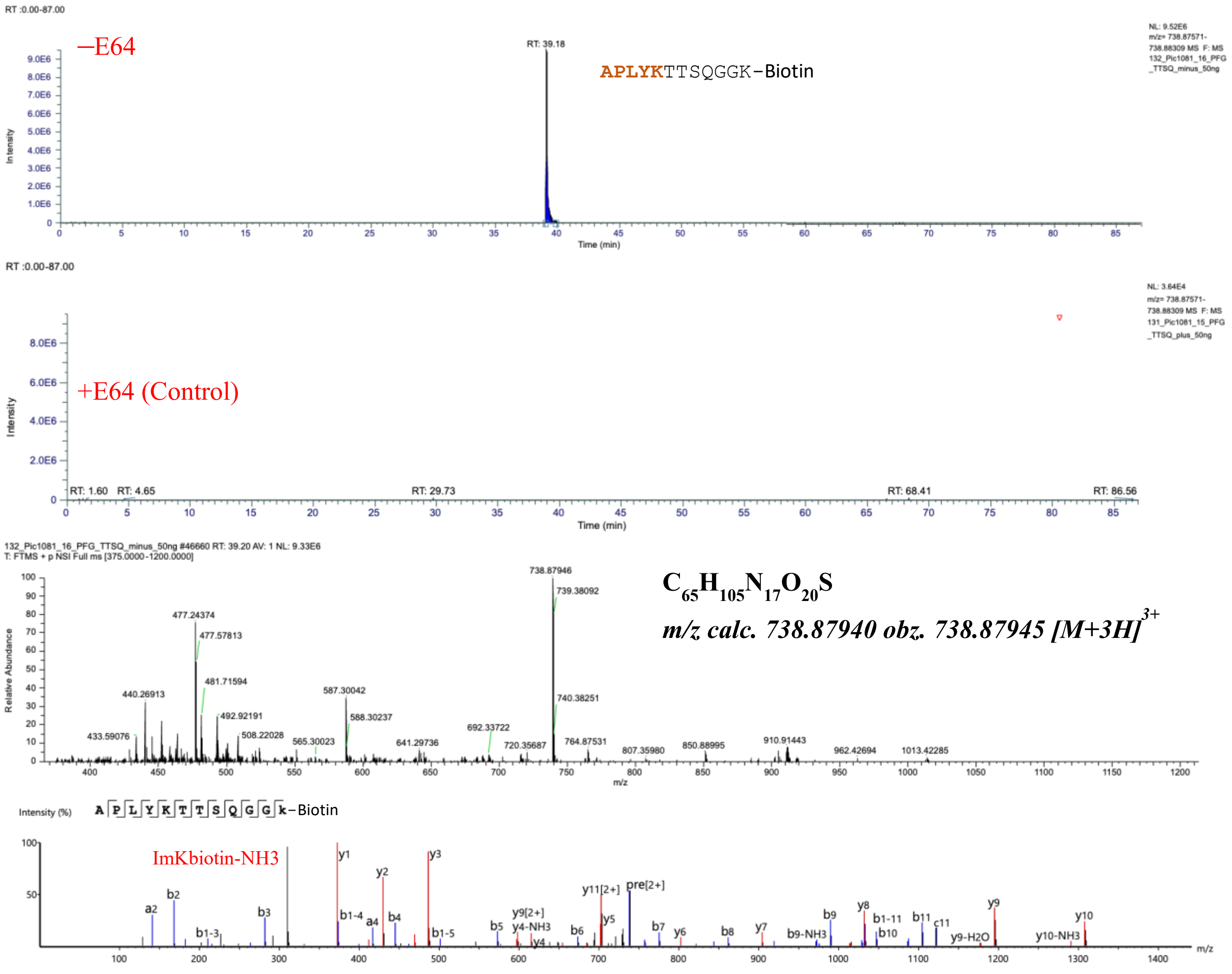

**Fig. S45.** APLYKTTSQGGK(Biotin) characterized by LC-MS/MS as a product of hCatS-catalyzed transpeptidation reaction. The figure displays a graphical representation of the extracted ion chromatogram (EIC), mass spectrum, and tandem mass spectrum of APLYKTTSQGGK(Biotin), identified from a reaction between TTSQGGK(Biotin) and Human PF4 in the presence of active hCatS. LC-MS/MS analysis was performed on samples treated with active hCatS (–E64) and compared to a control in which the enzyme was inhibited by E64 (+E64). The molecular ion was extracted based on its exact monoisotopic mass, and major fragment ions in the tandem mass spectrum were assigned.

LEVIKTTSQGGK(Biotin)

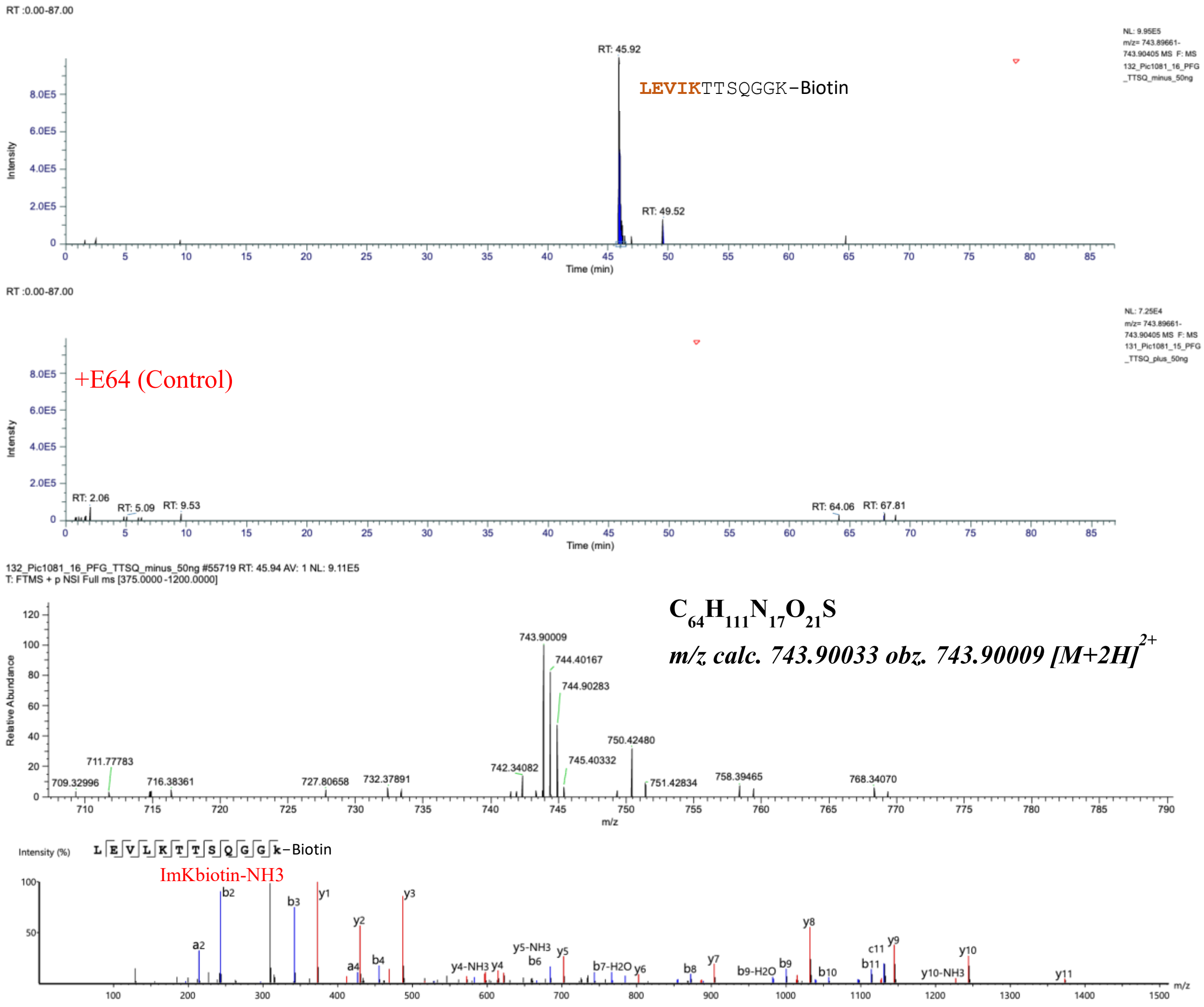

**Fig. S46.** LEVIKTTSQGGK(Biotin) characterized by LC-MS/MS as a product of hCatS-catalyzed transpeptidation reaction. The figure displays a graphical representation of the extracted ion chromatogram (EIC), mass spectrum, and tandem mass spectrum of LEVIKTTSQGGK(Biotin), identified from a reaction between TTSQGGK(Biotin) and Human PF4 in the presence of active hCatS. LC-MS/MS analysis was performed on samples treated with active hCatS (–E64) and compared to a control in which the enzyme was inhibited by E64 (+E64). The molecular ion was extracted based on its exact monoisotopic mass, and major fragment ions in the tandem mass spectrum were assigned.

LEVIKTTSQGGK(Biotin)

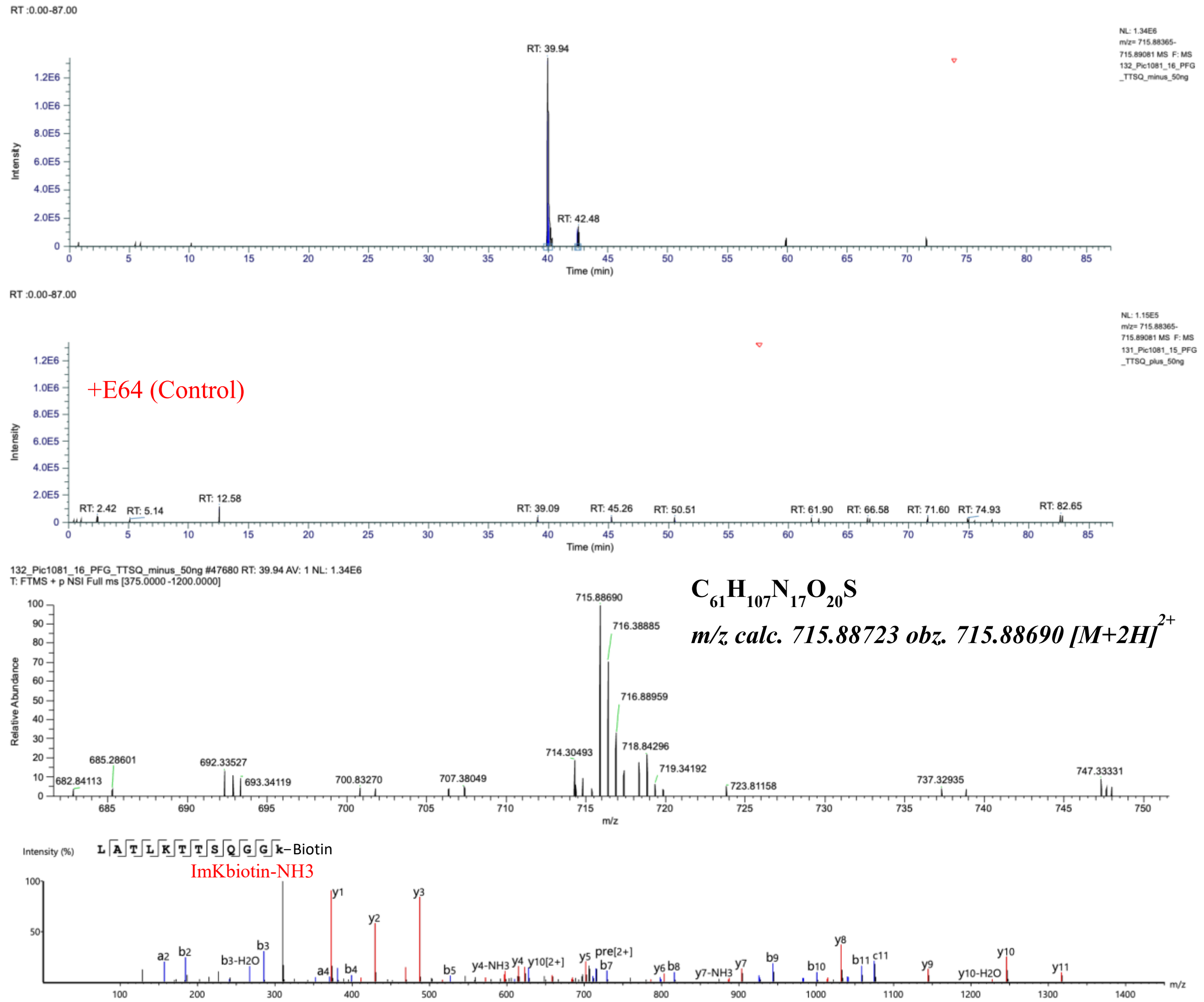

**Fig. S47.** LEVIKTTSQGGK(Biotin) characterized by LC-MS/MS as a product of hCatS-catalyzed transpeptidation reaction. The figure displays a graphical representation of the extracted ion chromatogram (EIC), mass spectrum, and tandem mass spectrum of LEVIKTTSQGGK(Biotin), identified from a reaction between TTSQGGK(Biotin) and Human PF4 in the presence of active hCatS. LC-MS/MS analysis was performed on samples treated with active hCatS (–E64) and compared to a control in which the enzyme was inhibited by E64 (+E64). The molecular ion was extracted based on its exact monoisotopic mass, and major fragment ions in the tandem mass spectrum were assigned.

LEVIKTTSQGGK(Biotin)

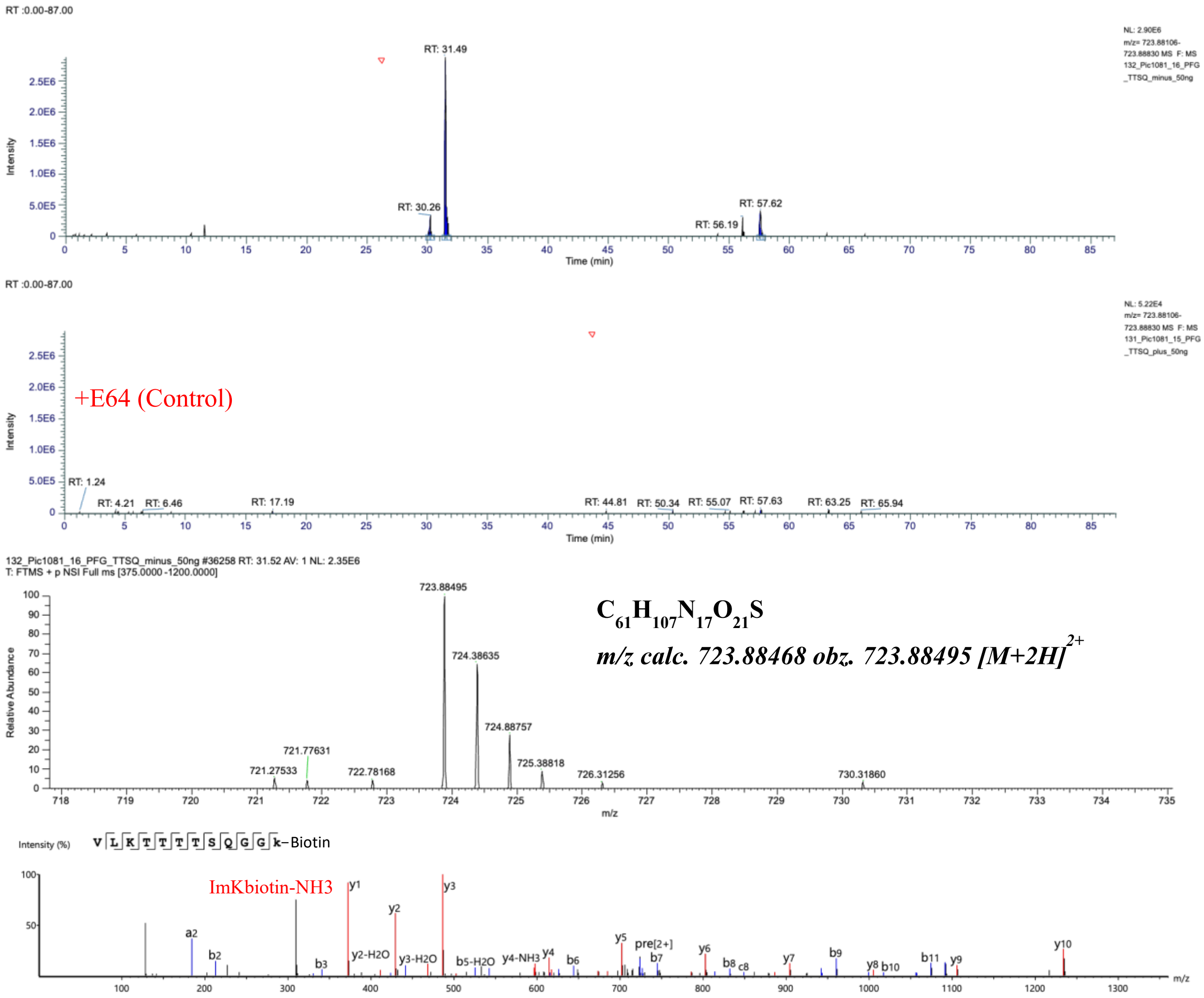

**Fig. S48.** LEVIKTTSQGGK(Biotin) characterized by LC-MS/MS as a product of hCatS-catalyzed transpeptidation reaction. The figure displays a graphical representation of the extracted ion chromatogram (EIC), mass spectrum, and tandem mass spectrum of LEVIKTTSQGGK(Biotin), identified from a reaction between TTSQGGK(Biotin) and Human PF4 in the presence of active hCatS. LC-MS/MS analysis was performed on samples treated with active hCatS (–E64) and compared to a control in which the enzyme was inhibited by E64 (+E64). The molecular ion was extracted based on its exact monoisotopic mass, and major fragment ions in the tandem mass spectrum were assigned.

LIATTTSQGGK(Biotin)

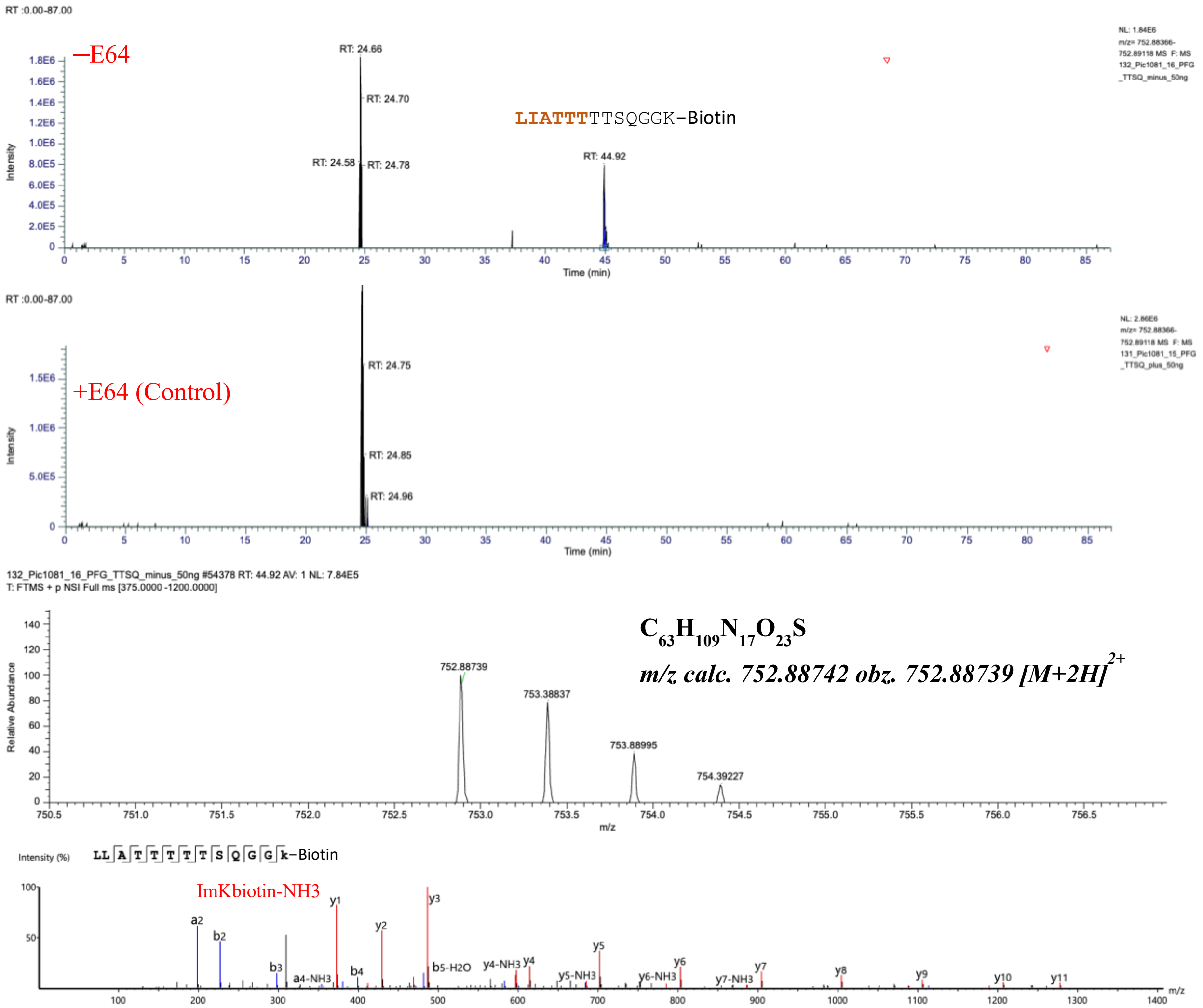

**Fig. S49.** LIATTTSQGGK(Biotin) characterized by LC-MS/MS as a product of hCatS-catalyzed transpeptidation reaction. The figure displays a graphical representation of the extracted ion chromatogram (EIC), mass spectrum, and tandem mass spectrum of LEVIKTTSQGGK(Biotin), identified from a reaction between TTSQGGK(Biotin) and Human PF4 in the presence of active hCatS. LC-MS/MS analysis was performed on samples treated with active hCatS (–E64) and compared to a control in which the enzyme was inhibited by E64 (+E64). The molecular ion was extracted based on its exact monoisotopic mass, and major fragment ions in the tandem mass spectrum were assigned.

KKIIKTTSQGGK(Biotin)

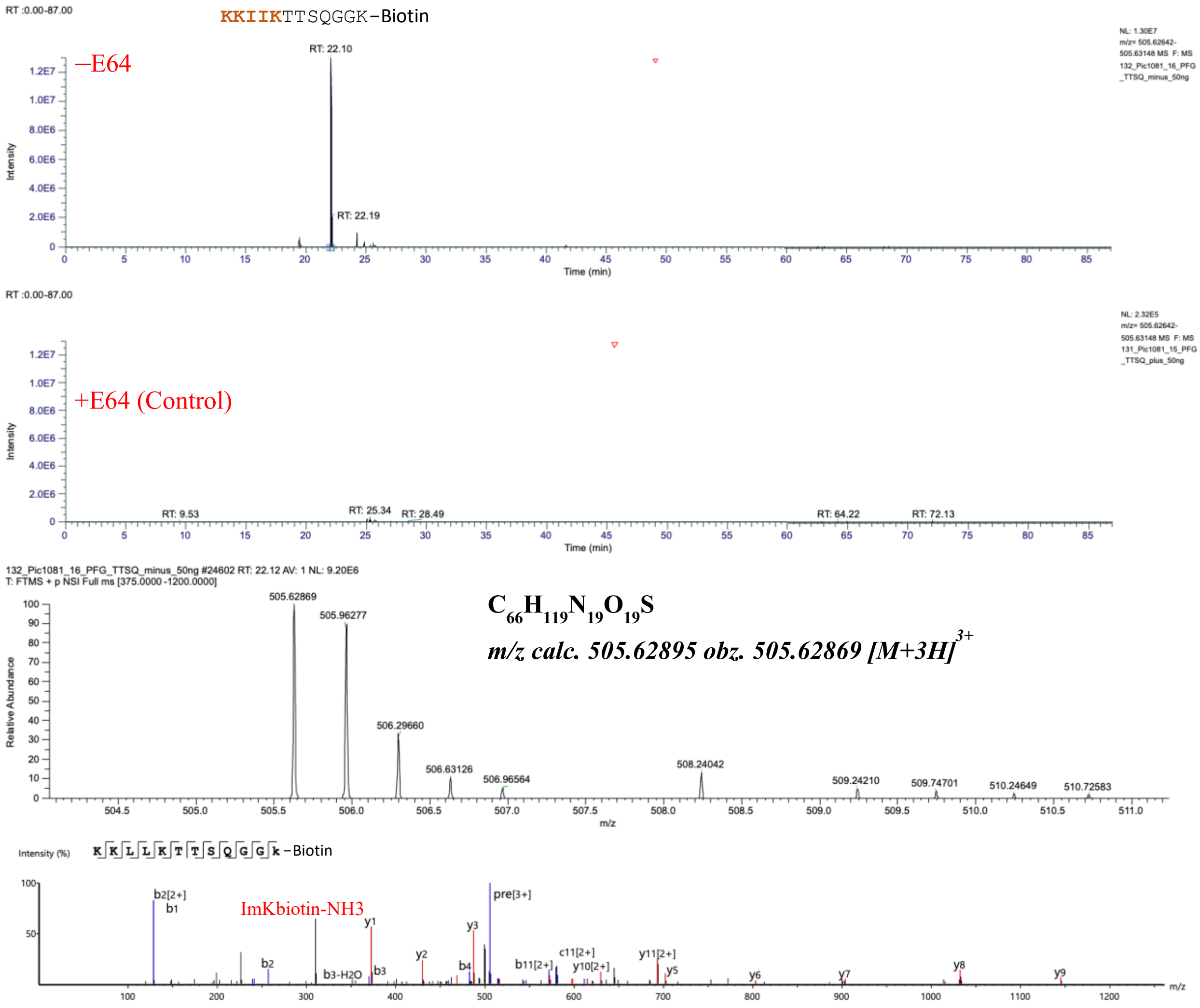

**Fig. S50.** KKIIKTTSQGGK(Biotin) characterized by LC-MS/MS as a product of hCatS-catalyzed transpeptidation reaction. The figure displays a graphical representation of the extracted ion chromatogram (EIC), mass spectrum, and tandem mass spectrum of KKIIKTTSQGGK(Biotin), identified from a reaction between TTSQGGK(Biotin) and Human PF4 in the presence of active hCatS. LC-MS/MS analysis was performed on samples treated with active hCatS (–E64) and compared to a control in which the enzyme was inhibited by E64 (+E64). The molecular ion was extracted based on its exact monoisotopic mass, and major fragment ions in the tandem mass spectrum were assigned.

VKTTTTTTSQGGK(Biotin)

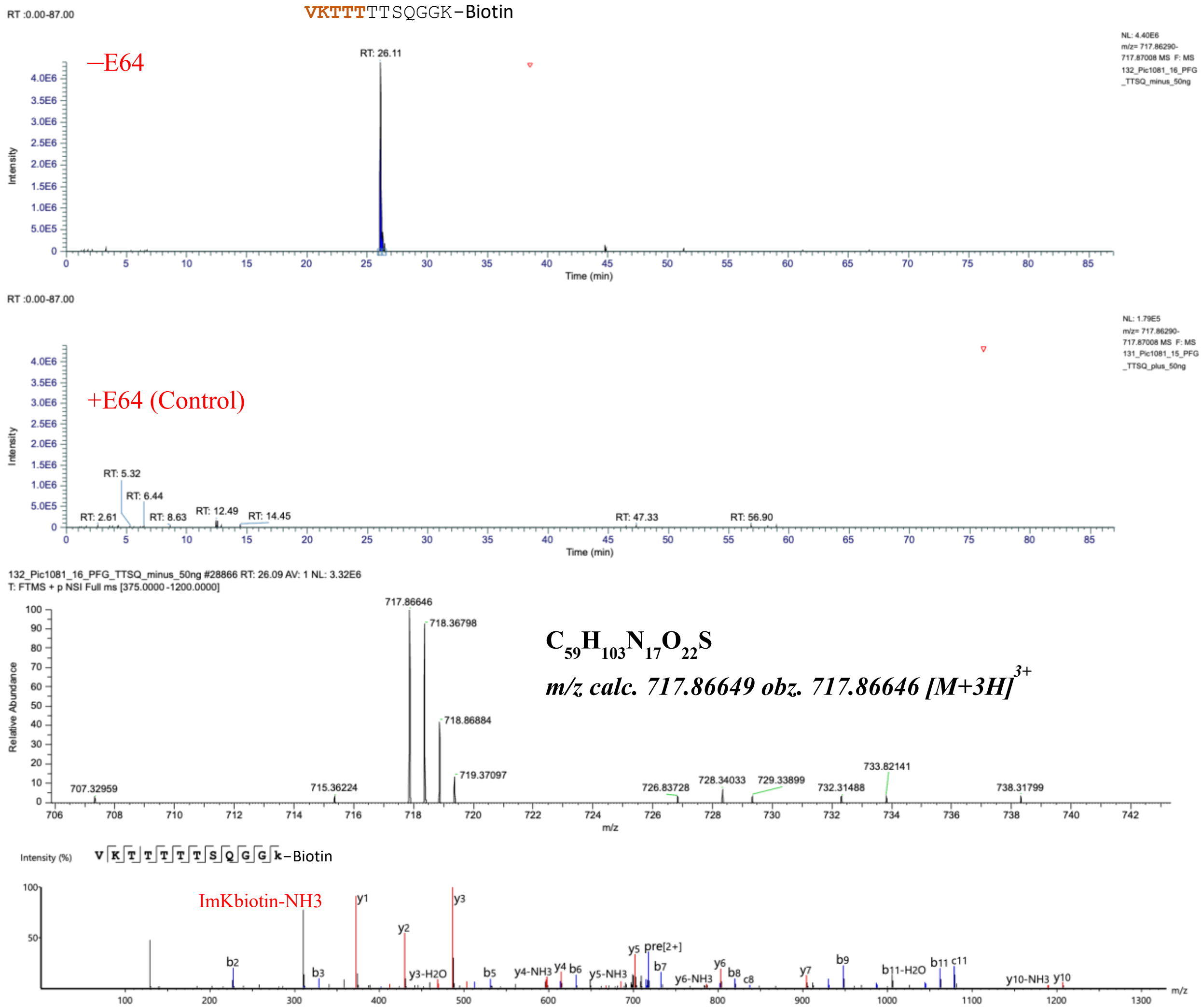

**Fig. S51.** VKTTTTTTSQGGK(Biotin) characterized by LC-MS/MS as a product of hCatS-catalyzed transpeptidation reaction. The figure displays a graphical representation of the extracted ion chromatogram (EIC), mass spectrum, and tandem mass spectrum of VKTTTTTTSQGGK(Biotin), identified from a reaction between TTSQGGK(Biotin) and Human PF4 in the presence of active hCatS. LC-MS/MS analysis was performed on samples treated with active hCatS (–E64) and compared to a control in which the enzyme was inhibited by E64 (+E64). The molecular ion was extracted based on its exact monoisotopic mass, and major fragment ions in the tandem mass spectrum were assigned.

TTSTTTTSQGGK(Biotin)

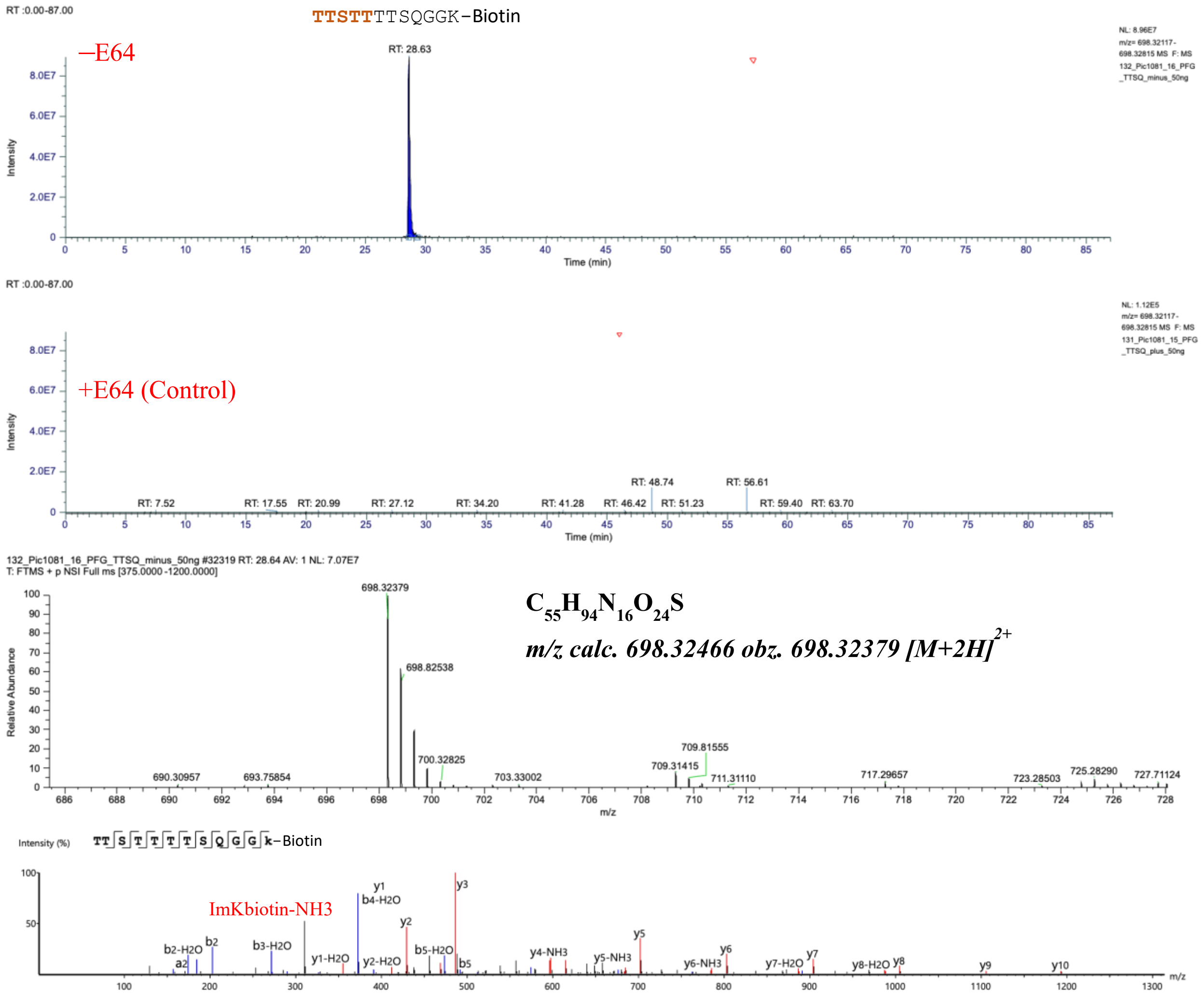

**Fig. S52.** TTSTTTTSQGGK(Biotin) characterized by LC-MS/MS as a product of hCatS-catalyzed transpeptidation reaction. The figure displays a graphical representation of the extracted ion chromatogram (EIC), mass spectrum, and tandem mass spectrum of TTSTTTTSQGGK(Biotin), identified from a reaction between TTSQGGK(Biotin) and Human PF4 in the presence of active hCatS. LC-MS/MS analysis was performed on samples treated with active hCatS (–E64) and compared to a control in which the enzyme was inhibited by E64 (+E64). The molecular ion was extracted based on its exact monoisotopic mass, and major fragment ions in the tandem mass spectrum were assigned.

TTSTTTSQGGK(Biotin)

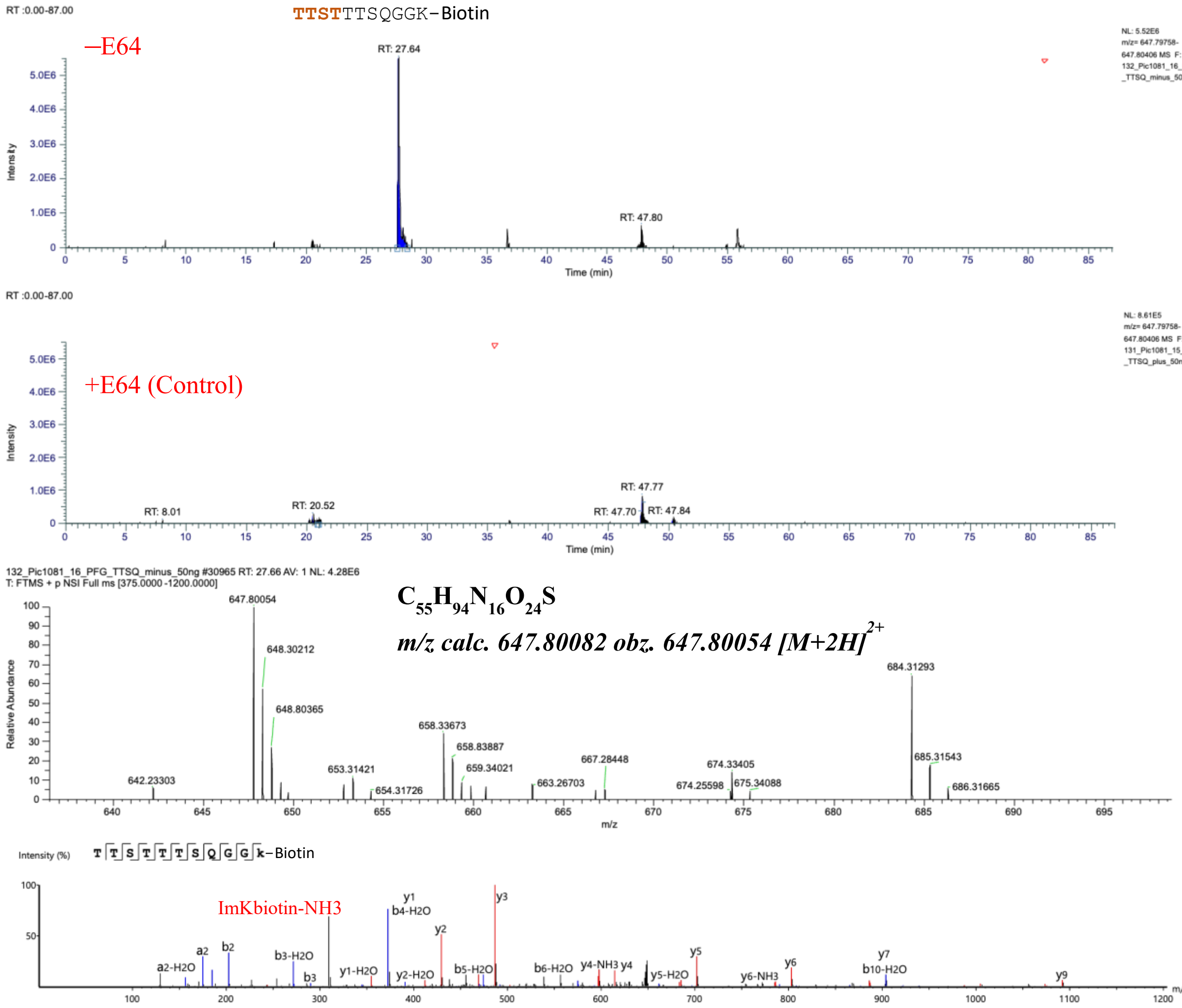

**Fig. S53.** TTSTTTSQGGK(Biotin) characterized by LC-MS/MS as a product of hCatS-catalyzed transpeptidation reaction. The figure displays a graphical representation of the extracted ion chromatogram (EIC), mass spectrum, and tandem mass spectrum of TTSTTTSQGGK(Biotin), identified from a reaction between TTSQGGK(Biotin) and Human PF4 in the presence of active hCatS. LC-MS/MS analysis was performed on samples treated with active hCatS (–E64) and compared to a control in which the enzyme was inhibited by E64 (+E64). The molecular ion was extracted based on its exact monoisotopic mass, and major fragment ions in the tandem mass spectrum were assigned.

TTSQTTSQGGK(Biotin)

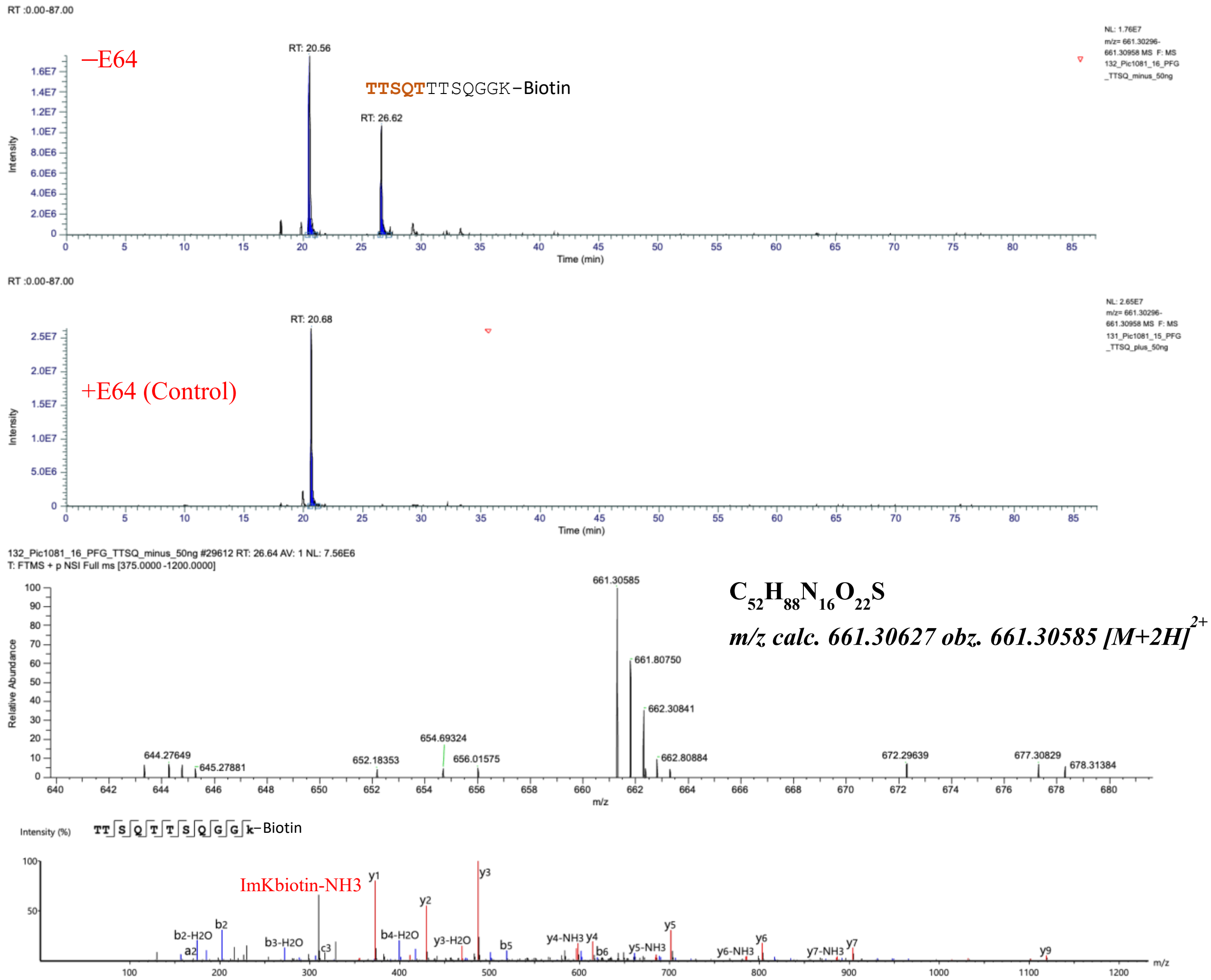

**Fig. S54.** TTSQTTSQGGK(Biotin) characterized by LC-MS/MS as a product of hCatS-catalyzed transpeptidation reaction. The figure displays a graphical representation of the extracted ion chromatogram (EIC), mass spectrum, and tandem mass spectrum of TTSQTTSQGGK(Biotin), identified from a reaction between TTSQGGK(Biotin) and Human PF4 in the presence of active hCatS. LC-MS/MS analysis was performed on samples treated with active hCatS (–E64) and compared to a control in which the enzyme was inhibited by E64 (+E64). The molecular ion was extracted based on its exact monoisotopic mass, and major fragment ions in the tandem mass spectrum were assigned.

SDS-PAGE analysis of S1 and S2 Subunits and Avidin-FITC blotting of Full length Spike of SARS-CoV-2

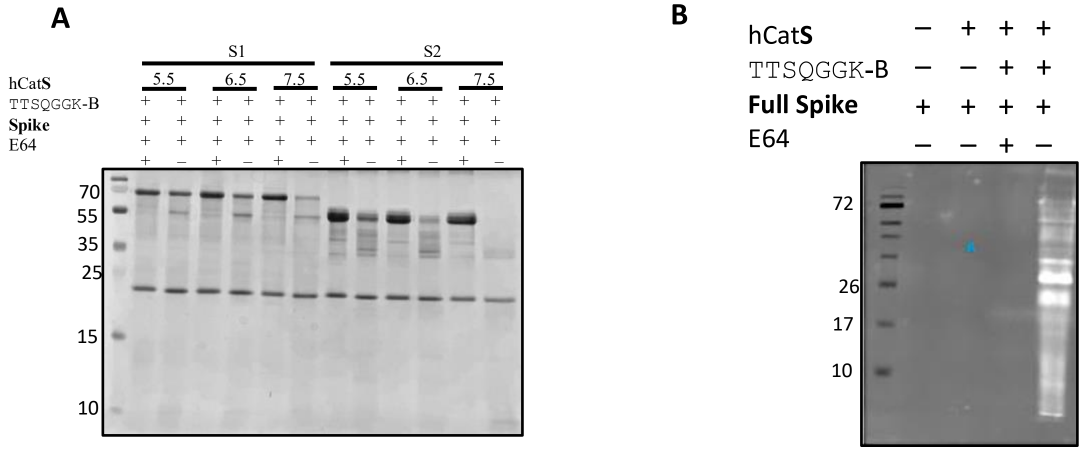

**Fig. S55: A:** SDS-PAGE (15%) analysis of the SARS-CoV-2 Spike protein subunits S1 (75 kDa) and S2 (56 kDa) (non-glycosylated, overexpressed in *E. coli*) incubated with (hCatS) under two conditions: inactive hCatS (+E64) and active hCatS (-E64), in the presence of TTSQGGK-Biotin as the substrate. A fluorescent image corresponding to this experiment is shown in Figure 4G (main text). **B:** Avidin-FITC blotting of Full length Spike of SARS-CoV-2 (mammalian cell expressed) subunits.

**Characterization of Spike derived biotinylated (TTSQGGK-Biotin conjugates) transpeptides**

ALEPLVDLPIGTTSQGGK(Biotin)

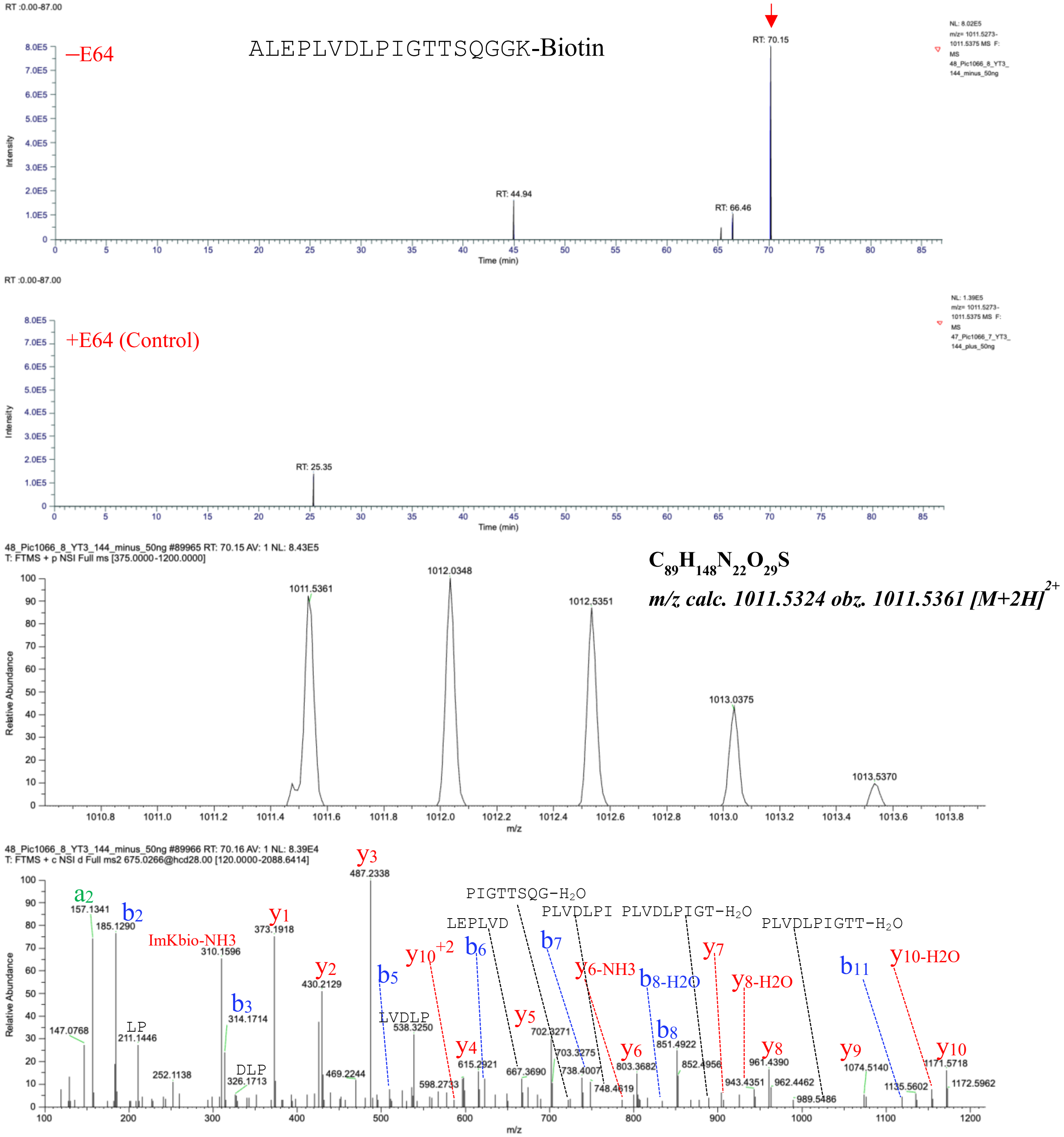

**Fig. S56.** ALEPLVDLPIGTTSQGGK(Biotin) characterized by LC-MS/MS as a product of hCatS-catalyzed transpeptidation reaction. The figure displays a graphical representation of the extracted ion chromatogram (EIC), mass spectrum, and tandem mass spectrum of ALEPLVDLPIGTTSQGGK(Biotin), identified from a reaction between TTSQGGK(Biotin) and Spike of SARS-CoV-2 in the presence of active hCatS.

NSYECDIPIGTTSQGGK(Biotin)

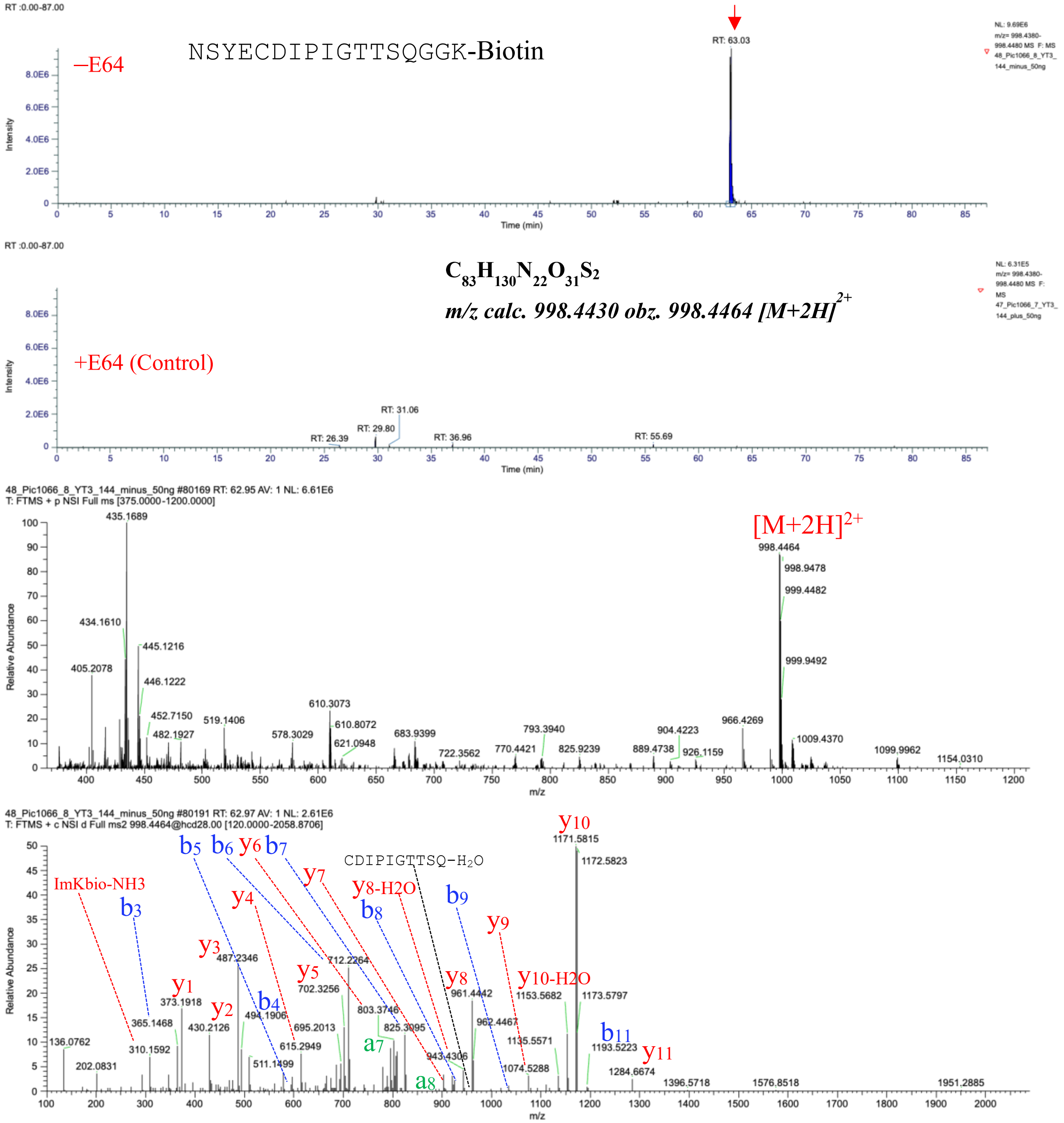

**Fig. S57.** NSYECDIPIGTTSQGGK(Biotin) characterized by LC-MS/MS as a product of hCatS-catalyzed transpeptidation reaction. The figure displays a graphical representation of the extracted ion chromatogram (EIC), mass spectrum, and tandem mass spectrum of NSYECDIPIGTTSQGGK(Biotin), identified from a reaction between TTSQGGK(Biotin) and Spike of SARS-CoV-2 in the presence of active hCatS.

FQPTNGVGTTSQGGK(Biotin)

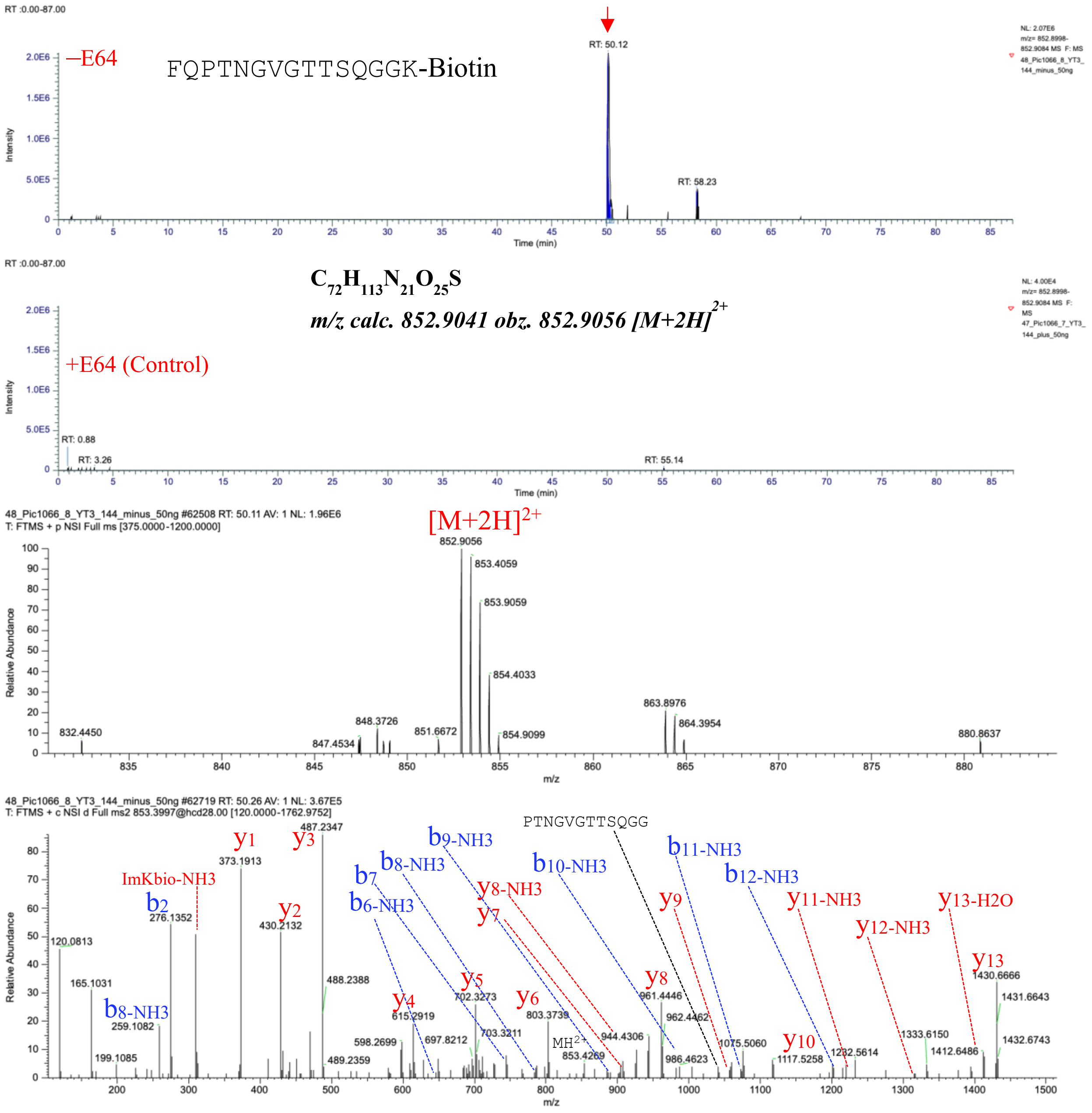

**Fig. S58.** FQPTNGVGTTSQGGK(Biotin) characterized by LC-MS/MS as a product of hCatS-catalyzed transpeptidation reaction. The figure displays a graphical representation of the extracted ion chromatogram (EIC), mass spectrum, and tandem mass spectrum of FQPTNGVGTTSQGGK(Biotin), identified from a reaction between TTSQGGK(Biotin) and Spike of SARS-CoV-2 in the presence of active hCatS. LC-MS/MS analysis was performed on samples treated with active hCatS (–E64) and compared to a control in which the enzyme was inhibited by E64 (+E64). The molecular ion was extracted based on its exact monoisotopic mass, and major fragment ions in the tandem mass spectrum were assigned.

LDPLGTTSQGGK(Biotin)

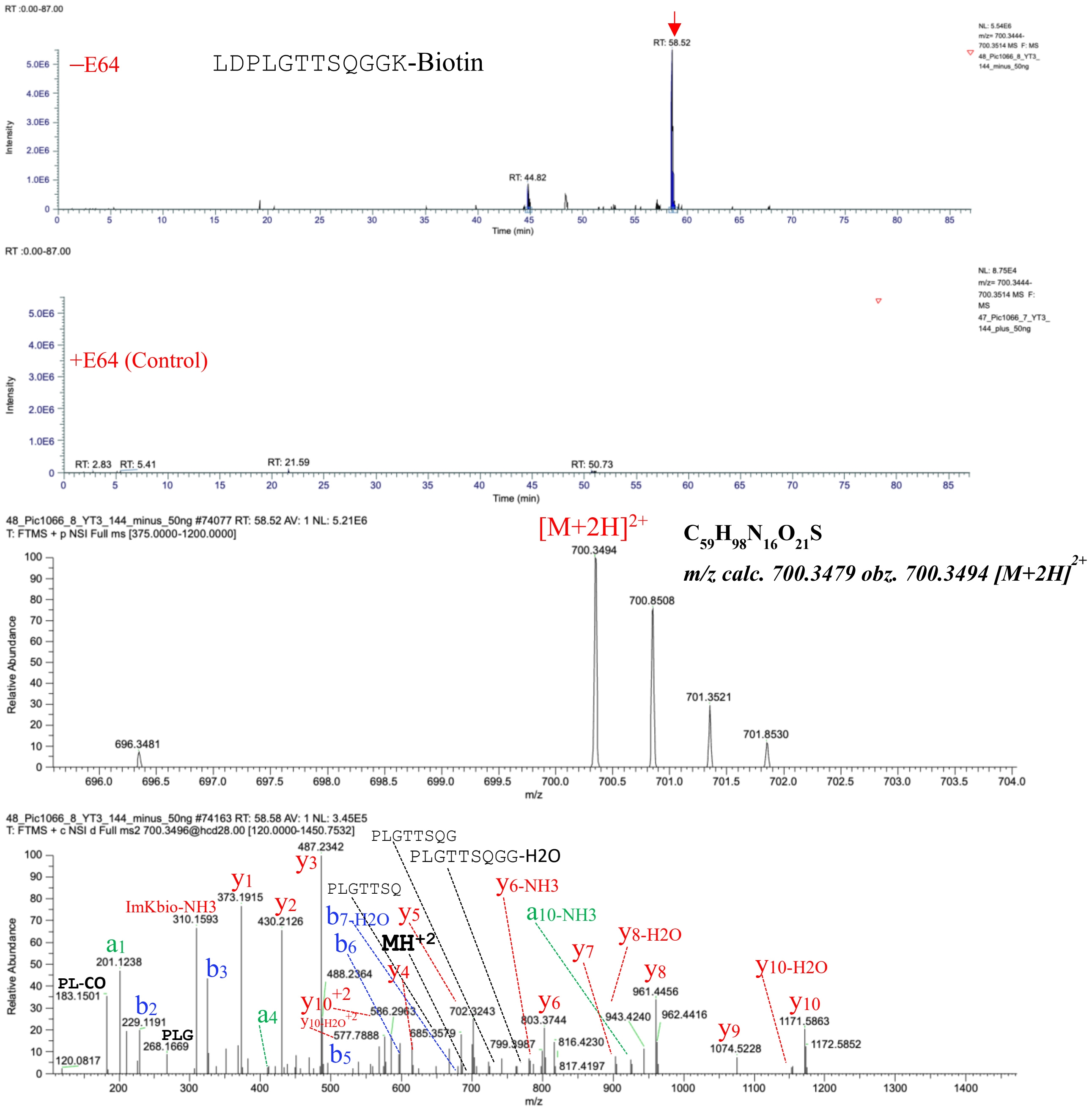

**Fig. S59.** LDPLGTTSQGGK(Biotin) characterized by LC-MS/MS as a product of hCatS-catalyzed transpeptidation reaction. The figure displays a graphical representation of the extracted ion chromatogram (EIC), mass spectrum, and tandem mass spectrum of LDPLGTTSQGGK(Biotin), identified from a reaction between TTSQGGK(Biotin) and Spike of SARS-CoV-2 in the presence of active hCatS.

DEMIAQYTTTSQGGK(Biotin)

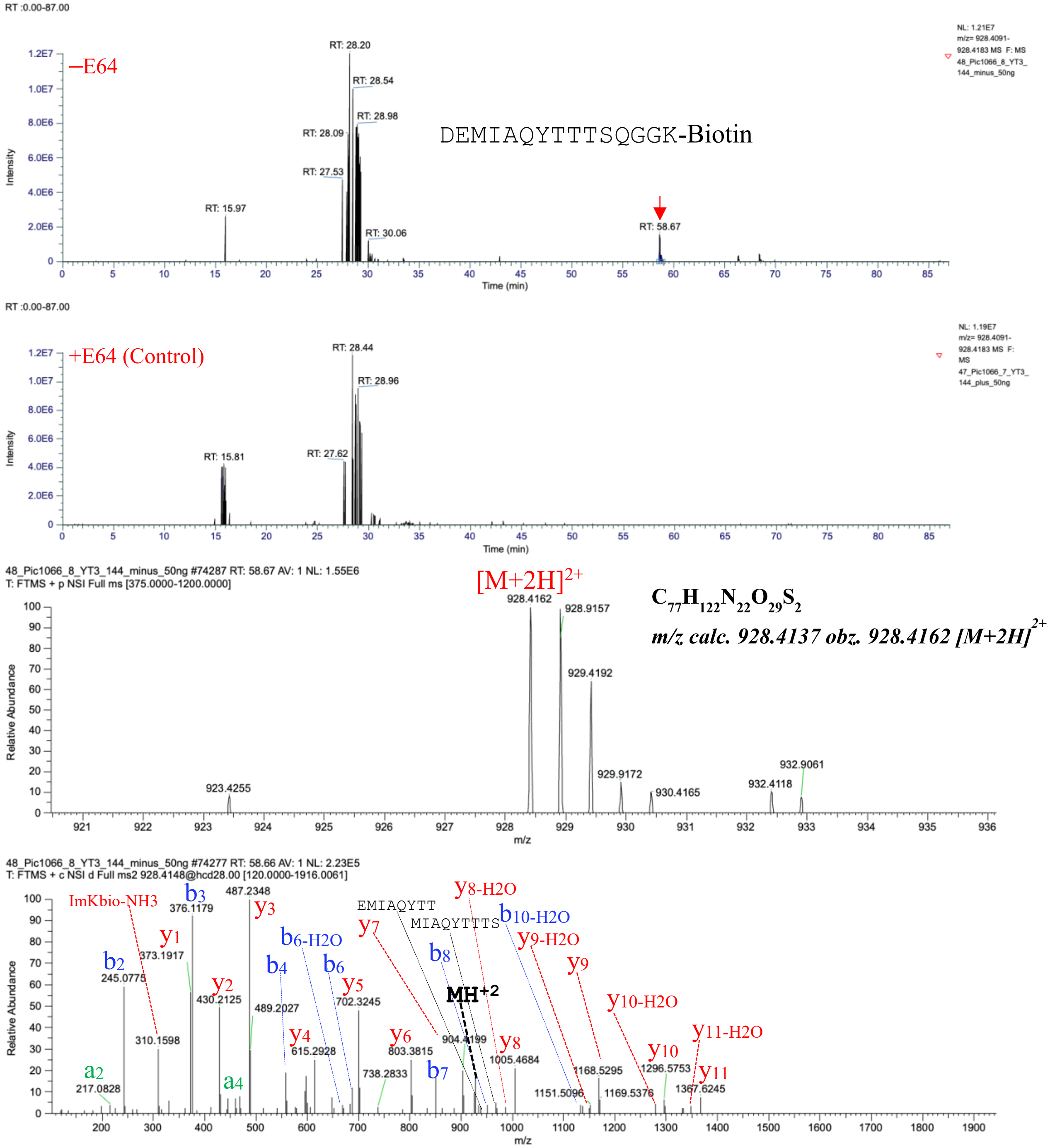

**Fig. S60.** DEMIAQYTTTSQGGK(Biotin) characterized by LC-MS/MS as a product of hCatS-catalyzed transpeptidation reaction. The figure displays a graphical representation of the extracted ion chromatogram (EIC), mass spectrum, and tandem mass spectrum of DEMIAQYTTTSQGGK(Biotin), identified from a reaction between TTSQGGK(Biotin) and Spike of SARS-CoV-2 in the presence of active hCatS. LC-MS/MS analysis was performed on samples treated with active hCatS (–E64) and compared to a control in which the enzyme was inhibited by E64 (+E64). The molecular ion was extracted based on its exact monoisotopic mass, and major fragment ions in the tandem mass spectrum were assigned.

HAPATTTSQGGK(Biotin)

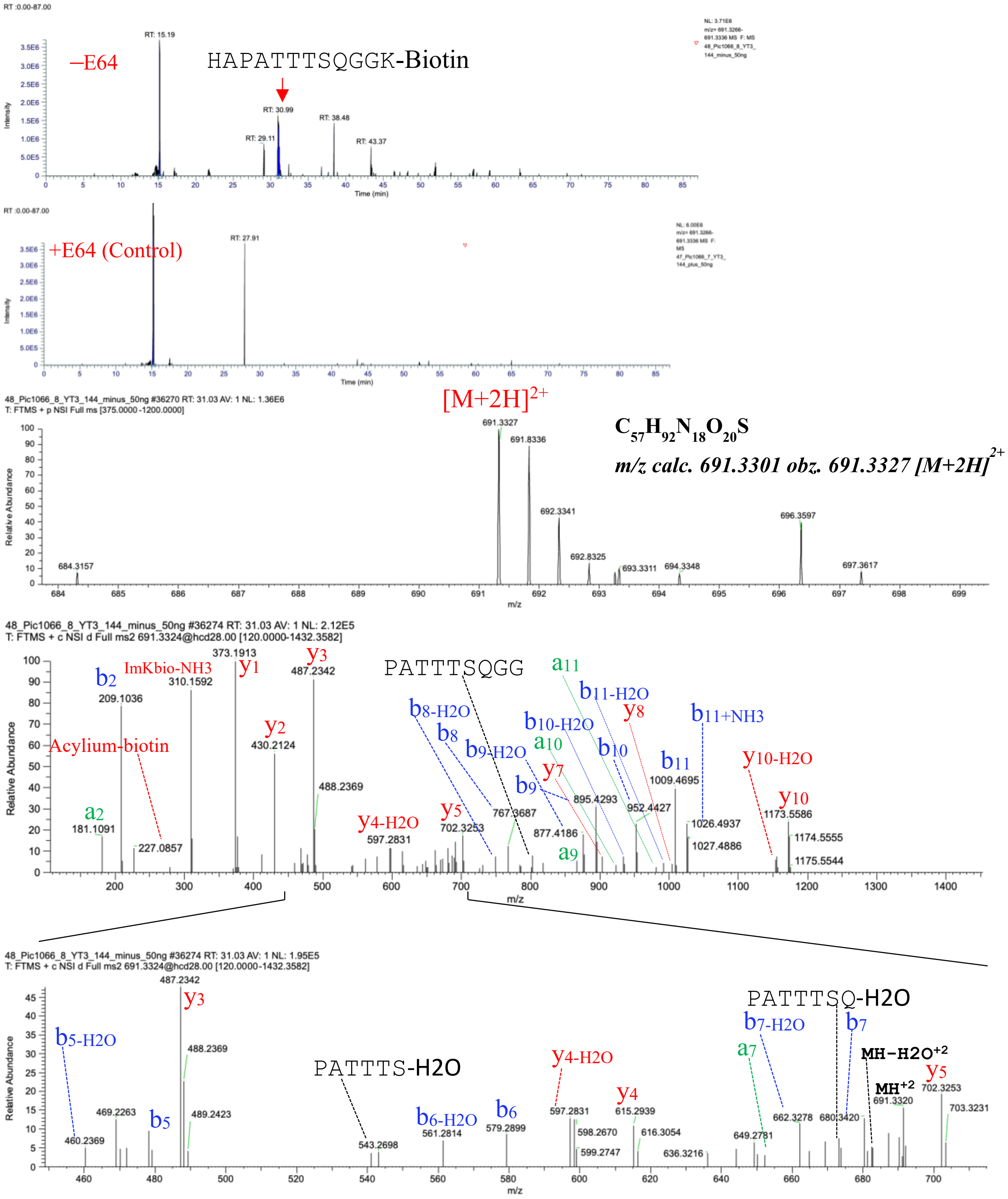

**Fig. S61.** HAPATTTSQGGK(Biotin) characterized by LC-MS/MS as a product of hCatS-catalyzed transpeptidation reaction. The figure displays a graphical representation of the extracted ion chromatogram (EIC), mass spectrum, and tandem mass spectrum of HAPATTTSQGGK(Biotin), identified from a reaction between TTSQGGK(Biotin) and Spike of SARS-CoV-2 in the presence of active hCatS. LC-MS/MS analysis was performed on samples treated with active hCatS (–E64) and compared to a control in which the enzyme was inhibited by E64 (+E64). The molecular ion was extracted based on its exact monoisotopic mass, and major fragment ions in the tandem mass spectrum were assigned

AGTITYTPKT

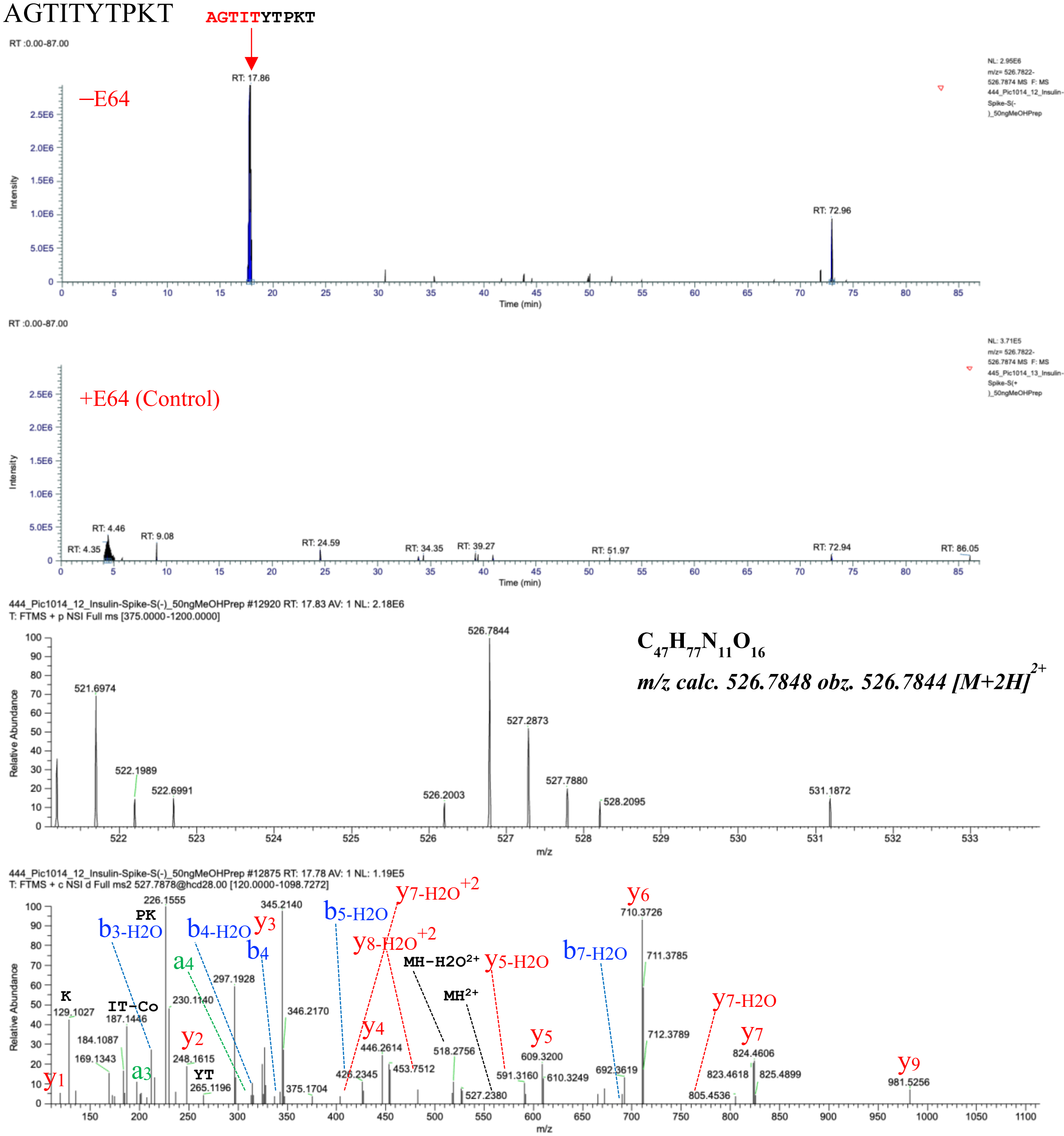

**Fig. S62.** **AGTITYTPKT** characterized by LC-MS/MS as a product of hCatS-catalyzed transpeptidation reaction. The figure displays a graphical representation of the extracted ion chromatogram (EIC), mass spectrum, and tandem mass spectrum of **AGTITYTPKT**, identified from a reaction between **Human proinsulin** and Spike of SARS-CoV-2 in the presence of active hCatS. LC-MS/MS analysis was performed on samples treated with active hCatS (–E64) and compared to a control in which the enzyme was inhibited by E64 (+E64). The molecular ion was extracted based on its exact monoisotopic mass, and major fragment ions in the tandem mass spectrum were assigned.

FQPTNGVGLIATLK

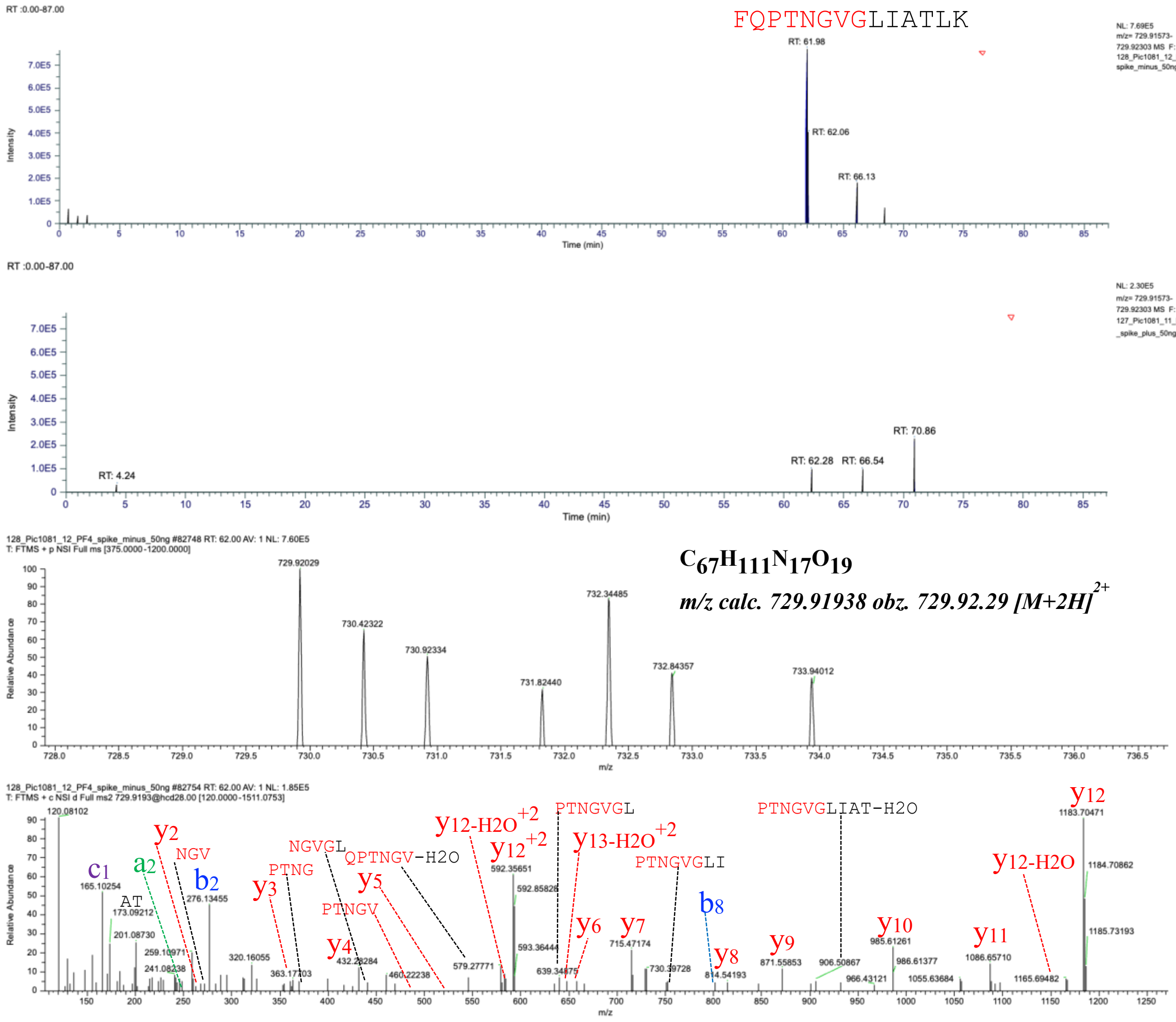

**Fig. S63.** FQPTNGVGLIATLK characterized by LC-MS/MS as a product of hCatS-catalyzed transpeptidation reaction. The figure displays a graphical representation of the extracted ion chromatogram (EIC), mass spectrum, and tandem mass spectrum of FQPTNGVGLIATLK, identified from a reaction between **Human proinsulin** and Spike of SARS-COV-2 in the presence of active hCatS. LC-MS/MS analysis was performed on samples treated with active hCatS (–E64) and compared to a control in which the enzyme was inhibited by E64 (+E64). The molecular ion was extracted based on its exact monoisotopic mass, and major fragment ions in the tandem mass spectrum were assigned.

QCCTYTPKT

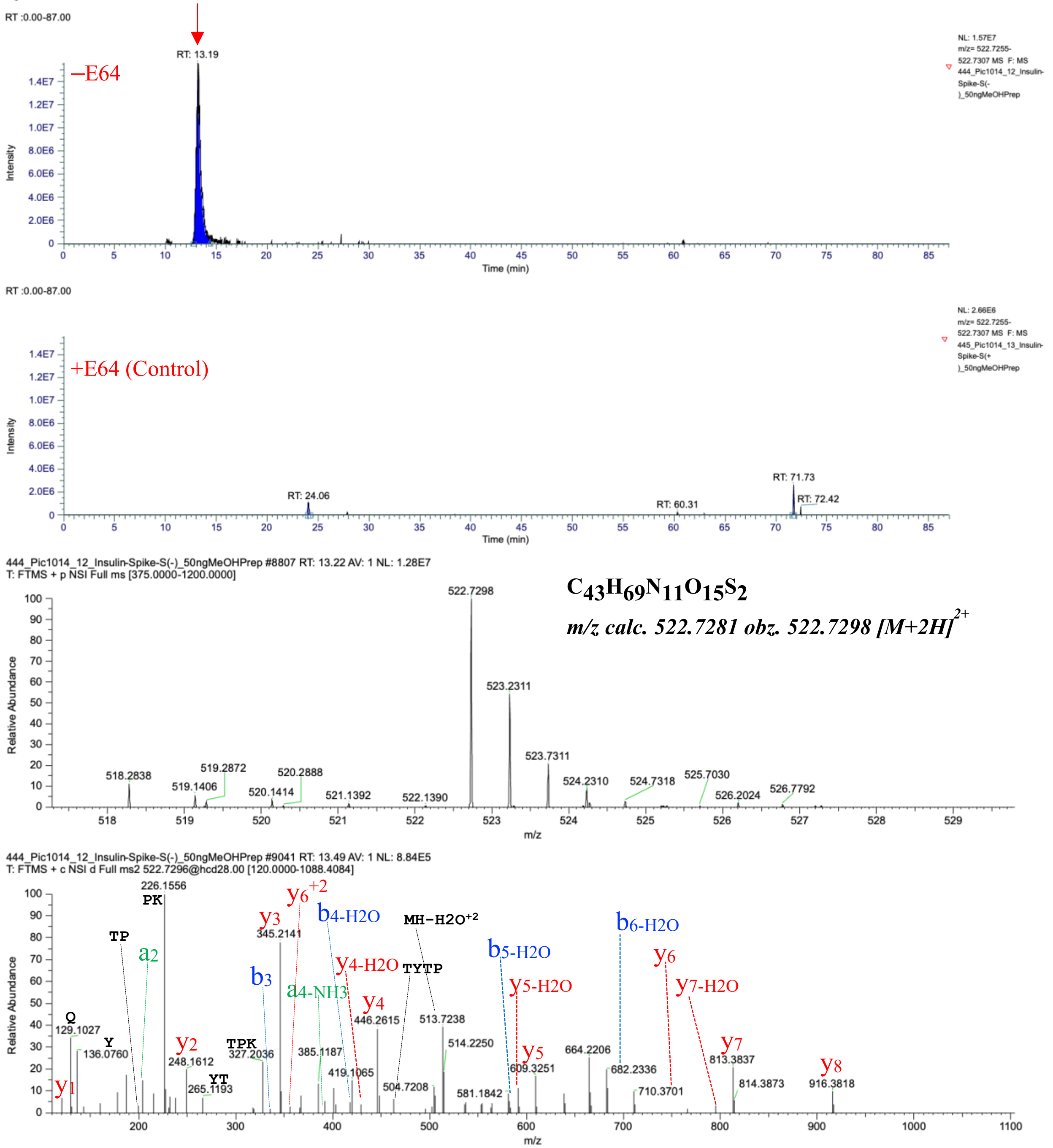

**Fig. S64.** **QCCTYTPKT** characterized by LC-MS/MS as a product of hCatS-catalyzed transpeptidation reaction. The figure displays a graphical representation of the extracted ion chromatogram (EIC), mass spectrum, and tandem mass spectrum of **QCCTYTPKT**, identified from a reaction of **Human proinsulin** in the presence of active hCatS. LC-MS/MS analysis was performed on samples treated with active hCatS (–E64) and compared to a control in which the enzyme was inhibited by E64 (+E64). The molecular ion was extracted based on its exact monoisotopic mass, and major fragment ions in the tandem mass spectrum were assigned.

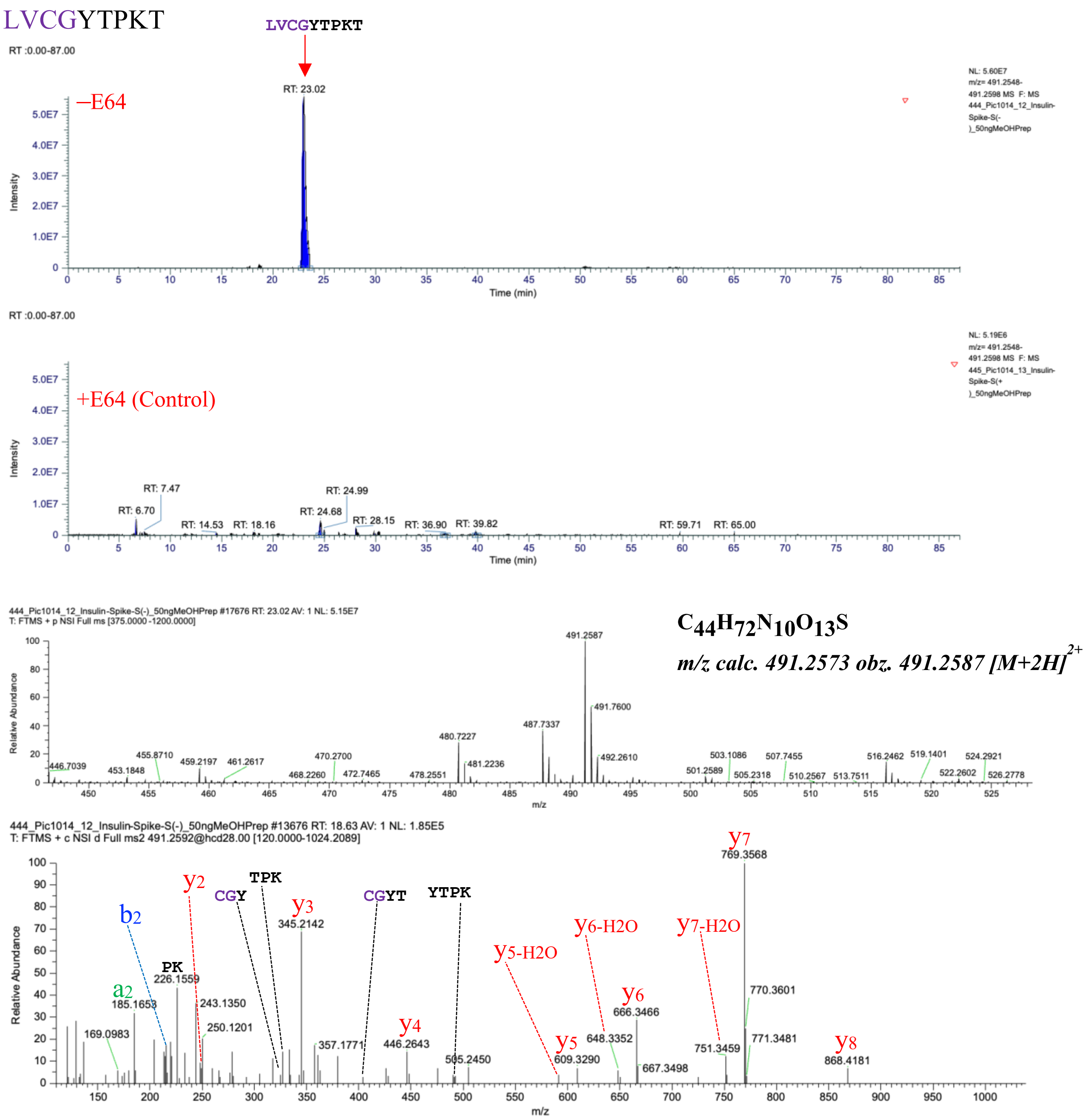

**Fig. S65.** **LVCGYTPKT** characterized by LC-MS/MS as a product of hCatS-catalyzed transpeptidation reaction. The figure displays a graphical representation of the extracted ion chromatogram (EIC), mass spectrum, and tandem mass spectrum of **LVCGYTPKT**, identified from a reaction of **Human proinsulin** in the presence of active hCatS. LC-MS/MS analysis was performed on samples treated with active hCatS (–E64) and compared to a control in which the enzyme was inhibited by E64 (+E64). The molecular ion was extracted based on its exact monoisotopic mass, and major fragment ions in the tandem mass spectrum were assigned.

SHLVETPKT

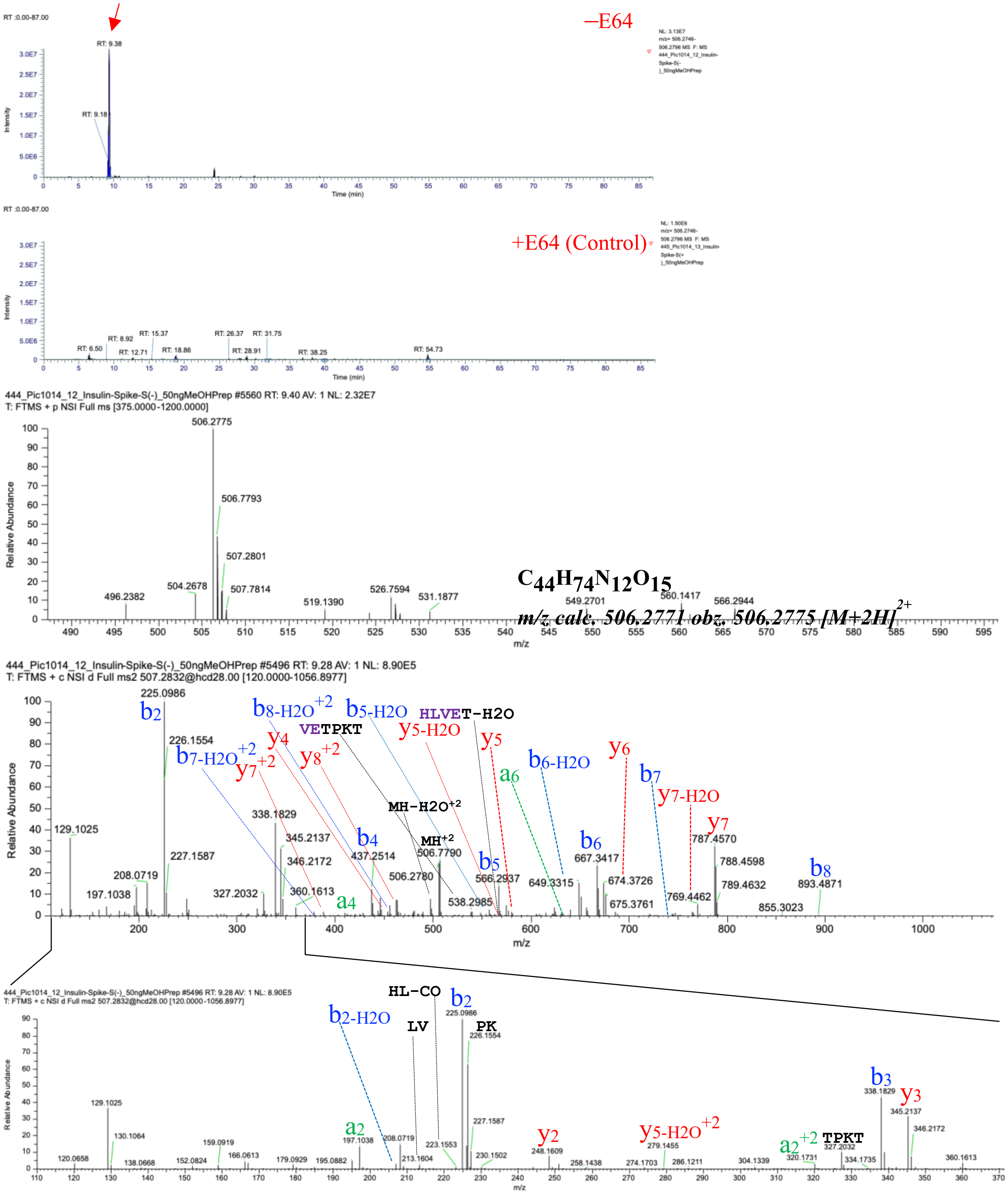

**Fig. S66.** **SHLVETPKT** characterized by LC-MS/MS as a product of hCatS-catalyzed transpeptidation reaction. The figure displays a graphical representation of the extracted ion chromatogram (EIC), mass spectrum, and tandem mass spectrum of **SHLVETPKT**, identified from a reaction of **Human proinsulin** in the presence of active hCatS. LC-MS/MS analysis was performed on samples treated with active hCatS (–E64) and compared to a control in which the enzyme was inhibited by E64 (+E64). The molecular ion was extracted based on its exact monoisotopic mass, and major fragment ions in the tandem mass spectrum were assigned.

LVCGFVNQHLC

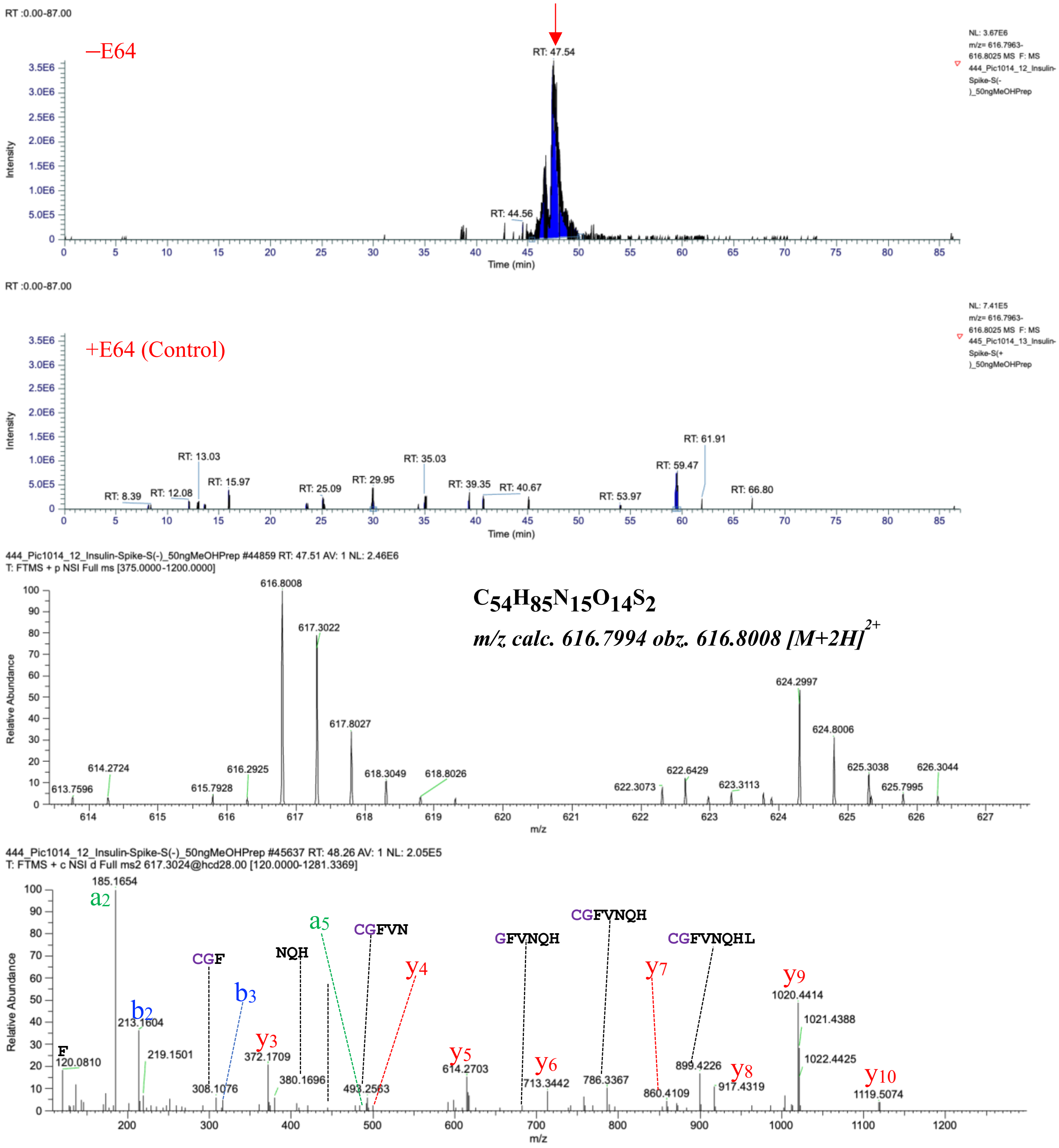

**Fig. S67.** **LVCGFVNQHLC** characterized by LC-MS/MS as a product of hCatS-catalyzed transpeptidation reaction. The figure displays a graphical representation of the extracted ion chromatogram (EIC), mass spectrum, and tandem mass spectrum of **LVCGFVNQHLC**, identified from a reaction of **Human proinsulin** in the presence of active hCatS. LC-MS/MS analysis was performed on samples treated with active hCatS (–E64) and compared to a control in which the enzyme was inhibited by E64 (+E64). The molecular ion was extracted based on its exact monoisotopic mass, and major fragment ions in the tandem mass spectrum were assigned.

### SLYQTPKT

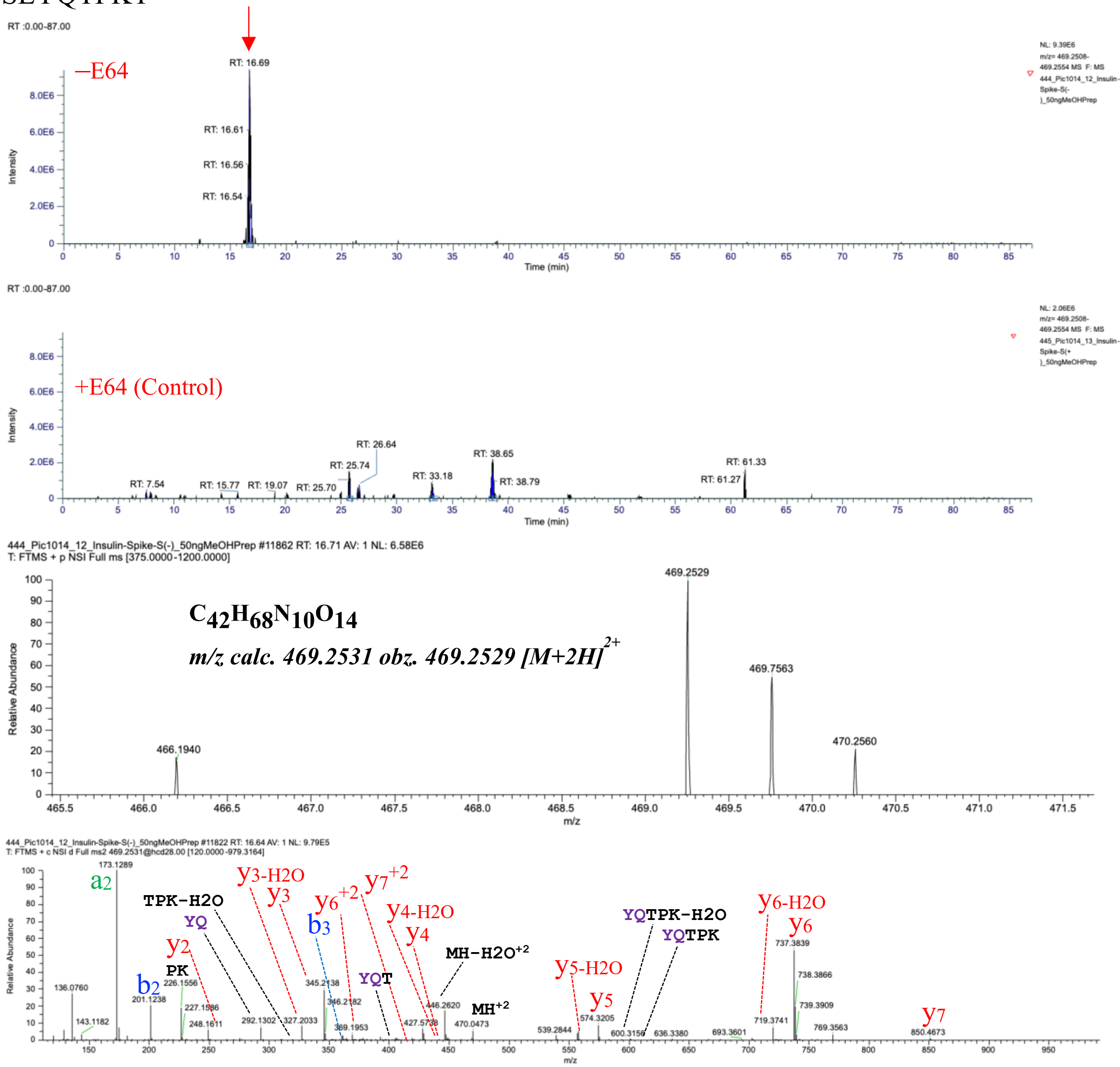

**Fig. S68.** **SLYQTPKT** characterized by LC-MS/MS as a product of hCatS-catalyzed transpeptidation reaction. The figure displays a graphical representation of the extracted ion chromatogram (EIC), mass spectrum, and tandem mass spectrum of **SLYQTPKT**, identified from a reaction of **Human proinsulin** in the presence of active hCatS. LC-MS/MS analysis was performed on samples treated with active hCatS (–E64) and compared to a control in which the enzyme was inhibited by E64 (+E64). The molecular ion was extracted based on its exact monoisotopic mass, and major fragment ions in the tandem mass spectrum were assigned.

GIVEQCCTTPKT

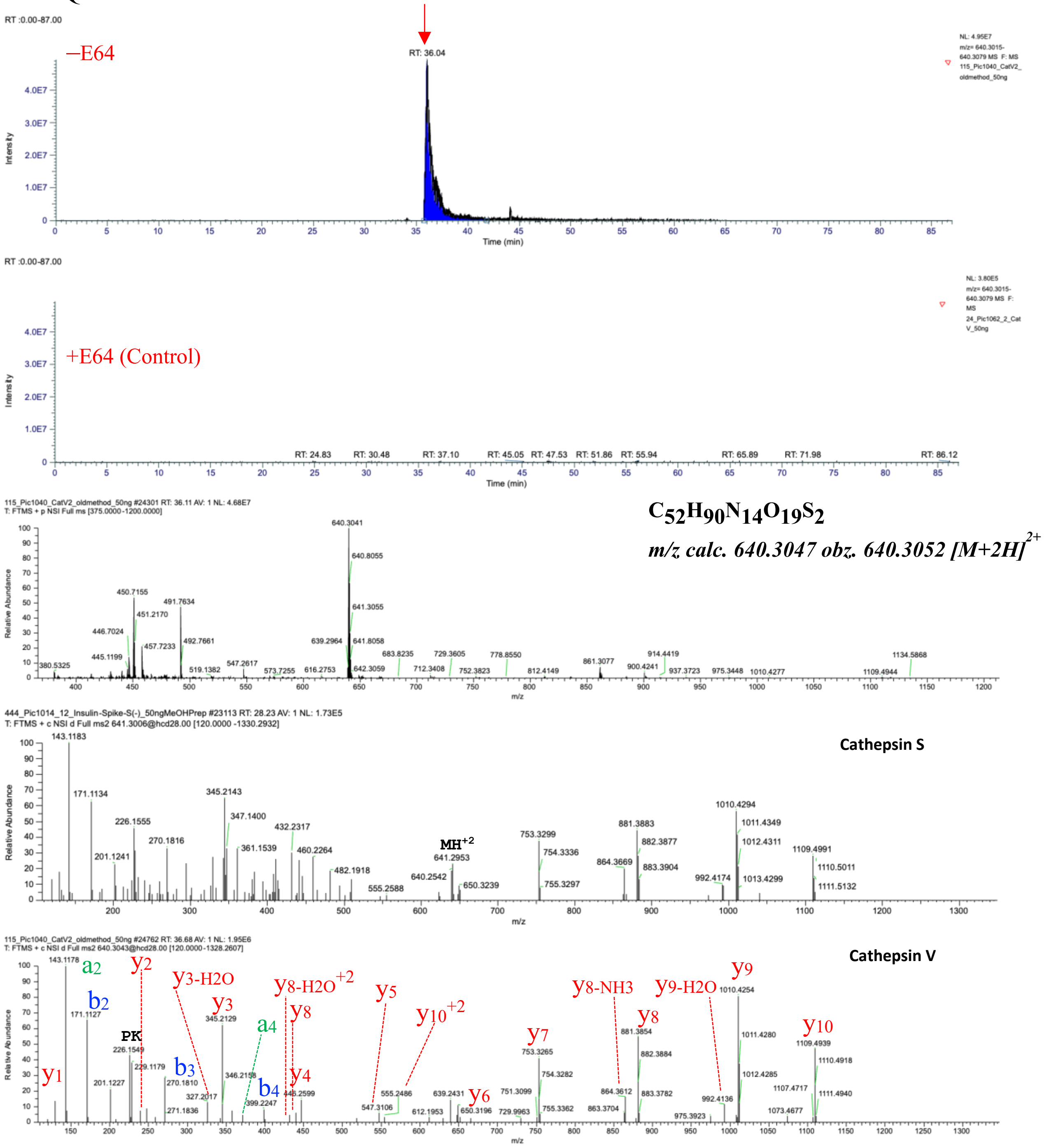

**Fig. S69.** **GIVEQCCTTPKT** characterized by LC-MS/MS as a product of hCatS-catalyzed transpeptidation reaction. The figure displays a graphical representation of the extracted ion chromatogram (EIC), mass spectrum, and tandem mass spectrum of **GIVEQCCTTPKT**, identified from a reaction of **Human proinsulin** in the presence of active hCatS. LC-MS/MS analysis was performed on samples treated with active hCatS (–E64) and compared to a control in which the enzyme was inhibited by E64 (+E64). The molecular ion was extracted based on its exact monoisotopic mass, and major fragment ions in the tandem mass spectrum were assigned.

GIVEQCCTYTPKT

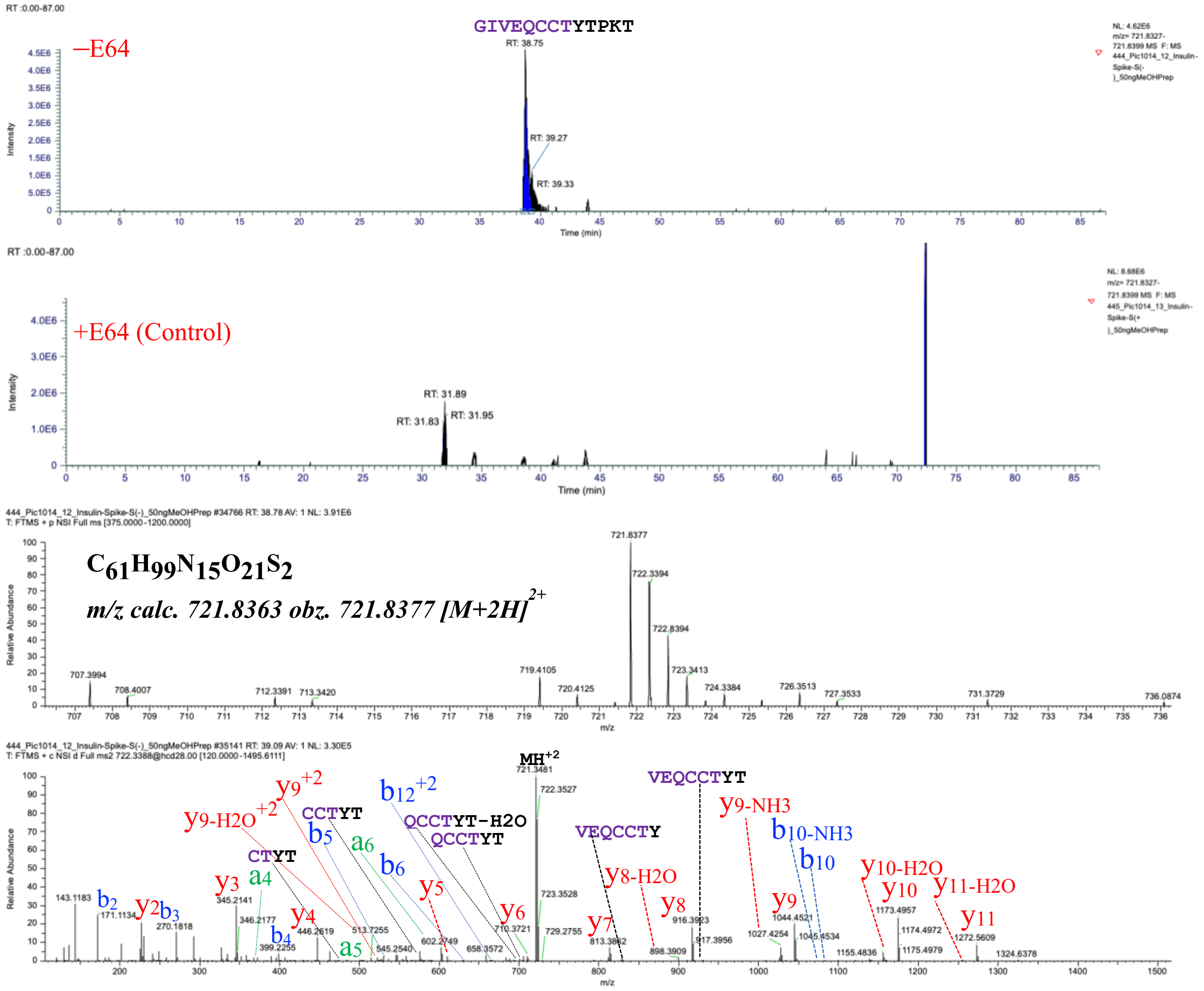

**Fig. S70.** **GIVEQCCTYTPKT** characterized by LC-MS/MS as a product of hCatS-catalyzed transpeptidation reaction. The figure displays a graphical representation of the extracted ion chromatogram (EIC), mass spectrum, and tandem mass spectrum of **GIVEQCCTYTPKT**, identified from a reaction of **Human proinsulin** in the presence of active hCatS. LC-MS/MS analysis was performed on samples treated with active hCatS (–E64) and compared to a control in which the enzyme was inhibited by E64 (+E64). The molecular ion was extracted based on its exact monoisotopic mass, and major fragment ions in the tandem mass spectrum were assigned.

### GIVEQCCTGSHLVE

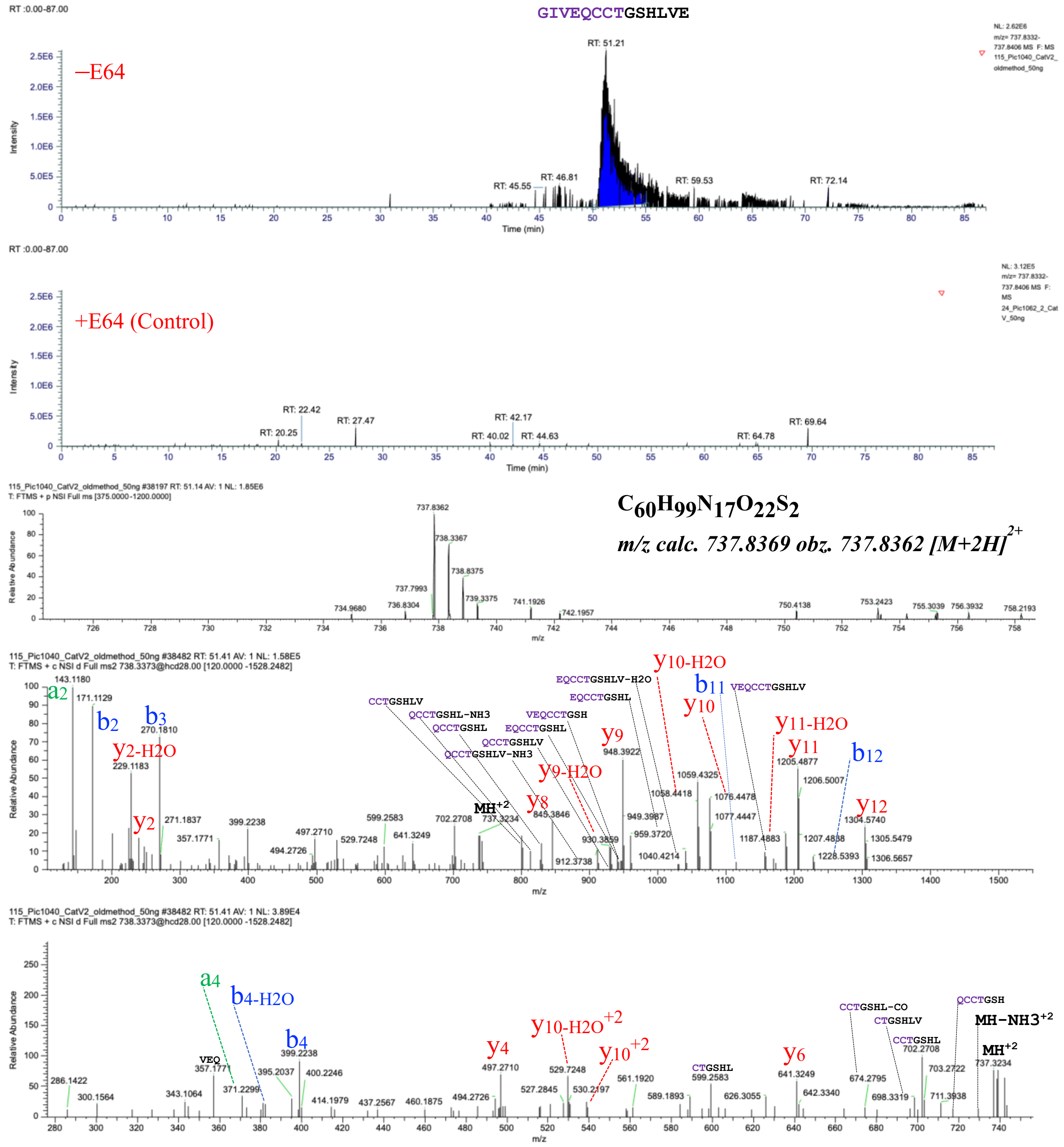

**Fig. S71.** **GIVEQCCTGSHLVE** characterized by LC-MS/MS as a product of hCatS-catalyzed transpeptidation reaction. The figure displays a graphical representation of the extracted ion chromatogram (EIC), mass spectrum, and tandem mass spectrum of **GIVEQCCTGSHLVE**, identified from a reaction of **Human proinsulin** in the presence of active hCatS. LC-MS/MS analysis was performed on samples treated with active hCatS (–E64) and compared to a control in which the enzyme was inhibited by E64 (+E64). The molecular ion was extracted based on its exact monoisotopic mass, and major fragment ions in the tandem mass spectrum were assigned.

FVNQHLC

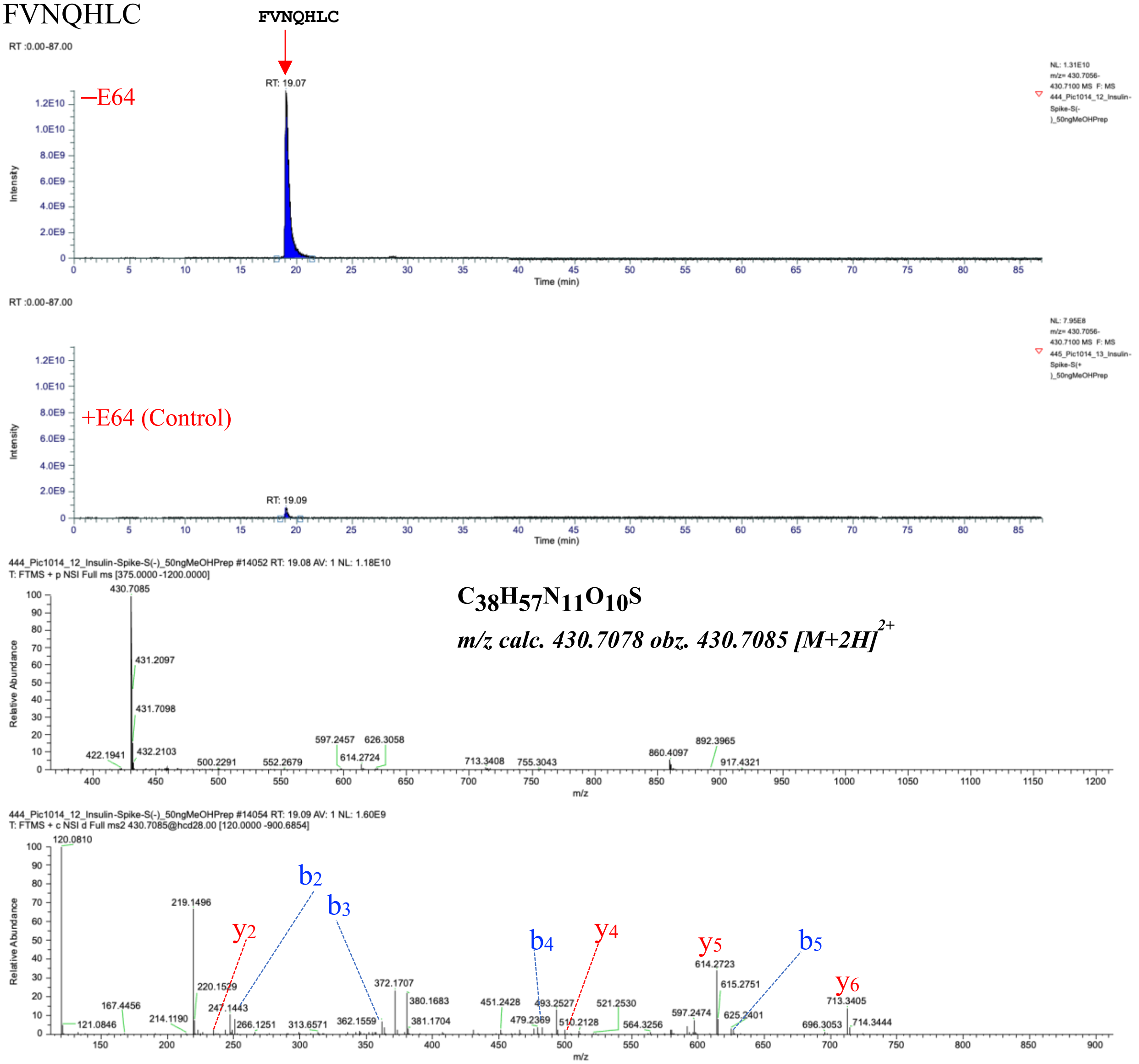

**Fig. S72. FVNQHLC** characterized by LC-MS/MS as a product of hCatS-catalyzed reaction. The figure displays a graphical representation of the extracted ion chromatogram (EIC), mass spectrum, and tandem mass spectrum of **FVNQHLC**, identified from a reaction of **Human proinsulin** in the presence of active hCatS. LC-MS/MS analysis was performed on samples treated with active hCatS (–E64) and compared to a control in which the enzyme was inhibited by E64 (+E64). The molecular ion was extracted based on its exact monoisotopic mass, and major fragment ions in the tandem mass spectrum were assigned.

DEMIAQYTFVNQHLC

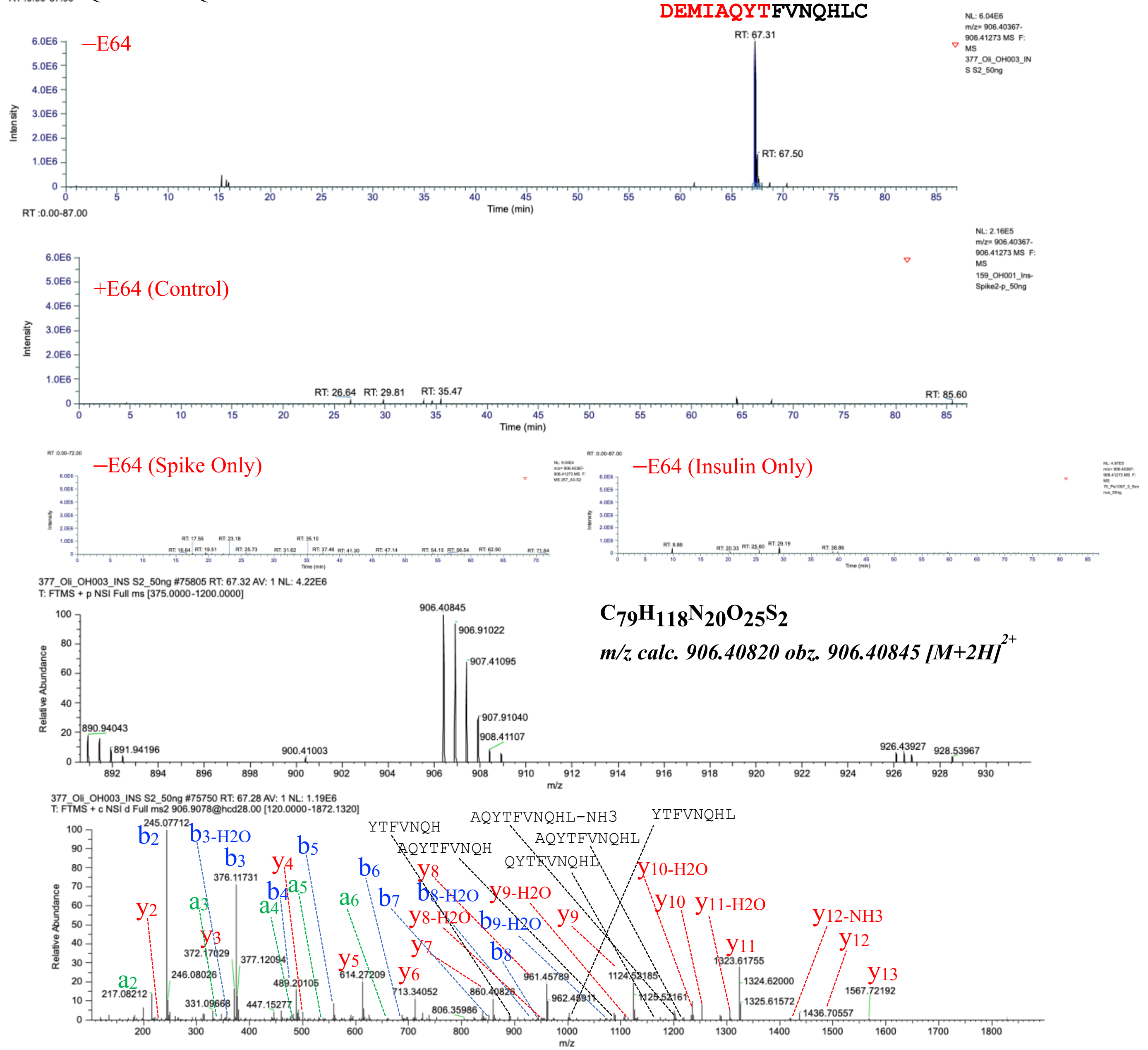

**Fig. S73.** **DEMIAQYTFVNQHLC** characterized by LC-MS/MS as a product of hCatS-catalyzed transpeptidation reaction. The figure displays a graphical representation of the extracted ion chromatogram (EIC), mass spectrum, and tandem mass spectrum of **DEMIAQYTFVNQHLC**, identified from a reaction of **Human proinsulin and Spike of SARS-CoV-2** in the presence of active hCatS. LC-MS/MS analysis was performed on samples treated with active hCatS (–E64) and compared to a control in which the enzyme was inhibited by E64 (+E64). –E64 (Spike only) control experiment has **Spike of SARS-CoV-2** in the presence of active hCatS. –E64 (Insulin only) control experiment has **human proinsulin** in the presence of active hCatS. The molecular ion was extracted based on its exact monoisotopic mass, and major fragment ions in the tandem mass spectrum were assigned.

2nd generation cis/transpeptides derived from Insulin

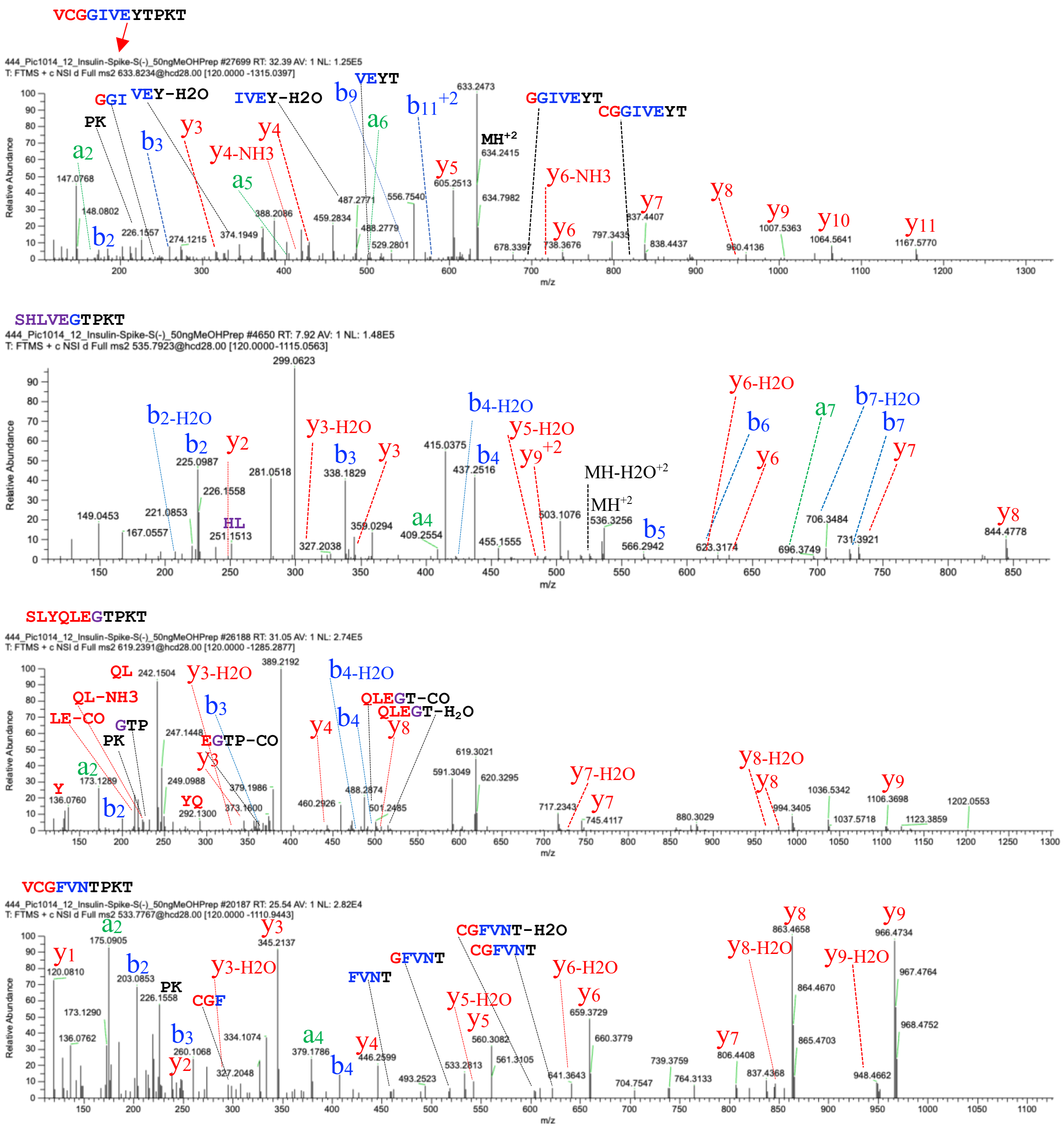

**Fig. S74.** Higher order generation of cis/transpeptides characterized by LC-MS/MS as a product of hCatS-catalyzed reaction. The figure displays a graphical representation of the extracted ion chromatogram (EIC), mass spectrum, and tandem mass spectrum of cis/transpeptides, identified from a reaction of Human proinsulin/Spike in the presence of active hCatS. LC-MS/MS analysis was performed on samples treated with active hCatS (–E64) and compared to a control in which the enzyme was inhibited by E64 (+E64). The molecular ion was extracted based on its exact monoisotopic mass, and major fragment ions in the tandem mass spectrum were assigned.

SLEVIKLIATLK

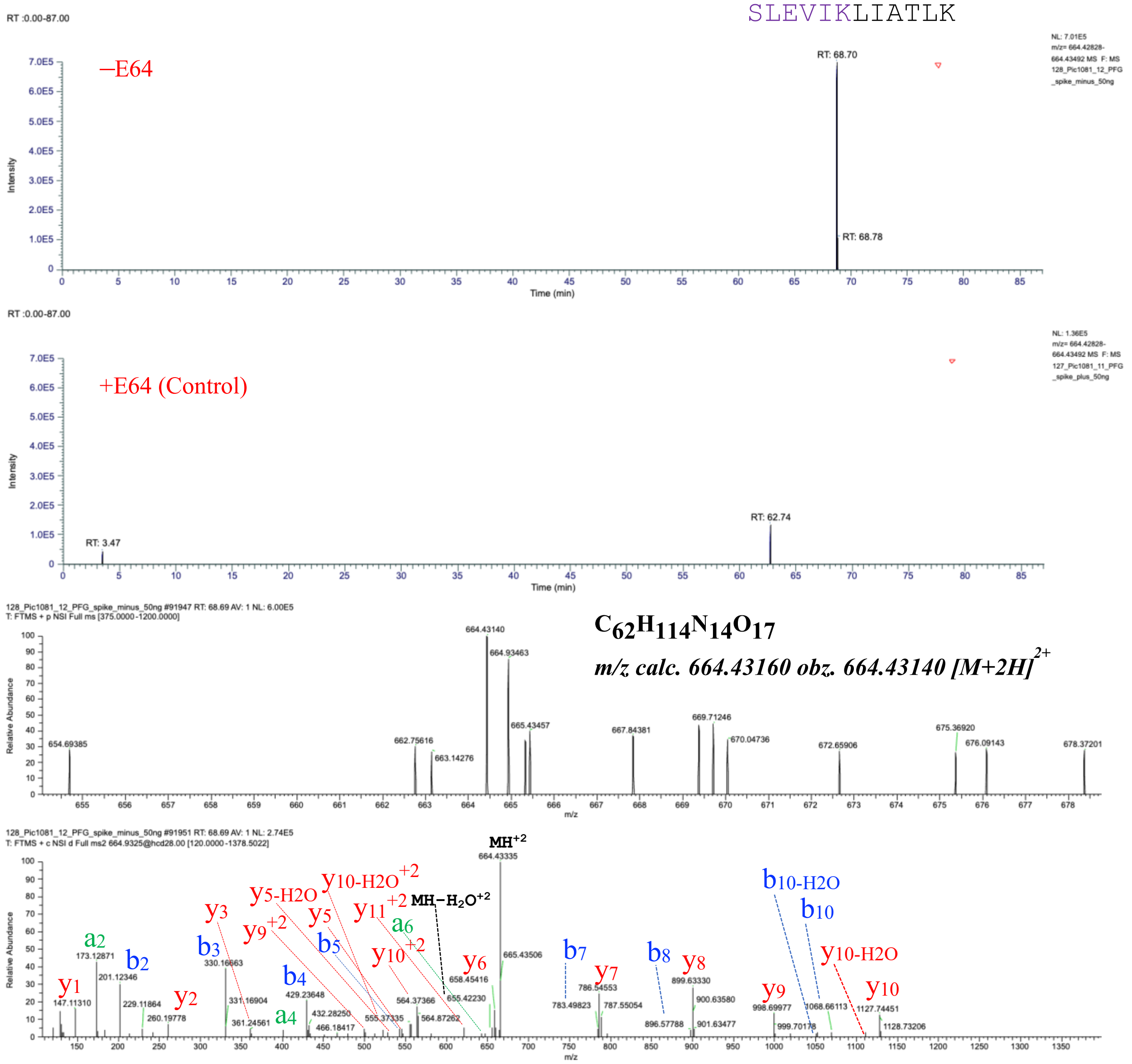

**Fig. S75.** SLEVIKLIATLK characterized by LC-MS/MS as a product of hCatS-catalyzed transpeptidation reaction. The figure displays a graphical representation of the extracted ion chromatogram (EIC), mass spectrum, and tandem mass spectrum of SLEVIKLIATLK, identified from a reaction of **Human PF4** in the presence of active hCatS. LC-MS/MS analysis was performed on samples treated with active hCatS (–E64) and compared to a control in which the enzyme was inhibited by E64 (+E64). The molecular ion was extracted based on its exact monoisotopic mass, and major fragment ions in the tandem mass spectrum were assigned.

Derivatization of TTSQGGK(N3)YGRKKRRQRRRK-5FM

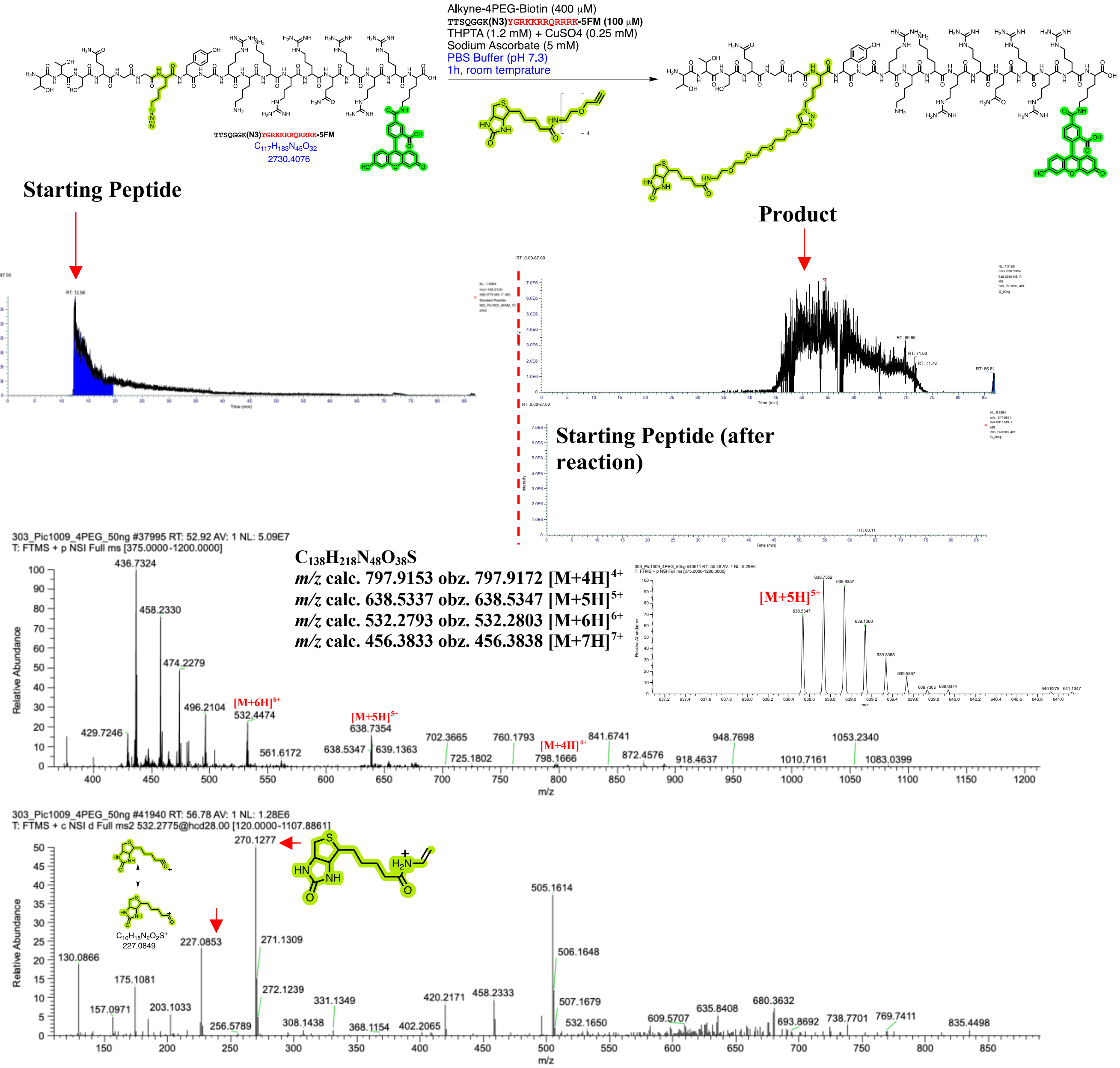

**Fig S76.** Derivatization of TTSQGGK(N3)YGRKKRRQRRRK-5FM by Alkyne-4PEG-Biotin using azide-alkyne click reaction. Generation of strong marker ions at 270.1277 and 277.0853 corresponds to 4PEG-Biotin tag

#### Uptake of TAT-peptide by RAW cells

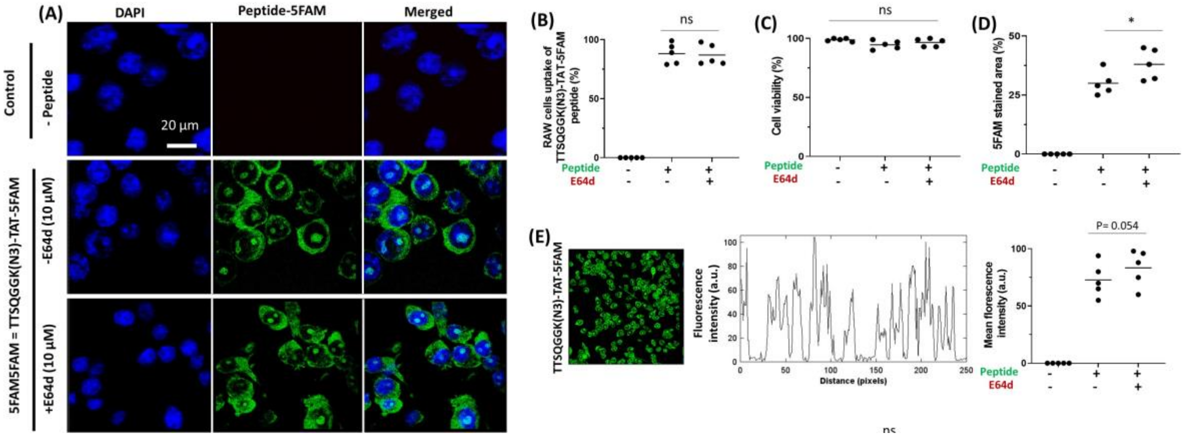

**Figure S77:** (A) Uptake of the TAT-peptide by RAW cells. (B–E) Quantification of TAT-peptide internalization in RAW cells.

GGK(**4PEG-Biotin**)*GLVEGLMTTVHALVHHR* isolated

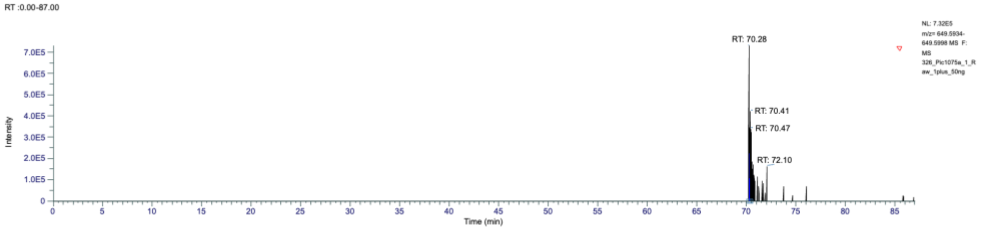

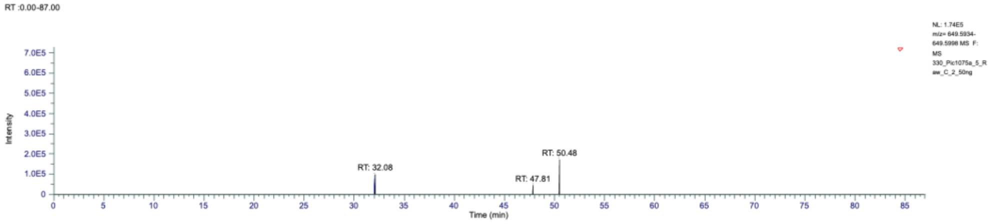

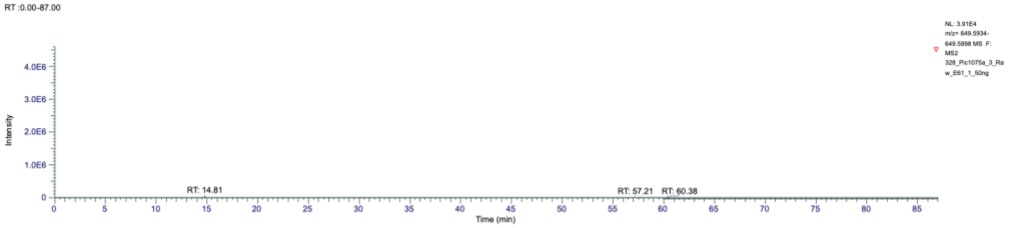

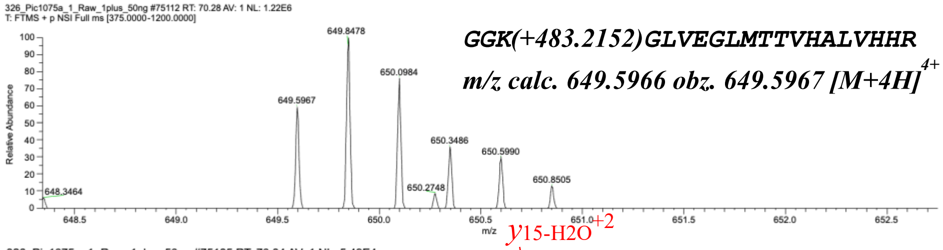

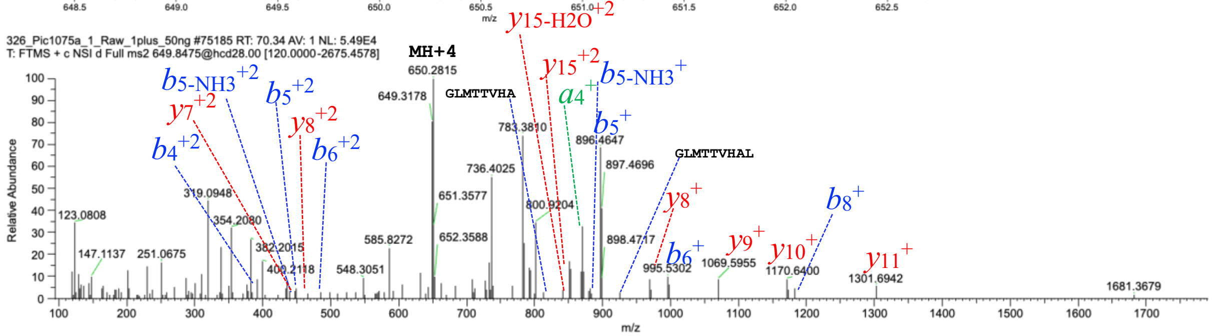

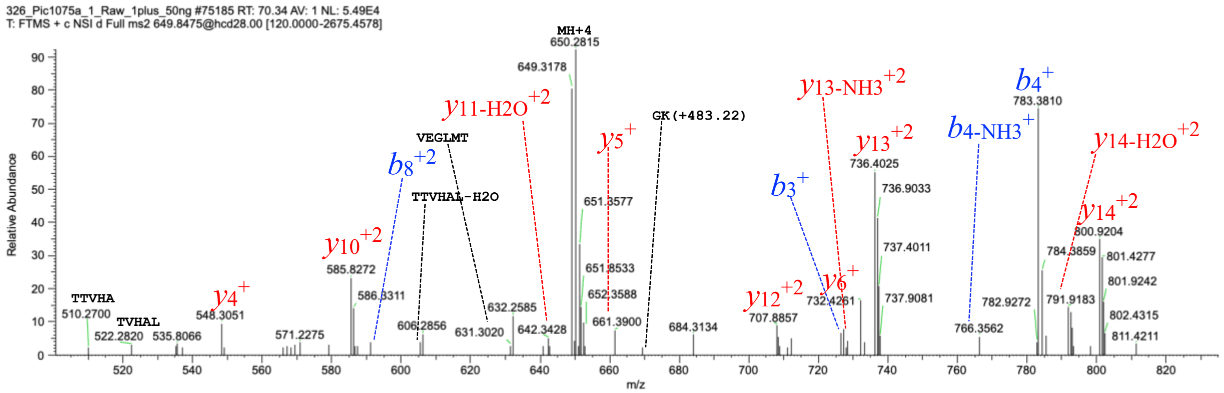

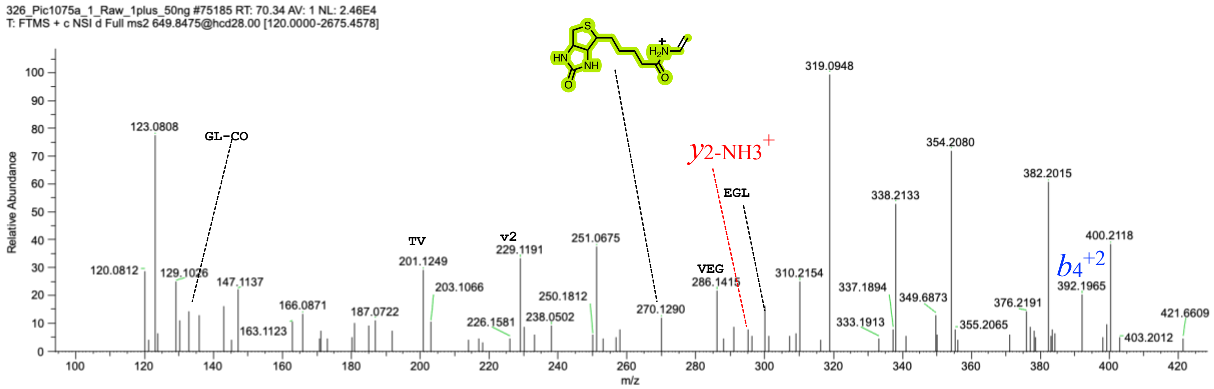

**Fig S78:** Mass spectrum and tandem mass spectrum of the peptide GGK(**4PEG-Biotin**)*GLVEGLMTTVHALVHHR* isolated from *Mus musculus* RAW cell line after treatment with TTSQGGK(N3)YGRKKRRQRRRK-5FM. Fragment ion peaks are annotated to confirm the sequence.

EAAGAGSARPEAPG(**4PEG-Biotin**)*WVRGDAR*

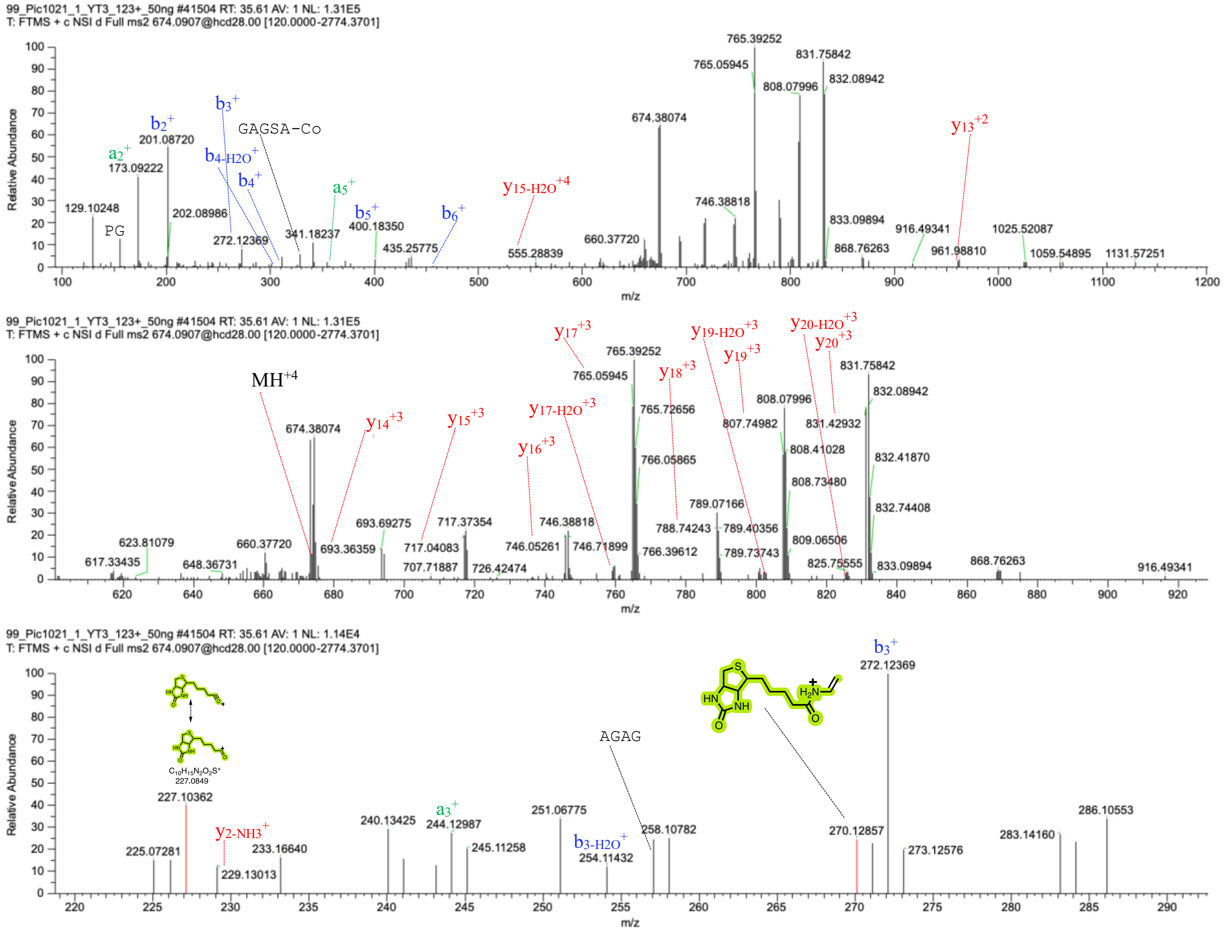

**Figure S79:** Mass spectrum and tandem mass spectrum of the peptide EAAGAGSARPEAPG(**4PEG-Biotin**)*WVRGDAR* isolated from *Mus musculus* RAW cell line after treatment with TTSQGGK(N3)YGRKKRRQRRRK-5FM. Fragment ion peaks are annotated to confirm the sequence
